# Unexplored regions of the protein sequence-structure map revealed at scale by a library of foldtuned language models

**DOI:** 10.1101/2023.12.22.573145

**Authors:** Arjuna M. Subramanian, Zachary A. Martinez, Alec L. Lourenço, Sonia C. Yuan, Shichen Liu, Ting-Yu Wang, Tsui-Fen Chou, Matt Thomson

## Abstract

Amino-acid sequence space is combinatorially vast, with well-folded proteins distributed sparsely and connected by vanishingly few permissible mutational paths. Novel-in-sequence versions of structures observed in nature promise to sample features such as new binding motifs and active site geometries but are rendered inaccessible to evolution or direct search by the extent of sequence perturbations required. Here we introduce a novel algorithm – termed “foldtuning” – that leverages principles of adversarial learning to drive protein language models (PLMs) to erase detectable homology to natural sequences while preserving a target structure, systematically traversing protein-space without being limited by evolutionary barriers. We build foldtuned PLMs for >700 targets including membrane-bound receptors, redox enzymes, and signaling domains. Foldtuned proteins are diverse and far-from-natural in sequence, filling out structurally-equivalent families defined by fundamental biophysical constraints invisible to traditional sequence-based bioinformatics methods. Experimental characterization demonstrates that foldtuned proteins express stably *in vitro* and function *in vivo*. By revealing sequence-structure information at scale beyond evolution, foldtuning promises to accelerate the reconstitution and realization of novel-to-nature systems for synthetic biology problems from therapeutics to catalysis.

## INTRODUCTION

Nature has likely sampled only a fraction of all protein sequences and structures allowed by the laws of biophysics.^1^ The 20 proteinogenic amino acids ensure a combinatorially vast sequence-space; to roughly comprehend this magnitude, consider that making one copy of each of the ∼ 10^78^ possible sequences for a small protein domain of length 60 would require more matter than exists in the visible universe. High-quality sequence databases, in contrast, contain ∼ 10^9^ unique protein sequences distributed across the tree of life.^2,3^ The observed protein catalog likely reflects selection for factors such as favorable folding kinetics, cofactor usage, and binding/catalytic functions.^4–8^ However, these proteins, no matter how evolutionarily fit, are not the only solutions of the sequence-to-structure mapping problem. Hydrophobic/polar patterning schemes distinguish energetically-favorable three-dimensional structures and generate stable *α*-helical bundle proteins encoded by novel sequences.^9–16^ Deep multiple sequence alignments (MSAs) capture sparse co-evolutionary signals sufficient to generate artificial proteins with comparable stability to natural examples.^17,18^ And measurements on random sequence libraries suggest that as many as 1-in-10^11^ amino-acid sequences may code for functional proteins, providing ample “sparks” for alternate protein populations beyond nature.^19,20^ Systematically locating stable, functional proteins that reconstitute known structural motifs but lie in regions of sequence-space with no meaningful similarity to nature promises to unlock expanded repertoires of binding partners, signaling interactions, and substrate scopes for synthetic biology, while revealing key amino-acid sequence rules and constraints undergirding the fundamental biophysics of molecular machines.

We posit that the problem of mining such “döppelganger” proteins can be met by a search strategy that balances large perturbations to sequence against small perturbations to backbone structure. Global sequence perturbations of this magnitude are not accessible to directed evolution – which searches sequence-space locally under strong stability and fitness restrictions – or to machine learning models trained on high-throughput but inescapably local fitness data collected in deep-mutational scanning (DMS) experiments.^21–23^ Inverse-folding structure-to-sequence design methods can diversify sequence more substantially, but enforce strict back-bone constraints that preclude the sorts of small structural innovations and ornamentations that have conferred new and/or expanded functionalities throughout natural evolution.^24–27^ In contrast, protein language models (PLMs) explicitly learn sequence-level amino-acid dependencies, implicitly internalizing the information flow from sequence to structure.^28–30^ Furthermore, when used as protein *generators*, PLMs are known to reach beyond natural sequences and structures.^29,31–34^

Given that PLMs understand the core determinants of sequence-to-structure mapping while retaining an innate explorative capacity, we introduce “foldtuning” as an approach that transforms PLMs into probes that trace structure-preserving paths through far-from-natural regions of protein sequence-space. Drawing conceptual inspiration from the counterfeiting games played by generative adversarial networks (GANs), foldtuning leverages competition and complementation between PLMs and structure prediction models to mine protein-space for examples that honor the “grammar” of a target backbone while exhibiting novel sequence semantics.^28,31,35–40^ We successfully apply foldtuning to > 700 structural motifs of interest from the SCOP and InterPro databases, covering all four major tertiary structure topology classes (all-*α*, all-*β*, *α* + *β*, and *α*/*β*) and wide-ranging functional families, including GPCRs, transcription factors, cell-to-cell signaling domains, and P450 enzymes. We show that, generally, successive rounds of foldtuning progressively reduce similarity between PLM-generated and wild-type sequences, reaching new-to-nature ‘rules of language’ for constructing proteins – novel strategies for accommodating the biophysical constraints on structure that hold beyond evolution. High-throughput screening of foldtuned sequence libraries for three target folds – the SH3 adapter domain, barstar, and insulin – identifies stable, functional candidate variants with 0-40% sequence identity to their closest respective neighbors in the known protein universe. Ultimately, sequence remodeling through foldtuning reflects a “novelty first” ethos that stretches the limits of how far common protein folds can be diversified at the sequence level while extracting and preserving critical minimal rules of structure and function.

## RESULTS

### Sequence exploration with ‘soft’ structure constraints

In order to robustly access far-from-natural sequences coding for many structurally diverse fold classes we develop “foldtuning,” a structure-oriented algorithm that drives a PLM to sample extreme sequence novelty while holding to a target fold class, summarized in Figure 1A. The PLM of choice is first finetuned on natural protein fragments (sourced from a custom SCOP-UniRef50 database or InterPro PDB-derived metadata depending on the target, as described further in Methods)^41^ that adopt the target backbone structure of interest; this initial step is analogous to “evotuning” on a functional family as has been applied previously for PLM-based enzyme design.^32,42^ Following this extra fold-specific pretraining, foldtuning proceeds through alternating rounds of: (1) sequence generation out of the current model state, and (2) model update by finetuning on a subset of self-generated artificial sequences that are predicted to coarsely adopt the target fold while differing maximally from natural counterparts in terms of sequence (Figure 1B). Selection for preserving the target fold is achieved by predicting each structure with ESMFold and assigning a SCOP or InterPro label with Foldseek-TMalign search; this is a “soft” structural constraint, using a TMscore > 0.5 global alignment threshold to place generated candidates within the target fold *family* or *distribution* (Figure 1C).^28,38,43,44^. Selection for sequence dissimilarity is enforced by ranking all structurally-validated sequences by semantic change – defined for a generated sequence 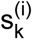 as the smallest L_1_-distance between the ESM2-650M embeddings of 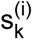 and any of the natural training sequences – in decreasing order, and taking the top 100 as the next synthetic training data for model updating. Dimension-reduced views of these embeddings for a representative subset of target folds suggest that ESM2-650M captures – and foldtuning navigates along – a representation of the sequence→structure map where structural classes (grouping corresponding pairs of natural and foldtuned artificial sequences) largely separate from one another, with artificial sequences drifting from their natural parents along concerted trajectories in the embedding-space (Figure 1D, Figures S1–S2). Each foldtuning cycle is consequently analogous to a step along a path that drives a PLM to access subpopulations of progressively further-from-natural artificial sequences while preserving the broad form of the fixed target structure.

**Figure 1.**
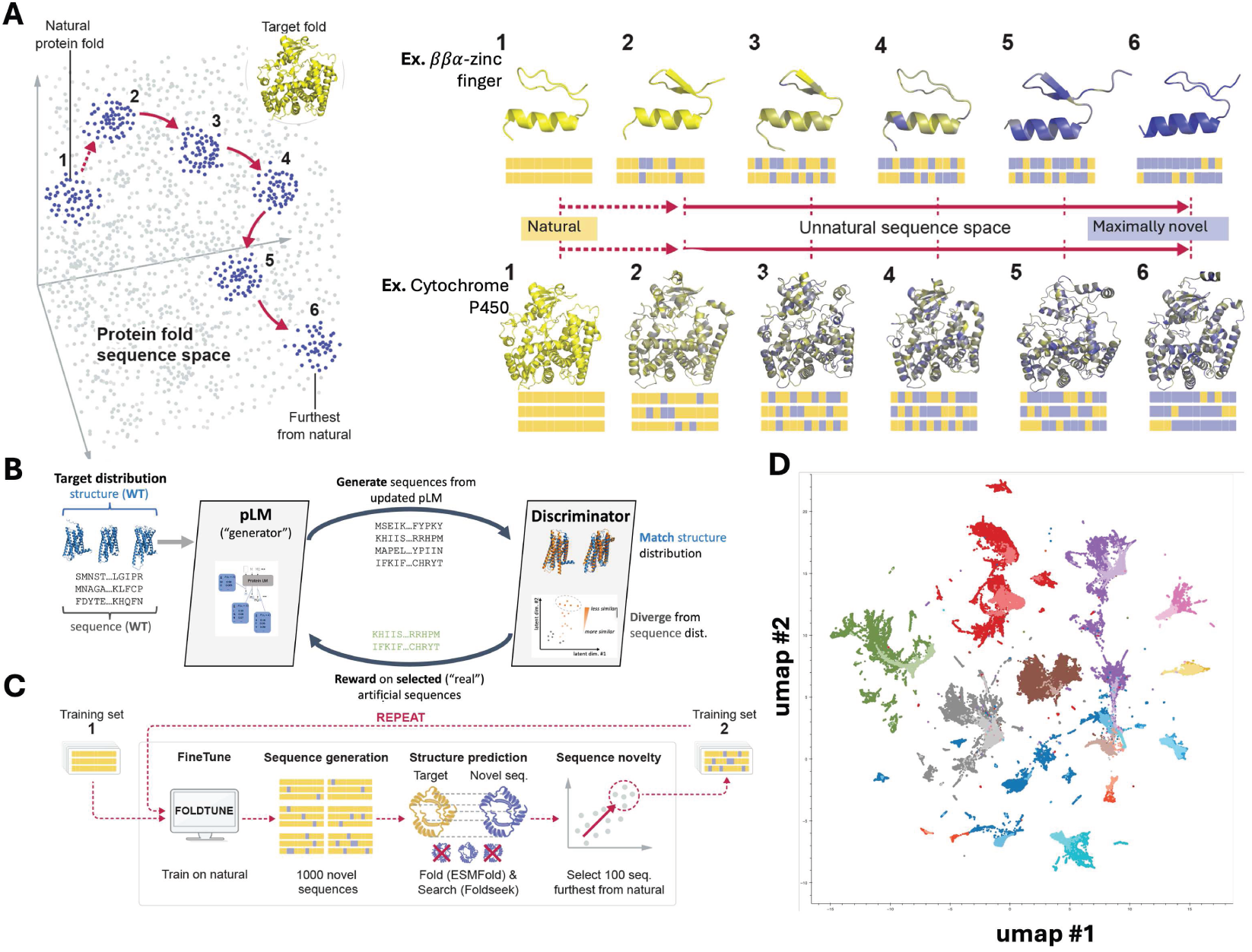
Foldtuning explores far-from-natural sequences encoding alternate versions of natural protein structures. (A) Foldtuning uses a protein language model (PLM)-based strategy to probe “outwards” in sequence-space, detecting subpopulations of sequences with progressively decreasing similarity to natural sequences while preserving a given target backbone structure. Examples: (top) *ββα*-zinc finger, (bottom) cytochrome P450. (B) Foldtuning alternates between sequence generation and discrimination/selection rounds in a closed-loop, inspired by the architecture of generative adversarial networks (GANs). (C) For a provided backbone target fold, a PLM (ProtGPT2) is initially finetuned (**1**) on target fold examples harvested from deep mining of natural sequence/structure data; in subsequent rounds the PLM is finetuned on self-generated artificial sequences validated by structure prediction (ESMFold) and structure-based search (Foldseek) and selected for maximization of semantic change w.r.t. natural examples (**2**). (D) 2D UMAP representation of ESM2-650M embeddings of natural (dark) and foldtuned (light) sequence examples for eleven representative target fold classes.

Using ProtGPT2 as the base pretrained PLM, we foldtuned models for 727 structural targets; 708 SCOP folds (out of the top 850 ranked by natural abundance, for an 83.3% success rate), plus 19 cytokines and chemokines of interest curated from InterPro. Successfully foldtuned SCOP targets span numerous classes of functional interest for synthetic biology applications, including transcription factor DNA-binding domains, GPCR/small GTPase signaling components, modular cell surface receptor domains, and defense proteins (e.g. antimicrobial peptides, toxins). Foldtuned versions of ProtGPT2 are effective at landing on the target backbone fold, increasing from a median *structural hit rate* of 0.203 after evotuning alone to 0.565 after two rounds of updates on far-from-natural artificial sequences, falling slightly to 0.509 after four rounds (Figure 2A). The total structural hit rate over all targets was 52.2%; overall and per-fold hit rates far exceed hit rates for control sequences derived from representative pair-wise alignments, indicating that foldtuning extracts meaningful structure-informing sequence features beyond evolutionary statistical correlations (Table S1).^30^ Sequence novelty relative to natural examples increases with additional update rounds; the median *sequence escape rate* – the fraction of target structure matches that do not feature any detectable sequence homology to any protein in UniRef50 – does not change significantly from evotuning (0.134) through two rounds of foldtuning (0.135), but grows steadily to 0.211 after four update rounds (Figure 2A). Aggregated across all rounds and targets, 25.5% of foldtuning-generated sequences are sufficiently devoid of homology to satisfy this escape criterion. When generated sequences do exhibit homology to natural proteins, locally aligning subsequences tend to decrease in length and identity with each additional round of foldtuning, supporting the contention that foldtuning gradually relaxes sequence limitations even for target structures that are under tighter apparent constraints (Figure S3). Fold-by-fold semantic change also captures a clear and steady progression away from natural sequences, from a median value of 39.9 following evotuning, to 46.9 after two rounds, to 56.8 after four (Figure 2B). Notably, up to the four rounds performed, foldtuning does not display any significant tradeoff between structural hit rate and sequence escape rate. In many cases, these metrics can be simultaneously maximized, as they are for, e.g. TIM *β*/*α*-barrels, immunoglobulin *β*-sandwich domains, and methyltransferases (Figure 2C).

**Figure 2.**
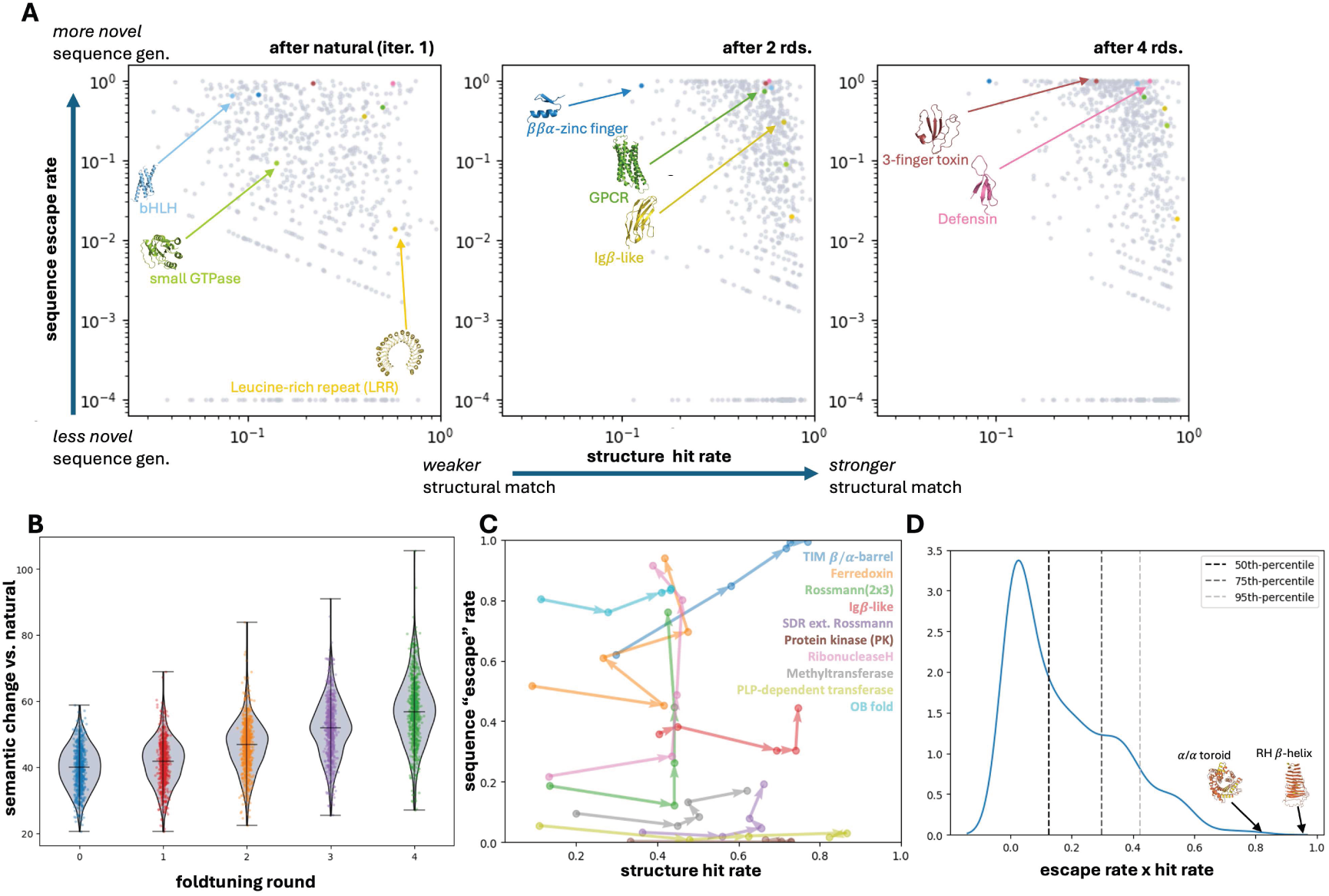
Foldtuned models readily sample novel sequences for > 700 targets. (A) Sequence escape vs. structural hit rates after natural-only evotuning or two or four rounds of foldtuning for 727 targets. Selected structural/functional targets are highlighted: transcription factors (blue), GPCRs/small GTPases (green), cell surface receptor domains (gold), and small antimicrobial/toxin proteins (red). (B) Semantic change between generated and natural sequences increases with additional rounds of foldtuning. (C) Sequence escape rate is maximized without compromising structural hit rate for 10 naturally-abundant target folds. (D) Ranking of target folds by sequence “designability”, as estimated by the product of structural hit and sequence escape rates.

Achieving a high structural hit rate and a high sequence escape rate suggests that a given fold tolerates substantial sequence plasticity without major disruption to structure; in other words, that fold is highly *designable*, being encoded by sequences numerous and variable. Treating the product of structural hit rate and sequence escape rate as a measure of “designability,” we find the right-handed *β*-helix, ribbon-helix-helix (RHH) domain, TIM *β*/*α*-barrel, anti-parallel *β*/*α* (PT) barrel, and *α*/*α* toroid ranked as the most designable SCOP motifs, followed by transmembrane *β*-barrels, Sm-like barrels, defensins, the winged helix domain, and the POU domain (Figure 2D, Table S2). Five of these ten motifs are symmetric or periodic in structure, three are transcription factor DNA-binding domains, and the remaining two – spliceosomal Sm proteins and defensins – exhibit ancient broad-spectrum activities as RNA-binders and antimicrobial agents respectively. Highly designable motifs span the four standard protein topology classes (all-*α*, all-*β*, *α* + *β*, and *α*/*β*); they are not restricted to the all-*α* helical bundles often derived from simple sequence design rules and that dominate PLM generative output in the absence of tuning or steering strategies.^13–16,45^ Furthermore, natural fold abundance in SCOP-UniRef50 is only weakly explanatory of designability, indicating that foldtuning detects inherent fold-to-fold variation in the strictness of sequence constraints with minimal bias induced by how evolution has sampled and diversified sequences (Figure S4).

### Foldtuning explores new sequence rules and familiar structure principles to build proteins

Having verified that foldtuning generalizes to several hundred targets covering structural and functional families of significant relevance to synthetic biology, we consider the finer-grained sequence features of foldtuning-generated proteins, taking G protein-coupled receptors (GPCRs) as an illustrative example and a target of substantial interest for engineering synthetic signaling systems. As foldtuned models propose GPCR sequences that drift further and further from natural training examples, structural fidelity to the target is preserved as far as high-level shape and connectivity, with the introduction of local plasticity on the order of a few-angstrom root mean square deviation (RMSD) in backbone atom coordinates vs wild-type (Figure 3A). For GPCRs, foldtuning achieves an aggregated structural hit rate of 0.550 and rapidly begins to generate far-from-natural sequences, dropping from a median sequence identity of 0.250 after the initial evotuning round on natural examples to the median sequence having no detectable homology against UniRef50 after the first round of foldtuning, a trend that persists over all four rounds (Table S1, Figure S3D). All-against-all pairwise sequence alignment of n_1_ = 2703 fold-tuned GPCRs and n_2_ = 34327 SCOP-UniRef50 entries reveals that at the sequence level, most foldtuned variants cluster into distinct subpopulations infilling regions of sequence-space not sampled by nature; more than one-in-seven (14.3%) are “orphan” variants dissimilar from natural sequences and from one another, appearing as isolated nodes in a network representation of sequence relationships (Figure 3B). Comparing amino-acid variation and conservation between natural and foldtuned GPCRs across all clusters and sequence positions, we found that foldtuned GPCRs consistently tolerate more variation (higher entropy), particularly at transmembrane positions – a departure from previous findings that diversity in PLM-generated sequences is concentrated in solvent-exposed and/or ligand-binding regions (Figure 3C).^33^ Individual foldtuned sequence clusters correspond to distinct strategies for replicating GPCR structure with site-specific amino-acid preferences that differ from nature (highest relative entropy). Across sequence clusters, foldtuned GPCRs exhibit substantial flattening in amino-acid usage distributions and widespread departure from the natural consensus, consistent with reduced position-wise conservation (Figures 3D–3F). Structurally, foldtuned GPCR models focus on redesigning the interfaces between the H1, H2, and H7 transmembrane helices; different clusters demonstrate remodeling of non-overlapping networks of hydrophobically packed and buried polar positions (e.g. T53, N54, N302 in cluster 1; N33, D64, R123, L314 in cluster 2) (Figures 3E–3F). These compensatory changes among residues in physical contact and the global magnitude of sequence modification would likely face substantial fitness barriers in stepwise evolution, but are detectable under foldtuning’s self-directed search thanks to its prioritization of large semantic displacements in sequence-space.

**Figure 3.**
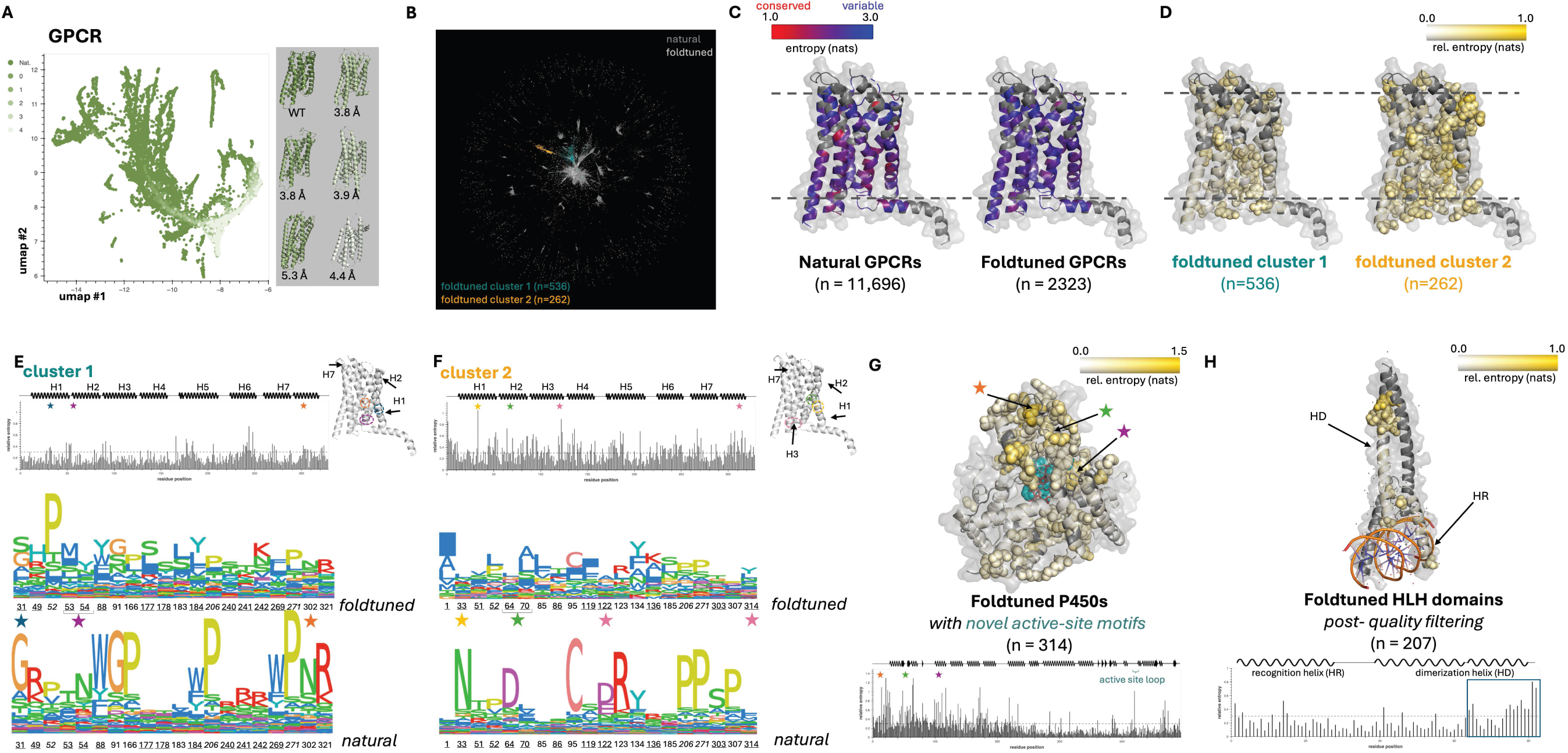
Foldtuning mimics target structures by globally rewiring constellations of interacting and/or functional residues. (A) UMAP of round-by-round foldtuning sequence diversification captured by ESM2-650M final-layer hidden states for G protein-coupled receptors (GPCRs, SCOP: 2000339). (B) Network representation of sequence similarity between natural (dark gray) and foldtuned (light gray) GPCRs; two subpopulations of foldtuned GPCRs are colored according to Louvain clustering assignments. (C) Position-wise entropy of amino-acid usage for natural (left) and foldtuned (right) GPCRs, mapped onto a representative structure of the *R. norvegicus* succinate receptor (pdb: 6IBB).^46^ (D) Position-wise relative entropy for foldtuned GPCR clusters from (B) vs. natural GPCRs (mapped onto pdb: 6IBB).^46^ Residues with relative entropy ≥ 0.3 nats are shown as space-filling spheres. (E) For foldtuned GPCR cluster 1 (n = 536) – *top*: Position-wise relative entropy mapped against secondary structure features; *bottom*: Sequence logos of amino-acid usage compared to natural GPCRs for the top-20 positions ranked by relative entropy. <u>Underlined</u> positions differ between cluster and natural consensus sequences; *italicized* positions are shared between clusters 1 and 2. Other key positions/contacts are denoted by colored stars. (F) Same as (E), for foldtuned GPCR cluster 2 (n = 262). (G) Position-wise relative entropy for foldtuned P450 enzymes with new-to-nature active-site heme-scaffolding loops (n = 314). Mapped onto – *top*: a representative structure of human P450 2C9 (pdb: 1R9O); *bottom*: secondary structure features.^47^ Key positions/contacts are denoted by colored stars. (H) Position-wise relative entropy for foldtuned helix-loop-helix (HLH) transcription factors, filtered to remove low-quality C-terminal regions (n = 207). Mapped onto – *top* a representative structure of the DNA-bound Mad-Max heterodimer (pdb: 1NLW); *bottom*: secondary structure features.^48^

This large-scale, spatially distributed epistasis phenomenon is similarly witnessed when foldtuning is applied to other functional protein targets. For the ubiquitous P450 enzyme super-family of monooxygenases, foldtuning achieves a 0.687 structural hit rate and a 0.367 median sequence identity vs natural examples (Table S1). Although foldtuned P450s retain shadows of detectable sequence homology, they liberally sample new active site electronic motifs – the 589,443 P450s cataloged in InterPro cover 24,228 unique versions of the crucial ten-residue heme-scaffolding loop, and foldtuning adds 295 more (occurring in 314/2996=10.5% of generated variants) that respect deeply-conserved heme-binding positions yet are unseen in nature (Figure 3G).^49^ Foldtuned P450s in this subset display novel amino-acid preferences in a three-dimensionally compact and contiguous region located between the active site and N-terminus (Figure 3G). At such an widespread level of sequence perturbation, it can be reasonably speculated that foldtuning is proposing seeds of new P450 families, pointing perhaps to new selectivity and activity for nature’s plastic and promiscuous xenometabolic reductases.^50^ Separately, when applied to the DNA-binding domain of basic helix-loop-helix (bHLH) transcription factors, foldtuning records a 0.568 structural hit rate and a 0.804 sequence escape rate (Table S1, Figure S3F). Foldtuned bHLHs mirror nature in amino-acid preferences where direct contacts are made with the DNA backbone and major groove, but vigorously sample laxity in the dimerization helix, both suggesting an avenue for nature-compatible engineering of transcriptional programs and hinting at minimal design determinants for *α*-helical dimers (Figure 3H).^51^

We also considered whether beyond exploring new sequence semantics at a global level, foldtuning might be favoring different “vocabularies” in its preferences for short local subsequences. To characterize vocabulary trends among foldtuning-generated sequences, we conducted an *n*-gram-based “vocabulary” analysis of foldtuned variants compared to SCOP-UniRef50 examples, splitting sequences into sliding windows of length 1-4 and calculating the usage frequencies of the 20, 400, 8000, and 16000 possible 1-grams, 2-grams, 3-grams, and 4-grams respectively. Considering the 12 most-abundant natural folds per the SCOP-UniRef50 database, all of which contain > 50, 000-250, 000 wild-type examples, we observe noticeable “vocabulary shifts” – that is, statistically significant upwards or downwards changes in *n*-gram frequency – among foldtuned sequences relative to natural ones for n = 1-4 across all folds analyzed (Figures S5–S8). For n = 1 (equivalent to simple amino-acid composition), 85-100%, or 17 to 20 of the twenty proteinogenic amino acids, shift in usage (Figure S5). For n = 2, 79.0-94.5% of dipeptide “words” shift (Figure S6). For n = 3, 26.5-75.9% of tripeptides shift (Figure S7). And for n = 4 – a length sufficient as a feature extractor for classifying protein families – as few as 5.7% (Rossmann2x3oid) and as many as 23.3% (PLP-dependent transferases) of “words” shift in one direction or the other (Figure S8).^52^ The substantial vocabulary shift magnitudes support the contention that foldtuning is stringing new local choices of subsequence motifs into globally perturbed full protein sequences – proposing novel fold-specific sequence languages in lieu of memorizing natural ones. This claim is reinforced by observing that rank-ordered *n*-gram usage by foldtuned models follows the same power-law-like distribution as within natural folds – identities of favored and disfavored short motifs change with foldtuning, but semantic breadth is still sampled without compression or collapse to a narrow non-diverse vocabulary (Figures S9–S12).

### Foldtuning is an implicit innovator of structure and function

We further observed that over the four rounds of foldtuning, without any explicit structural direction to do so, subsets of predicted structures tweak and elaborate on their formal backbone templates, discovering alterations both subtle (e.g. shortening disordered loops, rotating helices) and more substantial (e.g. reversing strand connectivity or altering global symmetry). For instance, the TIM *β*/*α* -barrel – shared throughout sequentially and functionally diverse enzyme families – undergoes rampant structural exploration in the course of attaining impressive structural hit (0.298 after evotuning to 0.770 after four rounds of foldtuning) and sequence escape rates (0.621 after evotuning to 0.995 after four rounds). All-against-all global structural alignment and clustering separates foldtuned TIM barrels into six prominent clusters, with only one cluster matching the expected 8-fold symmetry of the wild-type TIM barrel (Figure S13A).^53,54^ A second cluster ornaments the canonical barrel with a non-terminal surface *β*-hairpin resembling a putative natural feature also found in predicted structures of cofactor-F420-utilizing bacterial redox proteins. The remaining four clusters correspond to 9-fold, 10-fold (two subpopulations differing slightly in the manner of barrel closure), and 11-fold symmetries, none of which are known to nature according to experimental or predicted structure databases. Applied to fold-tuned immunoglobulin-like domains, the same structural clustering procedure distinguishes between subpopulations that differ in the relative orientations of the two sandwiched *β*-sheets, strand crossover locations, and loop geometry (Figure S13B).

Given the magnitude and diversity of sequence and structure modifications introduced by foldtuning, we evaluated the physical plausibility of these sequence-and-stucture-perturbed foldtuned proteins *in silico* by scoring their predicted structures with Rosetta to obtain ground-state energy estimates. For eleven target folds of interest, we compute estimated energies for all filtered and validated foldtuned variants and compare to n ≈ 100 natural training examples (Figure 4). For all eleven, foldtuned variants sit in the -(1-3) REU/aa regime typically invoked to distinguish physically reasonable structures from frustrated ones.^55^ The energy estimate distributions for many targets – e.g. *ββα*-zinc fingers, barstar-like proteins, defensins, GPCRs, small GTPases, basic helix-loop-helix (bHLH) domains – substantially overlap with those of wild-type examples. Co-localization is less complete for others such as immunoglobulin-like domains, SH3 domains, 3-finger toxins, and TIM barrels (Figure 4). This trend, combined with subtle shifts towards higher energies in later rounds, suggests that foldtuning gets gradually more permissive as far as energetics without abandoning the target fold, trading a slight penalty in folding stability vs wild-type for increased sampling into novel sequence basins. Still, when we score variants generated from 55 foldtuned models – targets chosen for potential use in bioengineering applications as hydrolase and oxidoreductase enzymes, nucleases and base-editors, kinases, proteases, and various scaffolds and mediators of catalysis and protein-protein interactions – with a PLM-based thermostability predictor, we see that significant fractions of fold-tuned proteins are predicted to exhibit melting temperatures > 60^◦^C, adding confidence that despite the level of remodeling involved, the far-from-natural artificial sequences unearthed by foldtuning encode realistic new proteins (Figure S14).^56^.

**Figure 4.**
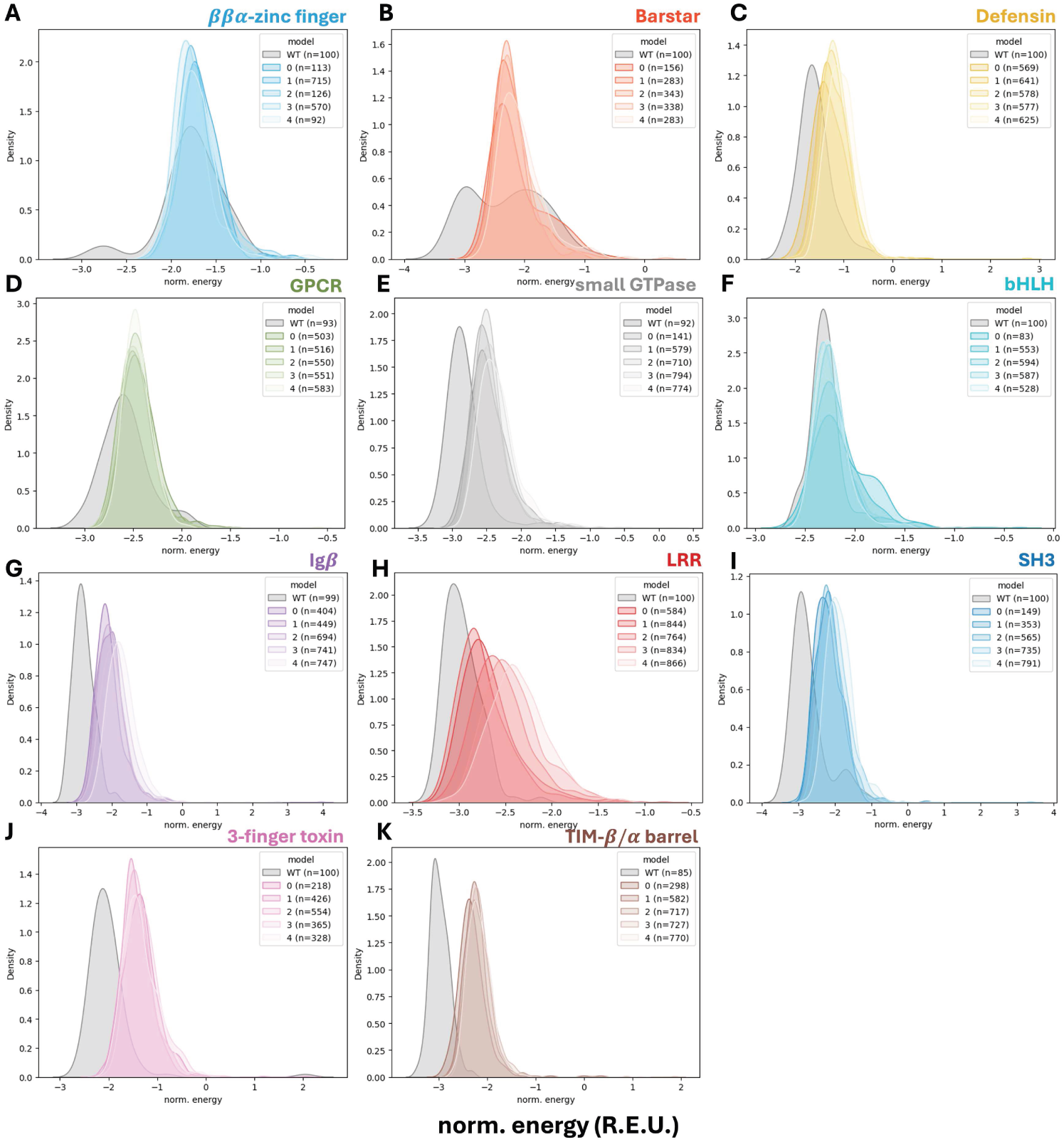
Foldtuning explores new basins in sequence-space by gradual adjustment of energetic constraints relative to nature. Histograms of length-normalized (REU / residue) Rosetta energy estimates for foldtuned (colored) and natural (gray) variants over five total rounds. Selected folds – (A) *ββα*-zinc finger. (B) Barstar. (C) Defensin. (D) G-protein coupled receptor (GPCR). (E) Small GTPase. (F) Basic HLH transcription factor (bHLH). (G) Immunoglobulin *β*-sandwich (Ig*β*). (H) Leucine-rich repeat (LRR). (I) SH3 domain. (J) Three-finger toxin domain (3FTx). (K) TIM *β*/*α* barrel.

Moving down one more level of information flow, we considered whether foldtuned proteins might recapitulate or even extend the functional capabilities of their parent folds. To this end, foldtuned variants for several SCOP folds corresponding to specific enzyme families or widely-distributed enzyme scaffolds (i.e. catalyzing diverse chemical transformations across nature) were assigned putative **E**nzyme **C**ommission classification numbers (EC #s) with a PLM-based predictor.^57^ For families with established reactivities and mechanisms, top-level EC #s are largely predicted as expected – P450s and nitrite/sulphite reductases are assigned as oxidoreductases, CRISPR Cas1s and *α*/*β* hydrolases are assigned as hydrolases, protein kinases are assigned as transferases, and chelatases are are assigned as lyases and ligases, covering their multiple roles in cofactor biosynthesis (Figure S15). Significant fractions of fold-tuned enzymes are annotated into categories associated with evolvability and and promiscuous activity against broad substrate spectra; for example, nearly one-in-five foldtuned P450s is labeled into EC 1.14.14.1, the catch-all “unspecified monooxygenase” category associated with the emergence of xenometabolism biocatalysts (Figure S15A). Similarly, foldtuned versions of CRISPR Cas1 – a metal-dependent non-site-specific DNA-specific endonuclease – are alternately labeled as site-specific, as exonucleases, or even as reverse transcriptases, pointing to fertile ground for engineering stable and sequence-specific gene-editing proteins from foldtuned starting points (Figure S15C). Foldtuned protein kinases span predicted activity as serine/threonine kinases (often with unknown or ambiguous specificity), (receptor)-tyrosine kinases, and as dual-specificity kinases capable of acting on all among serine, threonine, and tyrosine residues, presaging utility in designing bespoke signaling networks (Figure S15E). Foldtuned versions of common enzyme scaffolds, meanwhile, are typified by consistent annotation coverage spread across the six top-level EC reaction types, suggesting that foldtuning retains functional breadth when learning the sequence determinants of nature’s most widely-used and frequently repurposed domains (Figure S16).

### Foldtuned SH3 domains express stably

Emboldened by the ability of foldtuning to readily propose plausible far-from-natural protein sequences – as prefiltered computationally by structure prediction, search, and assignment – we sought to validate selected examples experimentally for expression and function. From a roster of small folds (≤ 84aa) for which coding DNA oligo pools could be easily synthesized, we focused first on the SH3-like barrel (SCOP ID: 2000090). The SH3 domain is a notable mediator of protein-protein interactions and regulator of signal transduction, particularly in tyrosine kinase pathways. Engineered SH3 domains have historically been desirable in synthetic biology for roles in designed artificial protein recognition and signaling cascades, but their utility has been plagued by *β*-barrel design difficulty and a lack of orthogonality to natural SH3s.^58^ Applying the standard evo+four foldtuning procedure to ProtGPT2 with SH3s as the target produced 2593 variants after *in silico* filtering, for a structural hit rate and sequence escape rate of 0.519 and 0.310 respectively. In contrast to, e.g. deep-mutational scanning libraries, proteins in foldtuned variant libraries – including for SH3s – boast high sequence diversity, featuring low pairwise sequence similarities and unique proteolytic digestion signatures (Figures S17A–S17C). This enables direct high-throughput characterization of protein expression and select biophysical properties by mass-spectrometry-based proteomics without the additional complexity of direct genotype-phenotype coupling methods such as yeast-, mRNA-, or cDNA- display.^59,60^ For the foldtuned SH3 library, 1347/2593 (51.9%) variants express at detectable levels in a reconstituted transcription-translation system as measured by untargeted mass-spectrometric profiling (Figures 5A–5C). Using length-normalized signal as a proxy for absolute abundance of expressed proteins, we observe signal intensity spanning ∼ 6 orders of magnitude, suggesting substantial variance in the intrinsic expressability of foldtuned SH3s; expression level neither correlates with sequence similarity to natural SH3s nor depends on the number of foldtuning cycles performed (Figure 5B).

**Figure 5.**
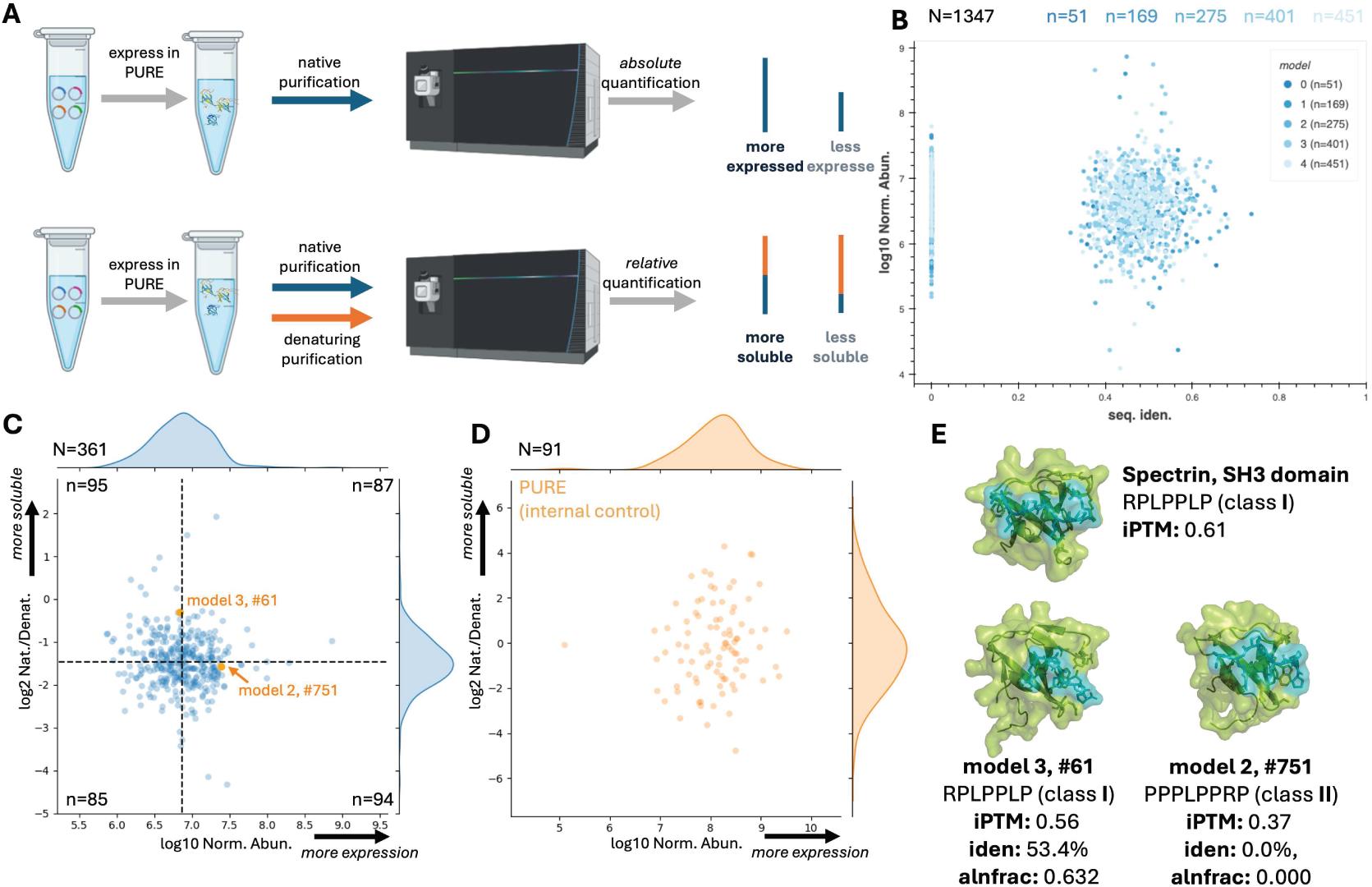
Foldtuned SH3s are expressable and stable. (A) Schematic of mass-spectrometry-based proteomics assays for variant library expression and folding stability. (B) SH3 expression assay signal intensity normalized by expected tryptic peptide count vs. sequence identity to closest UniRef50 hit for variants generated after 0-4 rounds of foldtuning (total N = 1347). (C) SH3 folding stability vs. expression assay results for N = 361 variants detected in both screens. Folding stability is measured by relative abundance ratio between native and denaturing purification fractions. Normalized expression is measured as as in (B) (D) Folding stability vs. expression for internal control set of N = 91 PURExpress components. (E) AlphaFold3 predicted structures and iPTM scores for selected SH3 variants (green) bound to a class-I or -II proline-rich peptide (teal), compared to the wildtype *G. gallus* spectrin SH3 domain.

To rule out cases where high cell-free expression intensity might mask solubility and/or aggregation issues from poor folding stability we compared foldtuned protein recovery under native and denaturing purification conditions; variants without folding pathologies are expected to show equivalent or greater signal in the native fraction relative to the denatured one (Figure 5A). Analysis of the native/denatured signal fold-change for an internal control of N = 91 *E. coli* proteins originating from the reconstituted transcription-translation system demonstrates that this stability/solubility proxy has a dynamic range spanning up to ∼ 10 orders of magnitude under the instrument conditions used (Figure 5D). A total of 361 variants are detected confidently in both the absolute and relative expression assay including a subpopulation of 87 foldtuned SH3s that are both highly abundant in the initial expression assay and displaced away from the denatured fraction in the solubility/aggreggation assay, suggesting expressability, relative folding stability, and low aggregation propensity (Figure 5C). We reasoned that foldtuned SH3 variants with high expressability and relative folding stability might recognize the proline-rich peptide motifs found in the binding partners of natural SH3 domains [61]. Indeed, *in silico* screening with AlphaFold3 predicts that certain physically-plausible foldtuned SH3 variants can bind either class I or class II proline-rich ligands in a hydrophobic aromatic-sidechain-rich cleft analogous to the wild-type interface as exemplified by the *G. gallus* spectrin SH3 domain (Figure 5E).^62–65^ One such example emerging from the AlphaFold-based screen, variant 3_61, is a distant homolog of the guanine nucleotide exchange factor Vav (involved in cytoskeletal remodeling during lymphocyte development and activation) and is predicted to recognize the canonical class I motif RPLPPLP. Another, variant 2_751, has no detectable sequence homology to any known protein, yet is predicted to recognize the canonical class II motif PPPLPPRP.

To clarify how foldtuned models might be preserving critical structural and functional features in SH3s, including ones responsible for stability or binding of polyproline motifs, we performed statistical coupling analysis (SCA), applied separately to natural SH3 domains and to the 2593 foldtuned putative SH3s.^17,18,66,67^ For both natural and synthetic SH3s, SCA retrieves a single sector of statistically-interacting residues (Figure S18). Natural and synthetic sectors are composed primarily of non-overlapping sets of core residues; only a single sector position interacts directly with the bound proline-rich motif and it is shared between the natural and synthetic cases (Figures S18C, S18G). This suggests that, consisent with the promiscuity and diversity of SH3-ligand binding, foldtuning is preserving a bare-minimum sequence rule for binding few-among-many polyproline-like targets (or even recognition of atypical peptide ligands at overlapping/alternate binding interfaces), while trying out novel evolutionarily-displaced solutions for stably assembling the SH3 *β*-barrel core.^68^

### Foldtuned barstars rescue bacteria from barnase toxicity

We next consider the barstar-like fold (SCOP ID: 2000624). With a single 3-stranded parallel *β*-sheet, the barstar-like fold is the simplest *α*/*β* family and features a well-described concerted folding pathway.^69,70^ Barstar’s native function *in vivo* is to inhibit, through a high-affinity active-site-occluding non-covalent interaction, the potent bacterial ribonuclease barnase before it is secreted into the surrounding environment. Applying foldtuning to barstar yields 1403 variants after *in silico* filtering, for a structural hit rate and sequence escape rate of 0.281 and 0.560 respectively. Variants were co-expressed with barnase from *B. amyloliquefaciens*; functional variants are expected to rescue host *E. coli* from the lethal effects of barnase expression in the absence of barstar (Figure 6A).^71^ Comparing sequencing counts of variant-coding amplicons, we found that 11 foldtuned barstar variants were significantly enriched (p < 0.05; Binyami-Hochberg correction for correlated tests) relative to uninduced (non-barnase-expressing) control under strong induction of barnase-barstar-variant co-expression, suggesting that the enriched variants are sufficiently functional mimics of barstar so as to mitigate the toxicity of barnase (Figure 6B). Additionally, enrichment does not correlate with sequence identity relative to wild-type barstars or any natural protein. To this point, 7/11 of survival-enriched foldtuned barstars do not exhibit any detectable homology to natural sequences at the domain or sub-domain level (Figure 6C).

**Figure 6.**
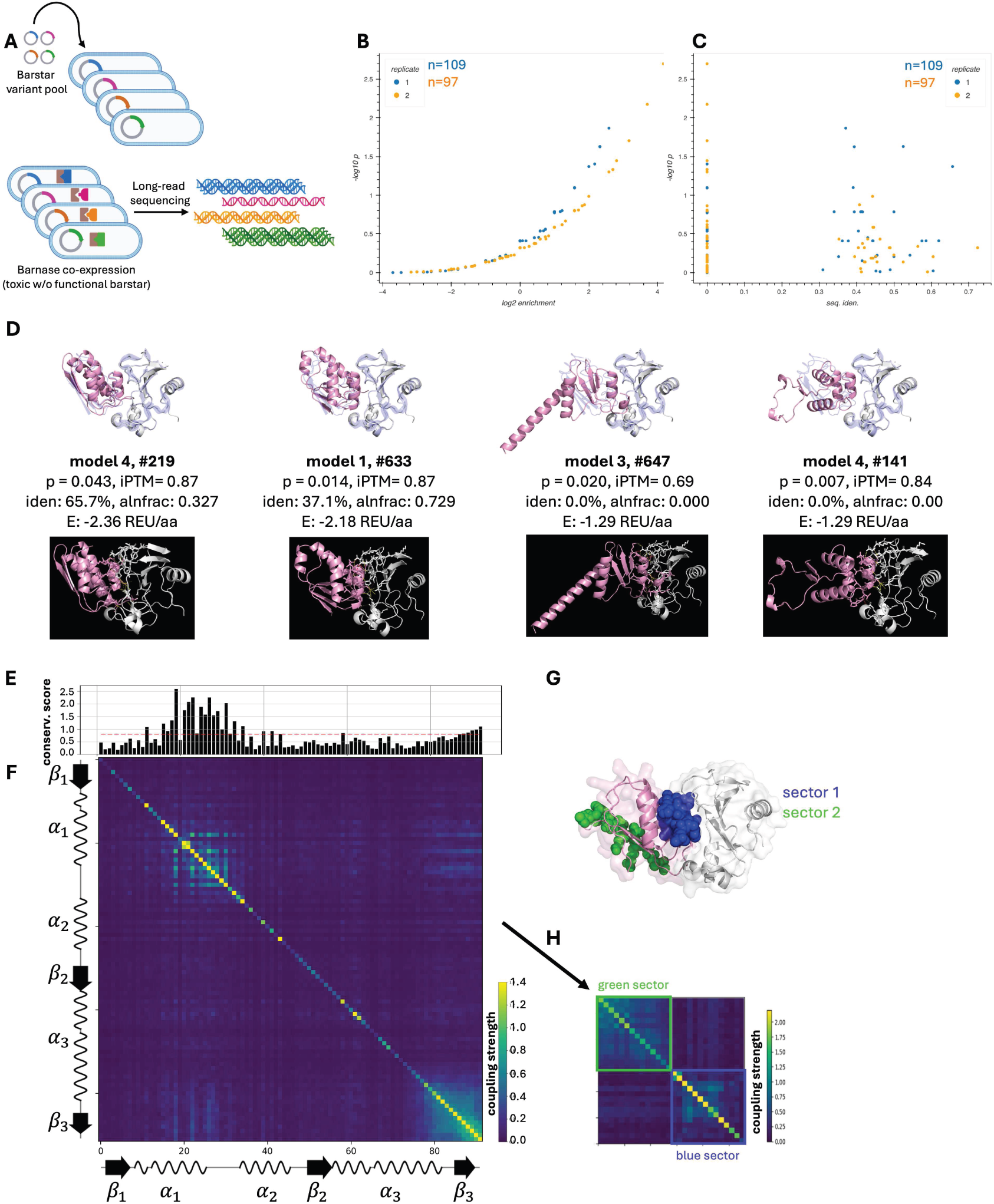
Functional foldtuned barstar variants reflect a minimal structure-function “grammar”. (A) Schematic of barnase-inhibition survival assay for barstar variant library stability and function. (B) Survival assay p-value rank plot. Enrichment of a given variant is calculated as the ratio of amplicon sequencing reads with and without induction of of barnase co-expression. (C) Survival assay p-values from (B) vs. sequence identity to most-similar UniRef50 hit. (D) Top row: AlphaFold3 predicted structures, iPTM scores, and Rosetta energy predictions for selected barstar variants (pink) in complex with barnase (white) with experimental crystal structure of the wildtype *B. aquaforiensis* barnase-barstar complex (pdb: 1BRS) overlaid in blue.^72^ Bottom row: Predicted complex structures with putative hydrogen bonds and electrostatic interactions indicated. (E) First-order positional conservation scores (divergence) from statistical coupling analysis (SCA) on n = 1493 foldtuned barstar sequences. (F) SCA Second-order positional correlation matrix for foldtuned barstars. (G) Visualization of SCA-identified sectors (blue, green) mapped onto a the barstar-barnase complex structure (pdb:2ZA4).^73^ (H) Compressed coupling matrix, blocked into two orthogonally “co-evolving” sectors identified by SCA.^73^

For mechanistic insight, we obtained AlphaFold3-predicted structures of the survival-enriched variants in complex with barnase. For four foldtuned variants – model 1 #633 (1_633), model 3 #647 (3_647), and model 4 #s 141 (4_141) and 219 (4_219) – these predicted complex structures indicate that barstar mimics are expected to bind barnase analogously to wild-type barstar, inserting an *α*-helix and adjoining loops into the binding pocket, obstructing the RNA hydrolysis active site (Figure 6D). Detailed examination of predicted binding interfaces reveals that foldtuned barstars are expected to form hydrogen-bonds and salt-bridges with barnase, without steric or electrostatic clashes. Comparison with a published experimental structure of the endogeneous *B. amyloliquefaciens* barnase-barstar complex (pdb: 1BRS) suggests that fewer such contacts are expected with variants than with wild-type barstar, potentially indicating weaker binding; these differences may alternatively stem from non-idealized bond geometry predictions by AlphaFold3 in the absence of a side-chain or backbone relaxation step (Figure 6D).^65^

Given the detection of foldtuned barstar mimics with antitoxin-like function and indications that at these mimics may utilize similar structural solutions to wild-type barstar, we asked what sequence and/or structure “rules” the relevant foldtuned models have learned and applied. Multiple sequence alignment of wild-type barstar along with the eleven survival-enriched fold-tuned variants reveals that in the contiguous nineteen-residue region (columns 38-56) spanning the barnase-binding interface, toxicity-rescuing variants preserve 6-11 (32-58%) of wild-type amino-acid identities, demonstrating that foldtuned models are not simply memorizing the semantics of barnase-binding and inserting them into redesigned flanks (Figure S19). Treating the n = 1403 foldtuned barstar variants as a synthetic protein family and repeating SCA reveals a prominent sector atop the barnase-binding interface (Figures 6E–6H). This suggests that foldtuning solves the barnase-binding problem by distilling the structural-functional “grammar” of barstar into a single rule that captures higher-order sequence dependencies in the critical inserted *α*-helix motif, while inventing wholly new ways to fill in the more biophysically malleable remainders of the barstar fold.

### Foldtuned insulins are INSR binders

Lastly, we deployed foldtuning to design mimics of insulin, a high-value translational target outside of the initial set of 727 templates. Insulin has been a frequent subject of computational design efforts due to the potential of synthetic variants to lower production and distribution costs, as well as to modulate receptor-binding and downstream signaling efficacy.^74,75^ Insulin poses an attractive challenge for foldtuning due to complexities including: (1) deep sequence conservation across eukaryotes, (2) a backbone structure neighborhood shared with related peptide hormones (e.g. IGF-1, relaxin, and ILPs) that exhibit multifaceted functions and receptor cross-reactivity, and (3) the extensive post-translational processing required to transform inactive proinsulin into active insulin through internal cleavage and intra- and inter-chain disulfide bond formation.^76^ To address the lattermost point and to align with typical expression and characterization processes in reactor-scale settings, we foldtuned ProtGPT2 to generate single-chain insulin variants that fuse the two (A- and B-) chains comprising active insulin and exclude the internal C-peptide region excised from proinsulin. Natural training sequences (n = 335, reduced to n = 193 after deduplication clustering) and reference structures were taken from InterPro (entry: IPR004825). Sequences were aligned to *H. sapiens* insulin to identify and remove putative C-peptide regions, yielding single-chain fusion training data. Foldtuning yielded 2889 putative insulin variants with structure hit and sequence escape rates of 0.578 and 0.0007 respectively; this structure hit rate, on par with typical foldtuning performance, validates the single-chain fusion approach, while the atypically low sequence escape rate, accompanied by a median 80.0% sequence identity to the closest natural counterpart of each foldtuned variant, highlights the difficulty of learning sequence-structure-function constraints from small training sets with high degrees of sequence conservation.

We used the Protein CREATE platform to screen all foldtuned putative insulins for INSR-specific binding as described in Lourenço et al.^77^. In brief, variants are displayed on T7 bacteriophage and screened against multiple receptor candidates ligated to magnetic beads, with a sequencing-based readout of amplicon counts before- and after- receptor-bead pulldown, resulting in a vector of enrichment scores across a palette of receptors. Foldtuned putative insulins were screened against the endogenous insulin receptor INSR for on-target binding and against IL7RA, a type I cytokine receptor, for generic off-target binding. For 307 foldtuned variants with sufficient reads to calculate enrichment in both channels, 41 (13.4% of detected; 1.4% of entire foldtuned pool) are enriched (enrichment > 1) for INSR-binding and de-enriched (enrichment < 1) for IL7RA-binding, suggesting specific binding activity against INSR (Figure 7A). AlphaFold3 predicts that the variants with the highest relative enrichment scores (#3_781, #4_404) bind to the INSR ectodomain through ligand-receptor contacts that partially, but not identically, overlap those formed by wild-type insulin, further supporting the notion that foldtuning retains only those sequence rules minimally necessary for structure and stability while injecting novelty that percolates to binding sites, behaviors, and downstream phenotypes. (Figure 7B).

**Figure 7.**
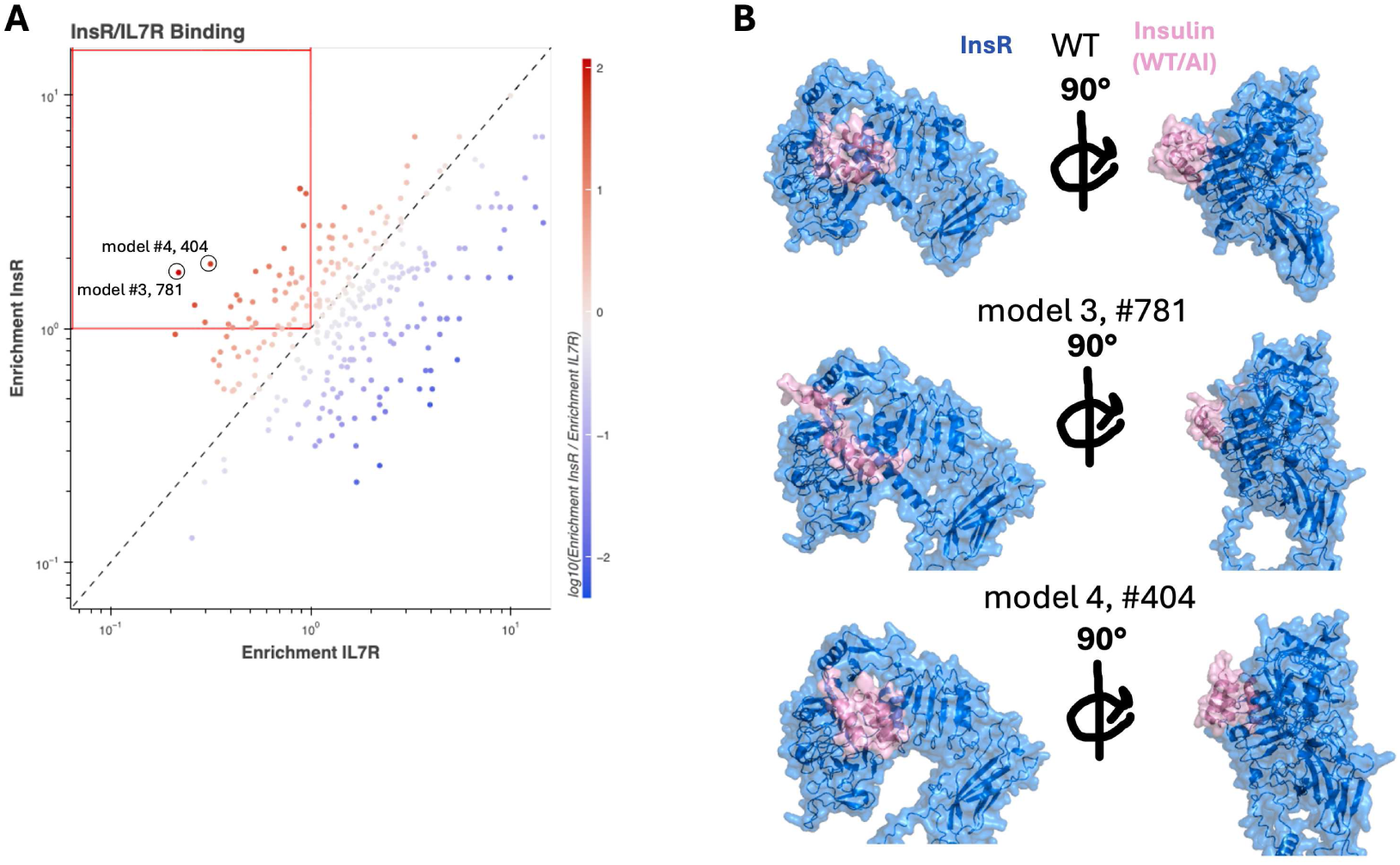
Foldtuned insulin variants bind the native insulin receptor. (A) Relative enrichment plot of foldtuned insulin variant binding to the native INSR receptor (on-target) vs the native IL7R receptor alpha-chain (off-target) using the Protein CREATE platform. (B) AlphaFold3 predicted structures of WT *H. sapiens* insulin and enriched foldtuned INSR binders (pink) in complex with the native INSR receptor ectodomain (blue).

## DISCUSSION

In this work, we introduced foldtuning, an adversarial learning paradigm that generates sequences predicted to fold into structures identical to those of natural proteins despite altogether lacking or possessing only minimal homology to sequences sampled by existing biology. Conceptually, foldtuning enables us to explore and organize the vast far-from-natural sequence space into pockets of related “protein doppelgängers” that map onto known structures. Through structural bioinformatics analysis of foldtuned sequences, we identify core biophysical constraints —- adhered to in different pockets via distinctive substantial epistatic shifts in amino-acid use —- that undergird the 3D architecture of critical protein families such as GPCRs and P450s. Practically, foldtuning generates libraries of novel proteins primed for engineering optimization and innovation, such as, for example, experimentally validated foldtuned binders of the insulin receptor. Our work produces structural mimetics for hundreds of protein families, providing a new resource for the rational design of far-from-natural proteins, a design space with potential benefits for reducing immunogenicity and witnessing emergent function.

In a departure from traditional sequence-based definitions, foldtuned models deduce biophysical definitions for protein families – interpretable strategies for solving requirements such as core packing and active-site scaffolding. While family definitions based on sequence homology are inherently limited in scope and biased by evolutionary selection for non-structural factors, the “evolution-free” populations reached by foldtuning are sufficiently diverse to elucidate higher-order features otherwise obscured by the stepwise fitness limitations of natural or directed evolution. More broadly, foldtuning hints at a far-from-natural protein sequence-space split into identifiable islands and pockets of common structure — miniature neutral nets separated from one other by combinatorial restrictions on amino-acid contacts that stem from the most basic principles of biomolecular interactions.^1,78^ Guided by this newly unveiled organization scheme, engineering directions for foldtuning are numerous, pointing to replacement of kinase binding domains to transduce phosphorylation between interaction partners never linked in nature, to composition of new ligands and cell-surface receptors that can send new signals and write new gene programs, or even to whole metabolic pathways and cellular structures, redesigned enzyme-by-enzyme and filament-by-filament in a steady advance towards nature-emulating synthetic cells.

Within the broader landscape of protein modeling and design, foldtuning highlights that the inherent sense of linguistic consistency captured by PLMs can be exploited to follow sequence trajectories that respect an overarching structure-specific grammar. In retaining enough plasticity to explore minor innovations, modifications, and degrees of freedom within structural families, foldtuning complements inverse-folding and diffusion methods that are optimized for sequence design towards strictly specified target backbones and free structure generation respectively.^24,25,79^ Moreover, in directly learning from synthetic data to push PLMs to operate at the limits of what is permitted by biophysics, foldtuning reinforces the emerging signs that high-quality synthetic data can expand the horizons of protein models towards maintaining fitness and boosting novelty in generative outputs, without experiencing model collapse.^80–84^ Looking forward, we envision that foldtuning will continue to grow in insight and applicability, through adaptions such as incorporation of bespoke scoring methods for individual protein engineering problems, reinforcement-learning-guided model updates based on experimental measurements, and combinatorial diversification of protein domains and subdomains for realization of novel biological pathways, programs, and phenotypes.

## METHODS

### Construction of the SCOP-UniRef50 Sequence-Structure Database

The SCOP-UniRef50 custom sequence-structure fragment database was constructed by performing reciprocal Foldseek searches (in fast TM-align mode) of the SCOP database of super-family representative PDB structures (n = 36, 900) against the UniRef50 portion (based on the 2021_04 release) included in the July 2022 update to the AlphaFoldDB (https://alphafold.ebi.ac.uk/) and made available as a precompiled Foldseek database, filtering for reciprocal hits with fractional query and target coverage > 0.8 and TMscore> 0.5, and clustering the filtered fragments at 100% identity.^38,85^

### Training, Validation, and Generation from Foldtuned Models

#### Target Fold Selection for Foldtuning

Out of 1562 folds categorized in SCOP v2, 1474 are present in the SCOP-UniRef50 database whose construction is described above. The top 850 most-abundant of these comprise the intial target set for foldtuning, a cutoff selected in part out of consideration for compute resource constraints and in part to exclude folds with potentially inadequate volumes of natural sequence starting material. As a second target set, motivated by functional protein engineering applications, we hand-selected 44 cytokine, chemokine, and growth factor entries from InterPro.^44^

#### Sequence Selection for Evotuning

For the preliminary foldtuning round on SCOP target fold f, termed the evotuning round, the base ProtGPT2 model was finetuned for 1-3 epochs on 100 natural sequences selected at random from the subset of sequences in the custom SCOP-UniRef50 database (construction described above) annotated to fold f.

For the evotuning round on InterPro target entry f_IP_, 100 natural sequences were selected at random from sequences associated with f_IP_ in InterPro v93.0, preliminarily clustered at 100% sequence similarity with mmseqs2 for deduplication and fragment removal.^44^

#### Finetuning of ProtGPT2

All finetuning of ProtGPT2 was performed with the Adam optimizer using a learning rate of 0.0001, and next-token prediction as the causal language modeling task. For the evotuning round, finetuning proceeded for 1-3 epochs, with the number of epochs for a specific SCOP fold f or InterPro fold f_IP_ determined by a pre-screen in which ProtGPT2 was finetuned for 1-5 epochs, generating 100 sequences per epoch, predicting and assigning structures as described below, and finding the minimum epoch such that ≥ 7% of sequences were assigned to fold f in order to ensure sufficient synthetic data to initiate foldtuning.

In subsequent foldtuning rounds, finetuning was performed with the same optimizer parameters, for 1 epoch only, on the top-100 previous-round sequences assigned to f or f_IP_ ranked in order of decreasing semantic change as described in the main text and below.

#### Sequence Generation from ProtGPT2

Sequences were sampled from ProtGPT2 by L-to-R next-token prediction with the default best-performing hyperparameters reported by Ferruz et al.;^31^ sampling temperature 1, top_k 950, top_p 1.0, repetition penalty 1.2. Per round, n = 1000 sequences were generated from the appropriate evotuned or foldtuned model. The termination condition was set following 0.4 × M tokens, where M is the median length of SCOP-UniRef50 natural sequences for target fold f, or the first STOP token, whichever occurred first; generated sequences were force-truncated to a maximum length of M aa. Truncated sequences containing rare or ambiguous amino acids (B, J, O, U, X, or Z) were filtered out as invalid. Inference batch size on a single NVIDIA A100-80G GPU ranged from 125-500 sequences depending on target sequence length.

#### Structure Prediction and Assignment

All structures were predicted with default ESMFold inference parameters as reported by Lin et al.^28^ Structures were inferenced in batches of 10-500, depending on sequence length, on single A100-80G GPUs, with compute resource collaboration through Oracle Cloud Infrastructure (OCI).

Predicted structures were annotated either to: (1) SCOP fold labels via Foldseek structure-based search against a custom database comprised of the n = 36, 900 superfamily-level representative structures in SCOP v2; (2) InterPro entry labels via Foldseek structure-based search against a custom database comprised of structures compiled from 44 chemokine, cytokine, and growth factor entries in InterPro v93.0. Irrespective of target databse, Foldseek was run in accelerated TMalign mode. The consensus SCOP fold or InterPro entry was defined as the fold/entry accounting for the most hits with TMscore > 0.5 and max(query_coverage, target_coverage) > 0.8. In the absence of at least one hit satisfying these criteria, a structure was considered to be unassignable.

#### Sequence Selection for Foldtuning

For each target fold f, f_IP_ and foldtuning round k = 1, 2, …N, the semantic change relative to natural versions was calculated for all generated sequences 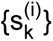 structurally assigned to fold f, f_IP_ as

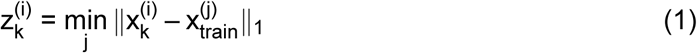

where 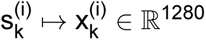 via embedding with ESM2-650M, and the “train” subscript denotes the natural sequences selected from SCOP-UniRef50 or InterPro for the initial foldtuning round.

The 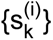 were ranked by their corresponding 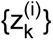 in descending-order; the top 100 comprising the finetuning sequence data for the (k + 1)-th round.

### *In Silico* Evaluation of Foldtuned Models & Outputs

#### Structural Hit, Sequence Escape, and Designability Rates

For a given foldtuned model with target fold f, structural hit rate was computed as the fraction of generated sequences with successful structure assignment to f. More formally, for a generated sequence s_i_ and fold f, it is Pr(s_i_ ∈ f). Sequence escape rate was computed as the fraction of *those sequences structurally assigned to the target* that do not return an alignment of any length to any cluster representative from UniRef50 in an mmseqs2 search with default easy-search parameters and maximum e-value 0.01.^86^ Or, formally, Pr(s_i_ ∉ N|s_i_ ∈ f), where we borrow N to stand in for the set of all natural/natural-resembling/homologous-to-natural sequences. The “designability” of a fold f was computed as the product of the corresponding structural hit and sequence escape rates, or d_f_ = Pr(s_i_ ∉ N|s_i_ ∈ f) × Pr(s_i_ ∈ f) = Pr(s_i_ ∉ N; s_i_ ∈ f).

#### PCA and UMAP Representations

Mean-pooled embeddings for natural and foldtuned sequences were inferenced with ESM2-650M and dimension-reduced from R^1280^ to R^100^ by principal component analysis (PCA) and further to R^2^ by Uniform Manifold Approximation and Projection (UMAP). For the eleven chosen folds depicted in Figure 1, Figure S1, and Figure S2, natural sequences were sampled from SCOP-UniRef50 at 5x the number of filtered, validated foldtuned sequences obtained after initial evotuning+four rounds.

#### Generation of Representative Control Sequence Sets

The median alignment length (n_f_) and sequence identity (l_f_) were calculated for a fold f by considering only the subset of foldtuned sequences returning an alignment to at least one cluster representative from UniRef50 in an mmseqs2 search with default easy-search parameters and maximum e-value 0.01.^86,87^. Synthetic control sequences were generated by treating natural sequences assigned to fold f in the custom SCOP-UniRef50 database as templates – for a template sequence of length n, a contiguous local amino-acid subsequence of length min{n, n_f_} was randomly mutated at (1 – f_f_) min{n, n_f_} positions and flanked with random amino-acids to the total length n. For a given fold f, 1000 synthetic control sequences were generated, 10 with each of the 100 natural sequences selected for evotuning as the template. Structure prediction and assignment was performed for synthetic control sequences as for ProtGPT2-generated sequences; structural hit rates and sequence escape rates were computed as for foldtuned models.

#### Sequence Similarity Analysis and Clustering

Sequence network analysis was carried out by separately preclustering foldtuned sequences and natural SCOP-UniRef50 sequence fragments assigned to fold f at 50% identity, via mm-seqs2 easy-cluster with default settings and covariance mode 1. Preclustered sequence sets were then merged and searched all-against-all using mmseqs2 easy-search with maximum e-value 10^−5^. Graph representations were constructed with preclustered sequences as nodes and edges joining pairs of nodes with reciprocal alignments of any length and no minimum sequence identity threshold. Node positions were calculated and visualized with networkx according to the force-directed representation of the Fruchterman-Reingold with spring constants k_ij_ ∝ { the max-normalized bit-score of the alignment between s_i_, s_j_} for all edges e_ij_.^88^ Cluster assignments were computed using the Louvain algorithm with default parameters as implemented in networkx.^89^

#### Sequence Conservation and Relative Entropy Analysis

Sequences were prepared for sequence conservation and relative entropy analysis by separately preclustering foldtuned sequences and natural SCOP-UniRef50 sequence fragments assigned to fold f at 50% identity, via mmseqs2 easy-cluster with default settings and covariance mode 1. Preclustered sequence sets were merged and multiple sequence alignments (MSAs) constructed with MAFFT in FFT-NS-2 mode.^90^ MSAs were processed to remove columns with a gap present in > 80% of rows. Position-wise entropy 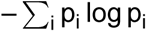 was computed as implemented in SciPy. Relative entropy 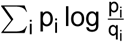 was computed as implemented in SciPy, with the {p_i_}, {q_i_} representing amino-acid usage frequencies for foldtuned and natural sequences respectively. Contributions from gap characters were ignored and {q_i_}’s were assigned a single artificial count before normalization where necessary to avoid the case where q_i_ = 0 and relative entropy is consequently undefined. Sequence logos were computed and visualized in biotite with default parameters. Mappings of entropy and relative entropy onto experimental structures were visualized with PyMOL v3.1.0. For foldtuned and natural (InterPro family IPR036396) P450 variants, active-site motifs were validated by regular expression pattern-matching to PROSITE pattern PS00086 (”Cytochrome P450 cysteine heme-iron ligand signature”; [FW]-[SGNH]-x-[GD]-F-[RKHPT]-P-C-[LIVMFAP]-[GAD]).^44,49^

#### Structural Similarity Analysis and Clustering

Structural clustering analysis for a fold f was carried out by conducting an all-against-all structural alignment of successfully assigned variants with Foldseek in fast TM-align mode. Missing values (no alignment passing filters) were imputed as having a TMscore of 0. Results were represented as a graph with individual variants as nodes, and an edge joining any pair of nodes with reciprocal average TMscore > 0.7. Cluster assignments were computed using the Louvain algorithm with default parameters as implemented in networkx to separate the network into fold motif clusters. Isolated nodes were excluded from clustering and visualization.^89^

#### Energy Scoring Calculations

Biomolecule energy scores were obtained using the default ‘ref2015‘ energy function and standard relaxation and scoring workflow in Rosetta v3.11.^55^ Energy scores are reported in **R**osetta **E**nergy **U**nits (R.E.U.), normalized to sequence length.

### Advanced Chemical Property Prediction and Visualization

Melting temperature bin predictions (T_m_) for thermostability were obtained for all foldtuned sequences using the 40^◦^C, 45^◦^C, 50^◦^C, 55^◦^C, 60^◦^C, and 65^◦^C binary classifiers released as part of TemStaPro v0.2.6.^56^

Functional enzyme reactivity annotation labels (**E**nzyme **C**ommission #s; EC#s) were inferred for thirty-one classes of foldtuned sequences using the fast “max-separation” mode of CLEAN v1.0.1.^57^ Where multiple EC#s were inferred for a given sequence, the closest centroid was retained as the best-scoring annotation. The full body of EC# annotations across all scored sequences for a given fold were visualized using KronaTools v2.8.1 with XML customization to maintain a consistent color scheme for top-level EC# classification: oxidoreductases (EC 1; red), transferases (EC 2; yellow), hydrolases (EC 3; green), lyases (EC 4; blue), isomerases (EC 5; purple), ligases (EC 6; pink).

### Sequence *N*-Gram Decomposition and Analysis

*N*-gram vocabulary analysis was carried out with custom code by splitting foldtuned sequences and SCOP-UniRef50 sequence fragments assigned to fold f into subsequences (”words”) of length 1,2, 3, or 4 and computing their respective frequency distributions and fold-change for foldtuned variants vs. natural SCOP-UniRef50 sequences. For each fold/word-length pair, n = 1000 non-parametric bootstrap replicates were drawn with the SCOP-UniRef50 sequences as the null distribution and significance testing for individual word frequency change performed at significance level *α* = 0.05, applying the Binyamini-Hochberg correction for positively correlated tests.^91^

### Statiscal Coupling Analysis

Statistical coupling analysis (SCA) was executed as implemented in pySCA v6.1 and as described in Rivoire et al.^92^ In brief: multiple sequence alignments (MSAs) of foldtuned sequences were obtained and pre-processed for SCA using FAMSA v2.2.3.^93^ MSAs were used as the input to statistical coupling analysis (SCA), which was performed with pySCA v6.1. Sequences with non-standard amino acids and sequences with more than 20% gaps were removed from the alignment. Additionally, columns with more than 20% gaps across all sequences were trimmed from the alignment. A reference sequence in the alignment was automatically selected and sequences with under 20% or over 80% sequence identity to the reference sequence filtered out. Finally, each sequence in the alignment was weighted by their similarity to the reference sequence to yield an MSA with dimensions M^′^ ×L, where M^′^ is the number of effective sequences, and L the number of alignment positions. This final processed MSA was used to calculate the sequence similarity matrix, first-order positional conservation, and second-order positional correlation matrix. Significant eigenmodes and independent components were identified, ranked, converted to physical sectors, and visualized, all as summarized in Rivoire et al.^92^

### Oligo Pool Design and Preparation

Foldtuning-generated sequences selected for experimental characterization were truncated to remove disorded N- and C-terminal tail regions as predicted by ESMFold and identified in C_*α*_ contact maps computed with biotite. Coding DNA sequences were designed by reverse translation with dnachisel, codon-optimizing for *E. coli*, with additional constraints on GC content (global ≥ 0.25, ≤ 0.65; never ≤ 0.19 or ≥ 0.71 over any subsequence of length 50) and homopolymers (restricted to < 14nt). Constant flanks – GACTACAAGGACGACGATGACAAG (5’) and GGTTCCCACCATCATCACCATCAT (3’) were added to code for a 5’ FLAG tag and a 3’ GSHHHHHH tag.

Oligo pools were ordered from Twist Biosciences as ssDNA fragments for sequences ≤ 300nt or as dsDNA fragments for sequences > 300bp and PCR-amplified with Q5 Hot Start High-Fidelity 2X Master Mix (NEB, M0494S) according to manufacturer instructions. T7RNAP promoter, ribosome binding site, start codon, stop codon, and T7 terminator elements were added in a subsequent PCR-amplification step with the same reagents, and purified, concentrated, and resuspended in ultra-pure water using the Monarch Spin PCR & DNA Cleanup Kit (NEB, T1130S) according to manufacturer instructions.

### *In vitro* Expression and Folding Stability Measurements

#### Protein Expression and Purification

Foldtuned variant pools were expressed *in vitro* with PURExpress (NEB, E6800) following the manufacturer’s protocol, with 500ng template dsDNA per 50 µL reaction volume, incubating 18hrs at 29 °C. Expressed protein was purified under native conditions by His-tag pulldown using NEBExpress Ni Spin Columns (NEB, S1427L); 400 µL of eluate was washed and concentrated with Amicon Ultra Centrifugal Filters, 3 kDa MWCO (Millipore, UFC5003) 4x with 400 µL phosphate-buffered saline pH7.4, centrifuging at 14,000*g* for 30min per exchange, and 50 µL of concentrate recovered by reverse spin (1000*g* for 2min). For the folding stability assay, the reaction volume was split post-expression into 2 × 25 µL aliquots, one purified under native conditions and the other under denaturing conditions (6 M guanidinium chloride) following manufacturer instructions.

#### LC-MS/MS

Concentrated purified protein samples were digested in an S-Trap micro spin column (Protifi, USA) according to the manufacturer’s instructions. Data was acquired in data dependent acquisition (DDA) mode on an Orbitrap Eclipse Tribrid mass spectrometer (Thermo Fisher Scientific, USA) coupled to a Vanquish Neo UHPLC system (Thermo Fisher Scientific, USA). 1ug of peptides were separated on an Aurora UHPLC Column (25 cm × 75 μm, 1.7 μm C18, AUR3-25075C18-TS, Ion Opticks) with a flow rate of 0.35 μL/min for a total duration of 1 hour and ionized at 1.8 kV in the positive ion mode. The active gradient was composed of 6% B (3.5 min), 6-25% B (41.5 min), 25-40% B (15 min), 40-98% B (2 min) and 98% B (5min); mobile phase A is 2% acetonitrile (ACN, Fisher Scientific, A9554) and 0.2% formic acid (FA, Fisher Scientific, A11750) in water (Optima LC-MS grade, Fisher Scientific, W6212), and mobile phase B is 8% ACN and 0.2% FA in water. MS1 spectra were collected in the Orbitrap at the resolution of 120,000 from 375 to 1,600 m/z with automatic gain control (AGC) target of 250% and a maximum injection time of 50 ms. MS2 scans were acquired in the ion trap using fast scan rate on precursors with 2-7 charge states and quadrupole isolation mode (isolation window: 1.2 m/z) with higher-energy collisional dissociation (HCD, 30%) activation type. Dynamic exclusion was set to 15 s and low/high mass tolerance was 5ppm. Ion transfer tube temperature was 300 °C and the S-lens RF level was set to 30.

RAW files were processed in Proteome Discoverer with SEQUEST (version 2.5, Thermo Scientific) against the UniProt Escherichia coli (strain B / BL21) proteome(UP000002032, 2021 version, 4,156 entries) and foldtuned variant sequences. Trypsin/P was set as the digestion enzyme, allowing a maximum of two missed cleavages. Dynamic modifications were set to oxidation on methionine (M, +15.995 Da), deamidation on asparagine (N,Q +0.984 Da) and protein N-terminal acetylation (+42.011 Da). Carbamidomethylation on cysteine (C, +57.021 Da) was set as a fixed modification. The maximum parental mass error was set to 10 ppm, and the MS2 mass tolerance was set to 0.6 Da. The false discovery threshold was set strictly to 0.01 using the Percolator Node validated by q-value. The relative abundance of parental peptides was calculated by integration of the area under the curve of the MS1 peaks using the Minora LFQ node. The mass tolerance used to align features across runs was set to 5 ppm in the Feature Mapper node.

For in vitro expression measurements, absolute abundance signal intensities were scaled by dividing by the expected peptide count from simulated tryptic digestion. For in vitro folding stability measurements, enrichment was calculated as the abundance ratio of the natural channel relative to the denatured channel.

### Barstar-Barnase Survival Assay

The barstar-like foldtuned variant pool was designed, ordered, and amplified to add regulatory elements as described above. Barstar variants were cloned as a single pool into barnase-barstar expression vector pMT416 (gift from Robert Hartley, Addgene plasmid #8607; http://n2t.net/addgene:8607; RRID:Addgene_8607), replacing the wild-type barstar-coding region, using NEBuilder HiFi DNA Assembly Master Mix (NEB, E2621S) according to manufacturer’s instructions. 1 µL of assembly product was transformed into 10 µL 5-alpha F’Iq Competent *E. coli* (NEB, C2992I) following the standard manufacturer heat-shock protocol. Outgrowth product was used to seed 2mL LB cultures at 1-in-200 dilution and incubated overnight at 37 °C, 250 rpm with carbenicillin as the selection marker. Upon reaching an OD600 of 0.6, cultures were split into two 1 mL aliquots; 1mM IPTG was added to one aliquot per pair, the other was kept as an untreated control; all aliquots were incubated at 37 °C for 3hrs to strongly induce protein expression. Barstar-variant-coding regions were amplified directly from 0.2 µL of culture using Q5 Hot Start High-Fidelity 2X Master Mix (NEB, M0494S). PCR product was purified as described above, diluted to 5 ng/µL, and Premium PCR Sequencing performed by Plasmidsaurus using Oxford Nanopore Technology with custom analysis and annotation.

Reads were translated and filtered to retain only protein sequences containing the expected N- and C-terminal tag leader sequences and not prematurely truncated by a misplaced STOP codon. Translated reads were mapped back to the foldtuning-generating barstar variant sequences with mmseqs2, requiring an aligned region of > 80aa with a minimum sequence identity of 98%. Variant enrichment was calculated as the ratio of mapped reads under barnase-barstar induction vs the uninduced control. P-values were computed non-parametrically by assuming a null model of random read allocation, drawing 10^6^ samples.

### Binding Mode Prediction and Analysis

Unless specified to the contrary, AlphaFold3 was used for all structure prediction tasks on complexes, via the AlphaFold-Server interface (https://alphafoldserver.com).^65^ For the SH3 domain, predicted complex structures were computed for foldtuning-generated putative SH3 variants in the presence of a representative class I (RPLPPLP) or class II (PPPLPPRP) proline-rich peptide motif. For the barstar-like fold, predicted complex structures were computed for foldtuning-generated putative barstar variants in the presence of wild-type barnase from *B. amyloliquefaciens* (uniprot:P00648). Predicted structures were compared to a wild-type reference, either the spectrin SH3 domain from *Gallus gallus* (pdb:1shg) or the barnase-barstar complex from *Bacillus amyloliquefaciens* (pdb: 1brs).^64,72^ For insulin, predicted complex structures were computed for foldtuning-generated and/or PLM-sampled putative insulin variants in complex with the monomeric full-length ectodomain of human insulin receptor INSR (uniprot:P06213).

All predicted structures were visualized with PyMOL v3.1.0. For the barnase-barstar complex, good hydrogen-bonds, acceptable hydrogen-bonds, and electrostatic clashes were inferred and displayed with the PyMOL “show_contacts” third-party plugin. For insulins, hydrophobicity was visualized using the “color_h” third-party plugin and electrostatic potential was calculated and visualized using the APBS Electrostatics plugin.

For toxicity-rescuing barstar variants, multiple sequence alignments (MSAs) were calculated using MUSCLE v5 via the EMBL-EBI webserver.^94^

### High-Throughput Insulin Binding Assay

A library of 2889 insulin variant amino-acid sequences was constructed by foldtuning on InterPro entry IPR004825, containing 335 natural insulin sequences (reduced to 193 sequences after deduplication clustering at 100% similarity with mmseqs2) integrated from overlapping entries in the PRINTS, CDD, and PANTHER databases.^44^ Foldtuning was executed as otherwise described, with the modification that generated variants were post-processed by aligning to the sequence *H. sapiens* insulin (uniprot: P01308) and removing residues aligning to the C-peptide region that is removed by proteolytic cleavage *in vivo* during the conversion of inactive proinsulin to active insulin, resulting in a library of *single-chain* insulin mimics.

High throughput binding measurements (sequencing read enrichment scores) were obtained using the Protein CREATE platform as described in Lourenço et al.^77^ with INSR as the on-target receptor and IL7RA as the off-target decoy receptor.

## Data and Code Availability

A streamlined implementation of foldtuning is distributed as part of the TRILL software package (https://github.com/martinez-zacharya/TRILL; https://pypi.org/project/trill-proteins/) from v1.8.3 onwards.^95^.

The mass spectrometry proteomics data have been deposited to the ProteomeXchange Consortium via the PRIDE partner repository with the dataset identifier PXD071745.^96^

Sequencing read data are deposited at CaltechDATA (doi:10.22002/24abg-6e603). Other inquiries should be directed to the corresponding author.

## Acknowledgements

We thank Steve Mayo, Richard Murray, Erik Winfree, and Carl Pabo, as well as all members of the Thomson Lab for helpful discussions and feedback. We thank Ashwin Rakhra, John Ng, and their colleagues at the Oracle Corporation for generous cloud computing support.

This work was supported by the National Institutes of Health under award number R01-GM150125, the Gordon and Betty Moore Foundation, the Caltech Rosen Bioengineering Center, and the David and Lucille Packard Foundation under a Packard Fellowship to M.W.T. The Proteome Exploration Laboratory is supported by the Caltech Beckman Institute Endowment Funds.

## Author Contributions

A.M.S. and M.W.T. conceptualized and developed the foldtuning framework. A.M.S. performed computational experiments and trained the foldtuned model library. Z.A.M. carried out software development and benchmarking. S.C.Y. carried out bioinformatics analyses. A.M.S., A.L.L., and S.L. designed and performed wet-lab experiments. A.M.S., A.L.L., and T.-F.C. designed proteomics experiments. T.-Y.W. performed proteomics experiments and analysis. A.M.S. and M.W.T. wrote the manuscript with input from all authors.

## Supplemental Figures

**Figure S1.**
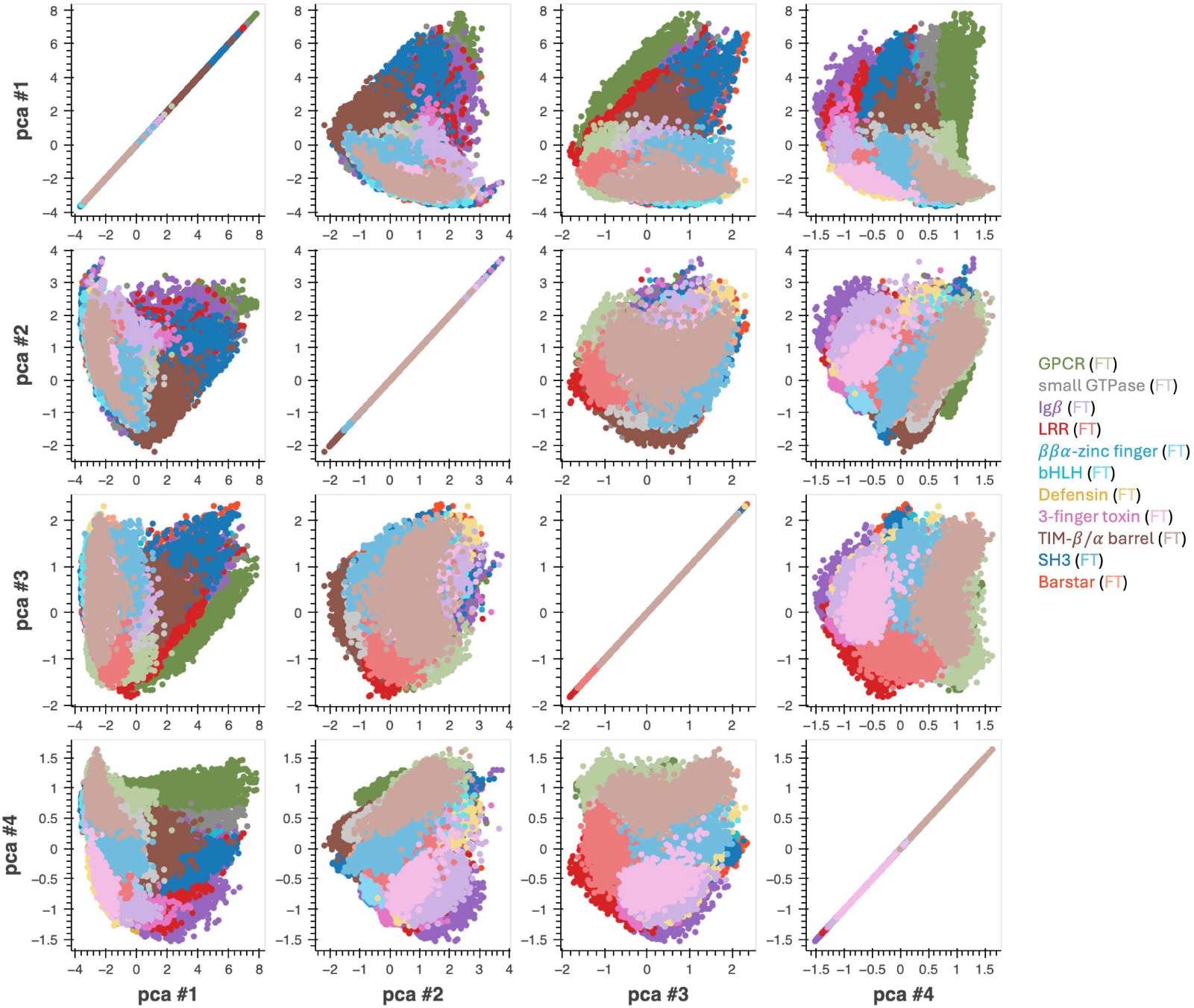
Principal component analysis (PCA) of ESM2-650M embeddings differentiates between natural and foldtuned versions of diverse structures. Pairwise plots of top 4 principal components (fractional variance: 0.386, 0.103, 0.047, 0.039, respectively) of ESM2-650M embeddings of natural (dark shades) and foldtuning-generated (light shades) proteins for 11 selected SCOP folds – G-protein coupled receptors (GPCRs, SCOP: 2000339), small GTPases (SCOP: 2001260), immunoglobulin *β*-sandwich domains (Ig*β*s, SCOP: 2000051), leucine-rich repeat domains (LRRs, SCOP: 2000631), *ββα*-zinc fingers (SCOP: 2000684), basic helix-loop-helix DNA-binding domains (bHLHs, SCOP: 2001201), defensins (SCOP: 2000440), three-finger toxins (3FTxs, SCOP: 2000373), TIM-*β*/*α* barrels (SCOP: 2000031), SH3 domains (SCOP: 2000090), and barstar-like domains (SCOP: 2000624).

**Figure S2.**
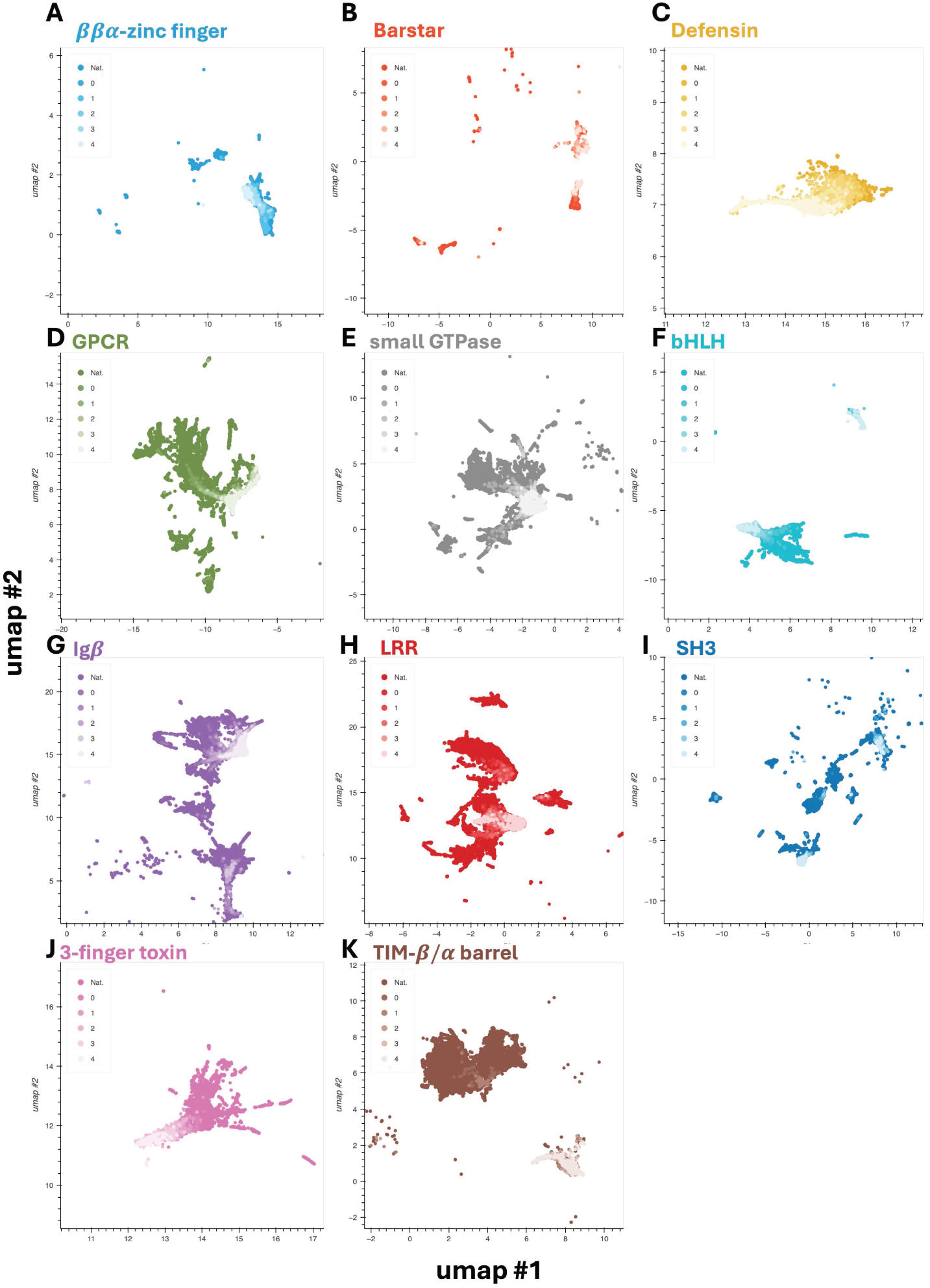
ESM2-650M embeddings capture round-by-round drift of foldtuned sequences from natural counterparts. 2D UMAP representation of ESM2-650M embeddings for eleven representative target fold classes, progressing from natural examples (darkest) through four rounds of foldtuning (lightest). Subfigure boundaries are set to the 5th- and 95th- quantiles in each UMAP component. Selected folds – (A) *ββα*-zinc finger. (B) Barstar. (C) Defensin. (D) G-protein coupled receptor (GPCR). (E) Small GTPase. (F) Basic helix-loop-helix DNA-binding domain (bHLH). (G) Immunoglobulin *β*-sandwich (Ig*β*). (H) Leucine-rich repeat (LRR). (I) SH3 domain. (J) Three-finger toxin domain (3FTx). (K) TIM *β*/*α* barrel.

**Figure S3.**
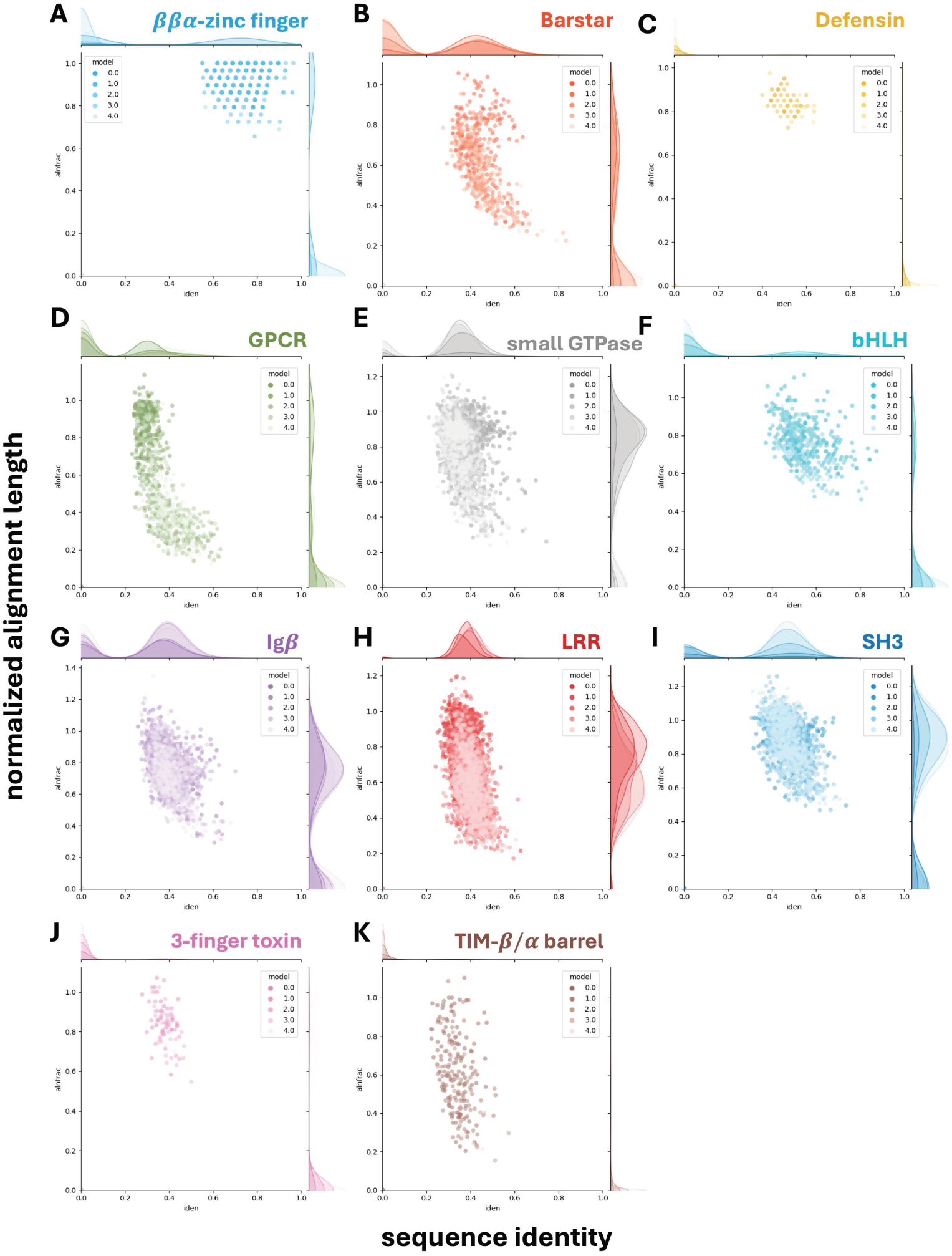
Sequence alignment length and similarity to natural proteins decreases over foldtuning rounds. Plots of normalized alignment length vs. sequence identity over the aligned region for the closest UniRef50 homolog to each foldtuned variant as identified by ultrasensitive search with MMSeqs2. Selected folds – (A) *ββα*-zinc finger. (B) Barstar. (C) Defensin. (D) G-protein coupled receptor (GPCR). (E) Small GTPase. (F) Basic helix-loop-helix DNA-binding domain (bHLH) (G) Immunoglobulin *β*-sandwich (Ig*β*). (H) Leucine-rich repeat (LRR). (I) SH3 domain. (J) Three-finger toxin domain (3FTx). (K) TIM *β*/*α* barrel. 3

**Figure S4.**
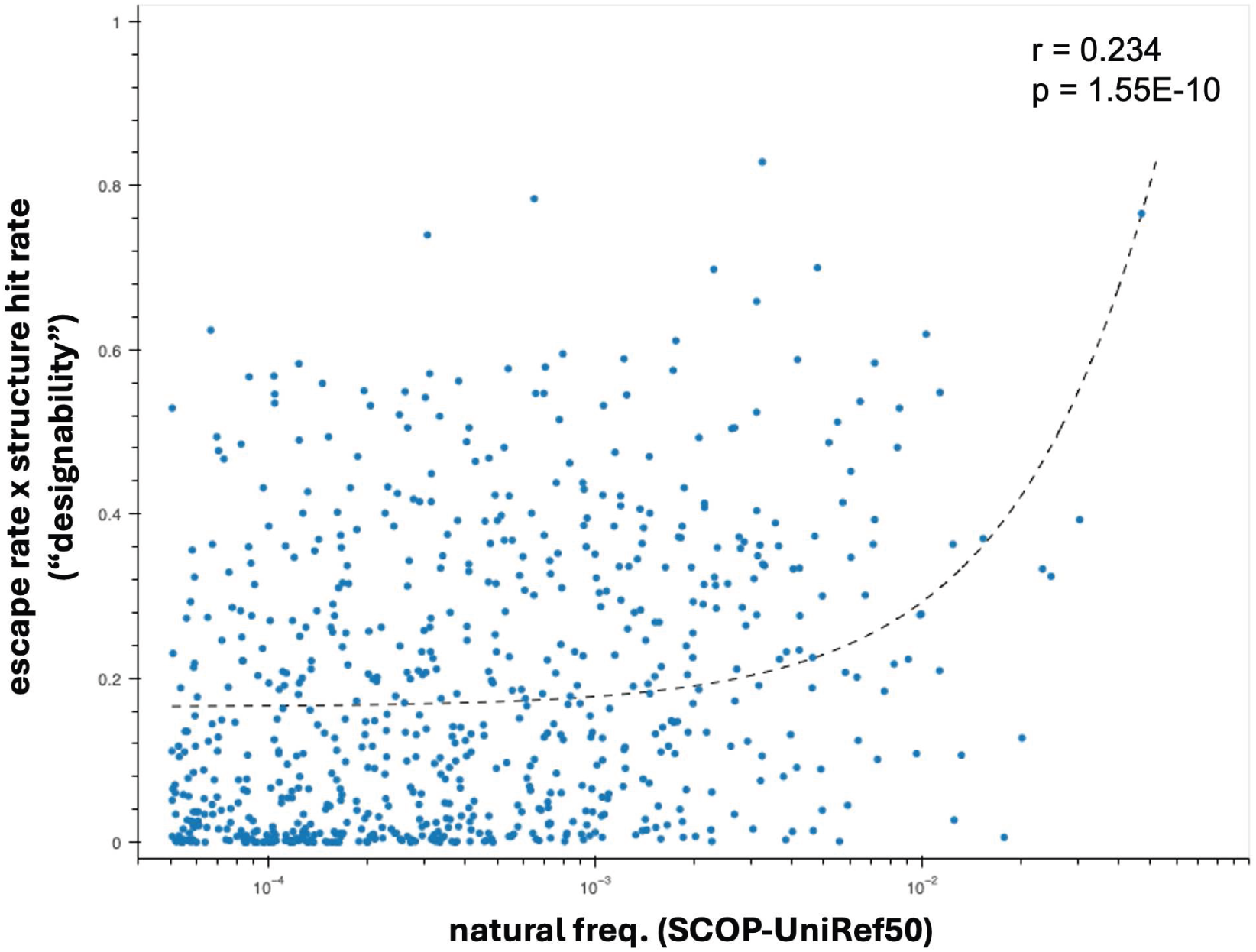
Fold designability is weakly explained by natural frequency. Designability proxy (structural hit rate × sequence escape rate) across n = 708 SCOP fold targets is only weakly explained by rate of occurrence in the custom SCOP-UniRef50 database of natural variants: linear regression t-test for positive slope; slope= 12.80, r = 0.234, p = 1.55 × 10^−10^.

**Figure S5.**
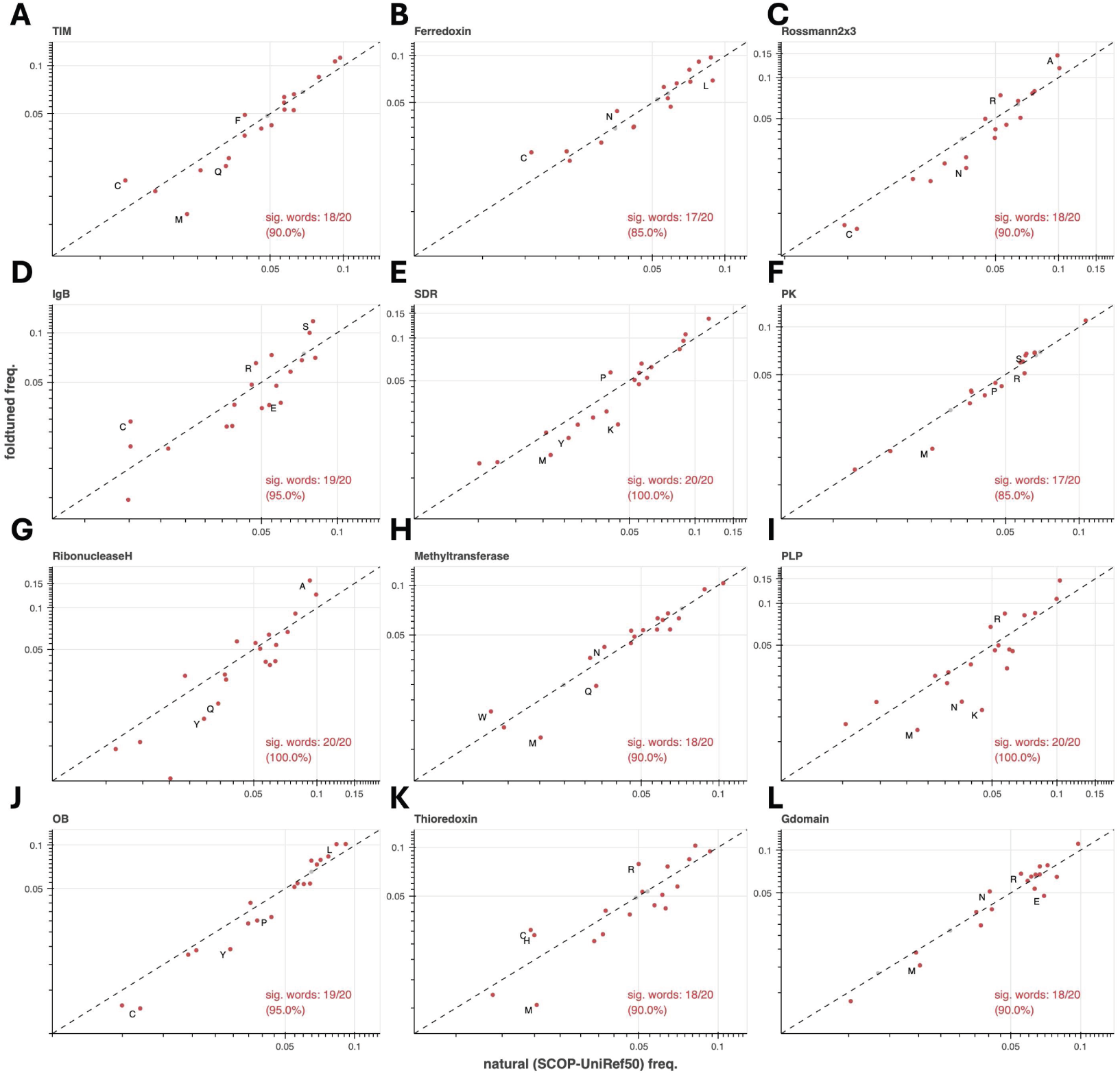
Usage patterns of the 20 canonical amino-acids in foldtuned sequences vs. natural sequences for most-abundant folds. Sig. words denotes the count/fraction of AAs with a statistically significant usage shift (colored red, vs n = 1000 bootstrapped SCOP-UniRef50 replicates, p < 0.05 under Binyamini-Hochberg correction for positively correlated tests). The top-4 most-shifted AAs as ranked by usage fold-change are labeled. Selected folds (top 12 most-abundant in SCOP-UniRef50 database): (A) TIM *β*/*α* barrel (TIM). (B) Ferredoxin (C) Rossmann2x3oid (Rossman2x3). (D) Ig*β*-like (Ig*β*). (E) Short-chain dehydrogenase (SDR). (F) Protein kinase (PK). (G) Ribonuclease H. (H) Methyltransferase. (I) PLP-dependent transferase (PLP). (J) OB fold (OB). (K) Thioredoxin. (L) small GTPase (Gdomain).

**Figure S6.**
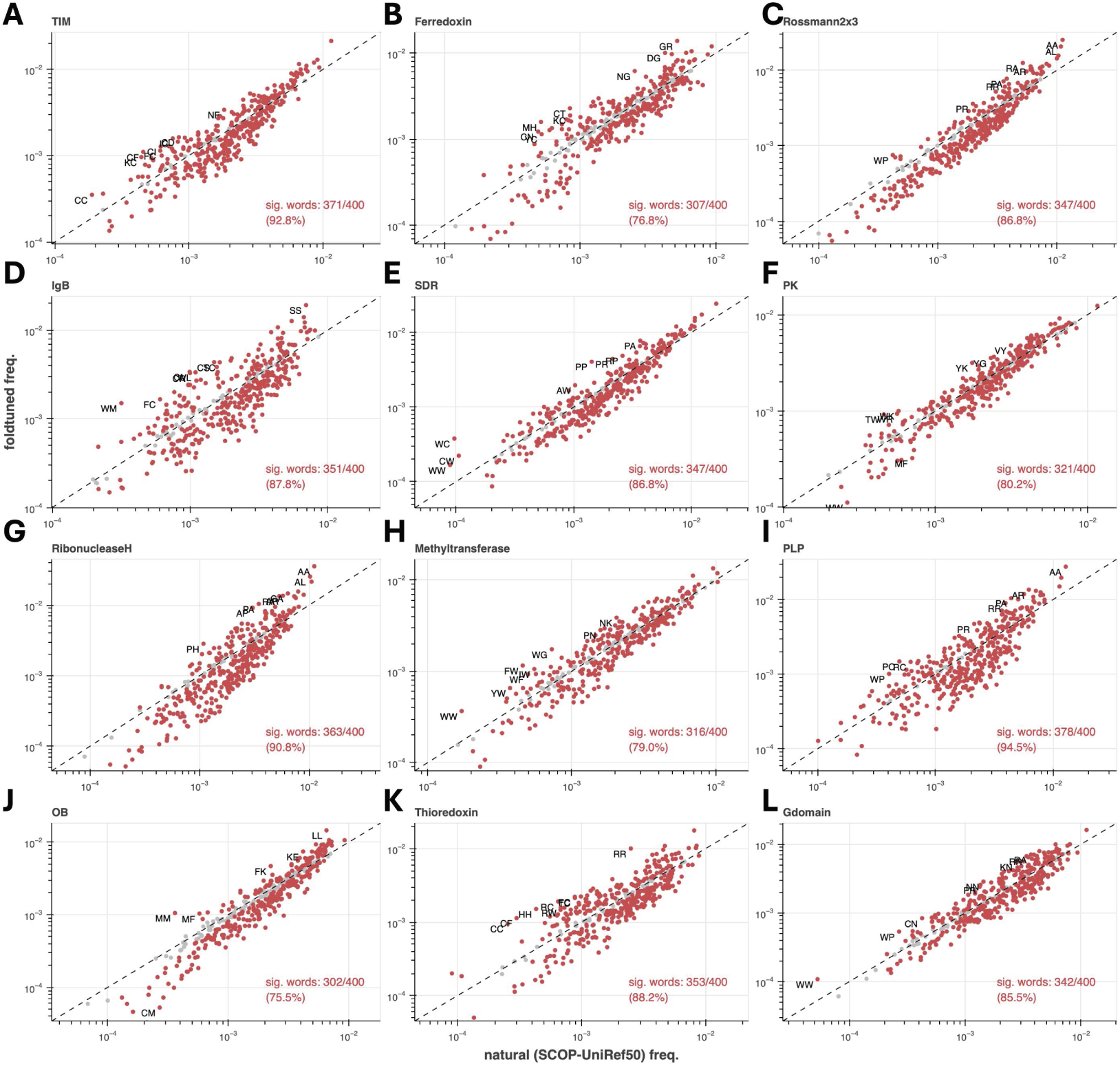
Usage patterns of amino-acid subsequences of length 2 in foldtuned sequences vs. natural sequences for most-abundant folds. Sig. words denotes the count/fraction of bigrams with a statistically significant usage shift (colored red, vs n = 1000 bootstrapped SCOP-UniRef50 replicates, p < 0.05 under Binyamini-Hochberg correction for positively correlated tests). The top-4 most-shifted bigrams as ranked by usage fold-change are labeled. Selected folds (top 12 most-abundant in SCOP-UniRef50 database): (A) TIM *β*/*α* barrel (TIM). (B) Ferredoxin (C) Rossmann2x3oid (Rossman2x3). (D) Ig*β*-like (Ig*β*). (E) Short-chain dehydrogenase (SDR). (F) Protein kinase (PK). (G) Ribonuclease H. (H) Methyltransferase. (I) PLP-dependent transferase (PLP). (J) OB fold (OB). (K) Thioredoxin. (L) small GTPase (Gdomain).

**Figure S7.**
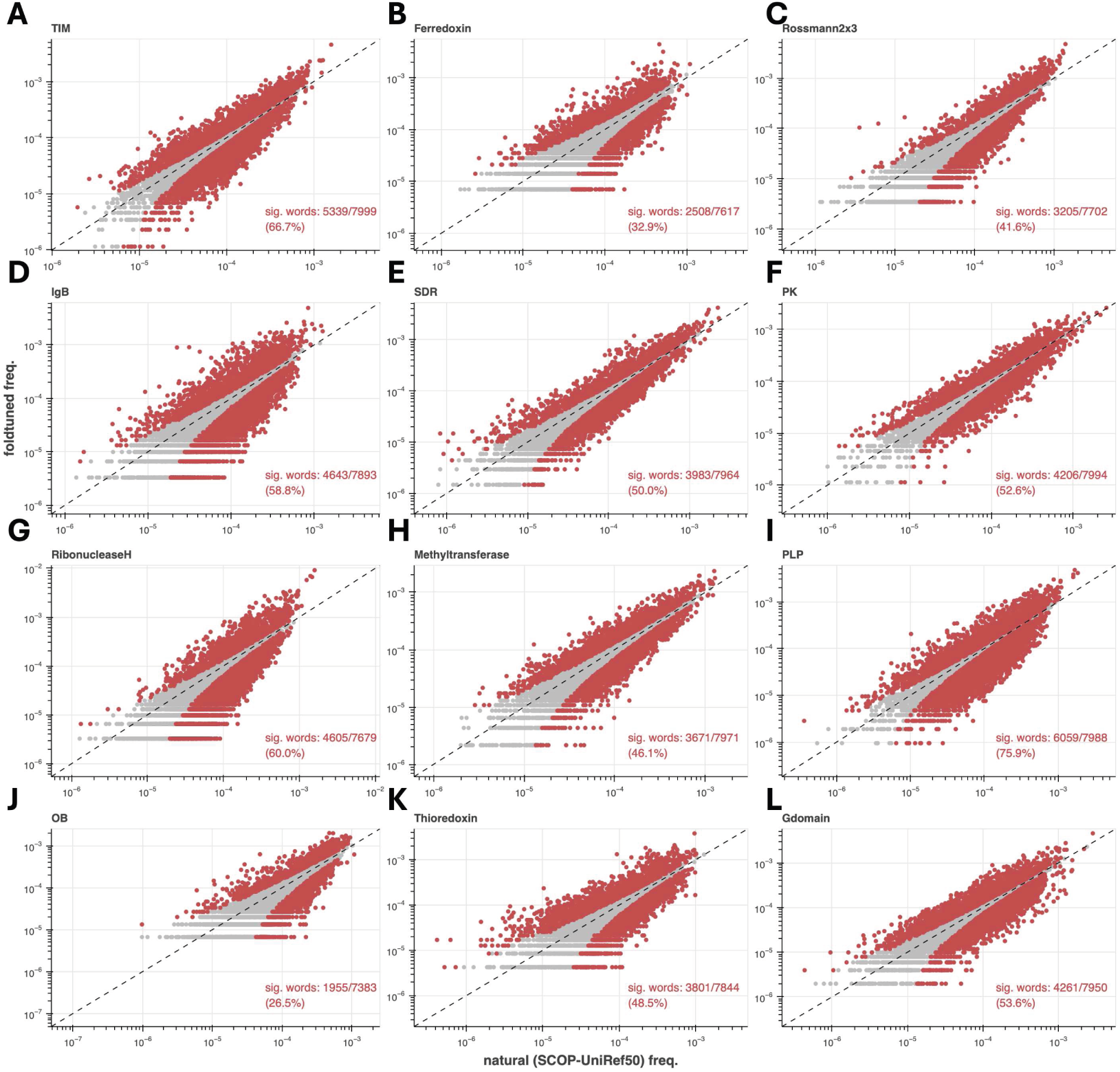
Usage patterns of amino-acid subsequences of length 3 in foldtuned sequences vs. natural sequences for most-abundant folds. Sig. words denotes the count/fraction of trigrams with a statistically significant usage shift (colored red, vs n = 1000 bootstrapped SCOP-UniRef50 replicates, p < 0.05 under Binyamini-Hochberg correction for positively correlated tests). Selected folds (top 12 most-abundant in SCOP-UniRef50 database): (A) TIM *β*/*α* barrel (TIM). (B) Ferredoxin (C) Rossmann2x3oid (Rossman2x3). (D) Ig*β*-like (Ig*β*). (E) Short-chain dehydrogenase (SDR). (F) Protein kinase (PK). (G) Ribonuclease H. (H) Methyltransferase. (I) PLP-dependent transferase (PLP). (J) OB fold (OB). (K) Thioredoxin. (L) small GTPase (Gdomain).

**Figure S8.**
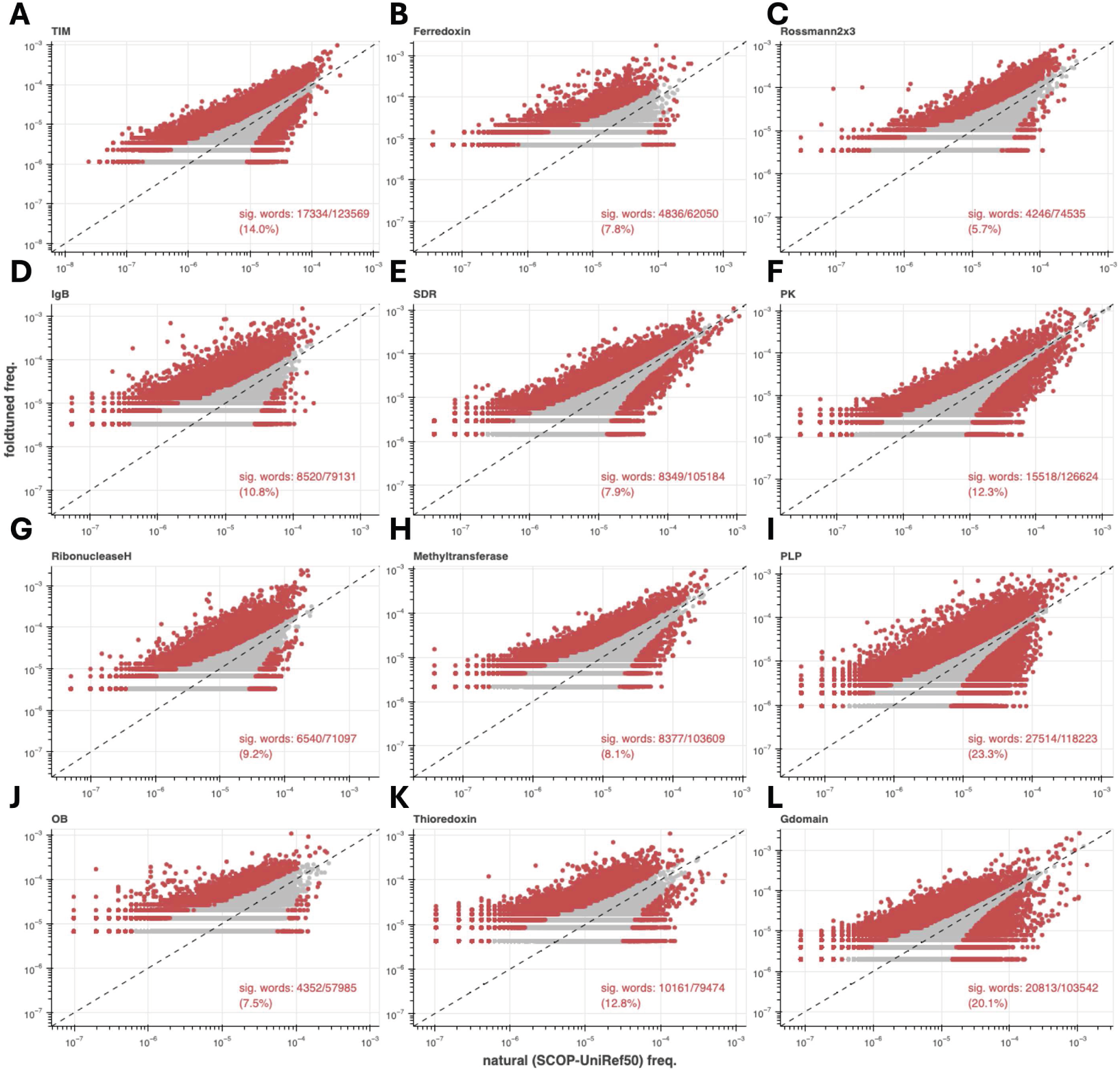
Usage patterns of amino-acid subsequences of length 4 in foldtuned sequences vs. natural sequences for most-abundant folds. Sig. words denotes the count/fraction of quadgrams with a statistically significant usage shift (colored red, vs n = 1000 bootstrapped SCOP-UniRef50 replicates, p < 0.05 under Binyamini-Hochberg correction for positively correlated tests). Selected folds (top 12 most-abundant in SCOP-UniRef50 database): (A) TIM *β*/*α* barrel (TIM). (B) Ferredoxin (C) Rossmann2x3oid (Rossman2x3). (D) Ig*β*-like (Ig*β*). (E) Short-chain dehydrogenase (SDR). (F) Protein kinase (PK). (G) Ribonuclease H. (H) Methyltransferase. (I) PLP-dependent transferase (PLP). (J) OB fold (OB). (K) Thioredoxin. (L) small GTPase (Gdomain).

**Figure S9.**
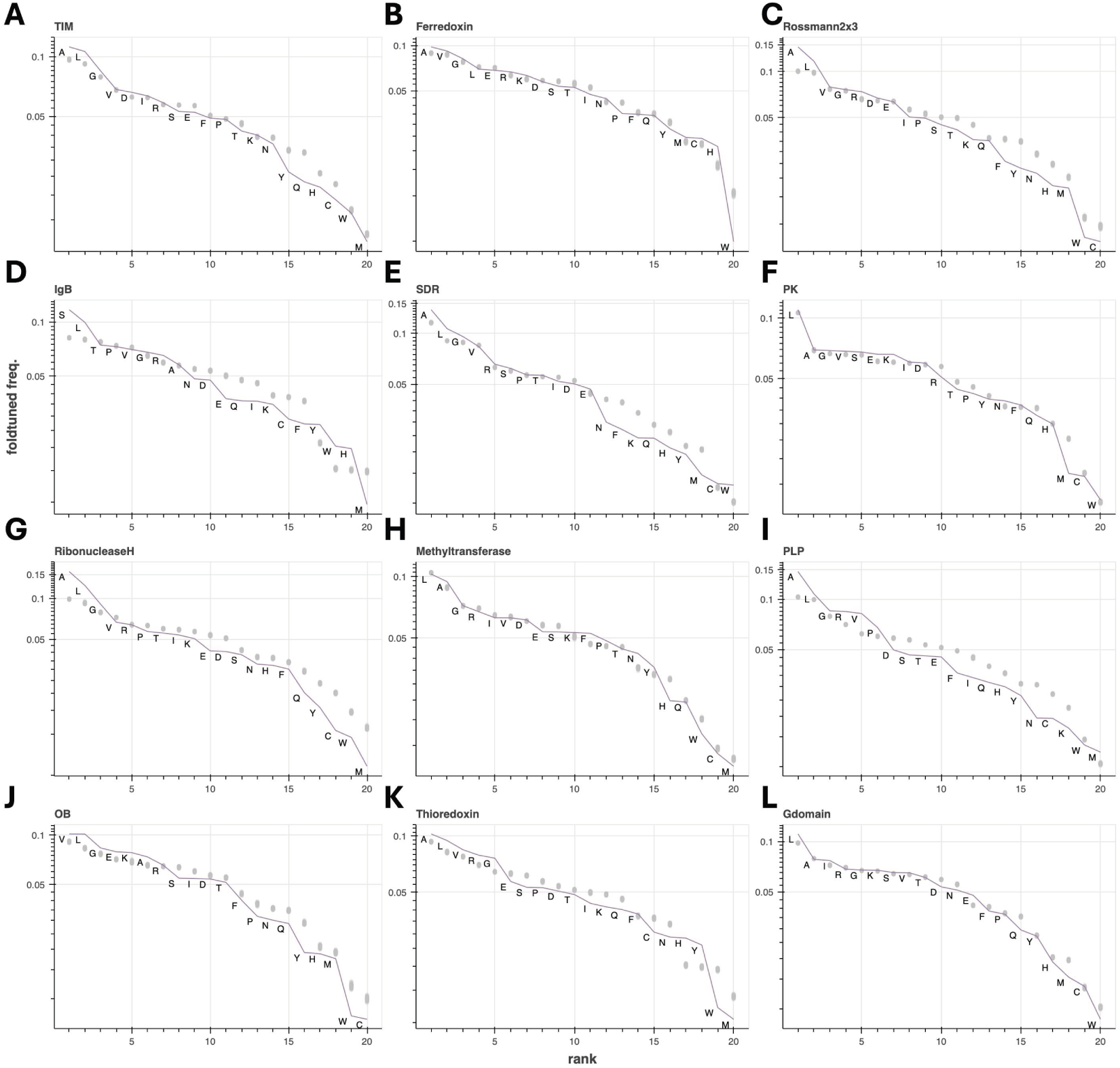
Rank-ordered usage of individual amino-acids in foldtuned sequences relative to natural background for most-abundant folds. Amino-acid usage frequencies for foldtuned variants (purple, labeled) and natural bootstrap samples (gray) are computed as in Figure S5. Selected folds (top 12 most-abundant in SCOP-UniRef50 database): (A) TIM *β*/*α* barrel (TIM). (B) Ferredoxin (C) Rossmann2x3oid (Rossman2x3). (D) Ig*β*-like (Ig*β*). (E) Short-chain dehydrogenase (SDR). (F) Protein kinase (PK). (G) Ribonuclease H. (H) Methyltransferase. (I) PLP-dependent transferase (PLP). (J) OB fold (OB). (K) Thioredoxin. (L) small GTPase (Gdomain).

**Figure S10.**
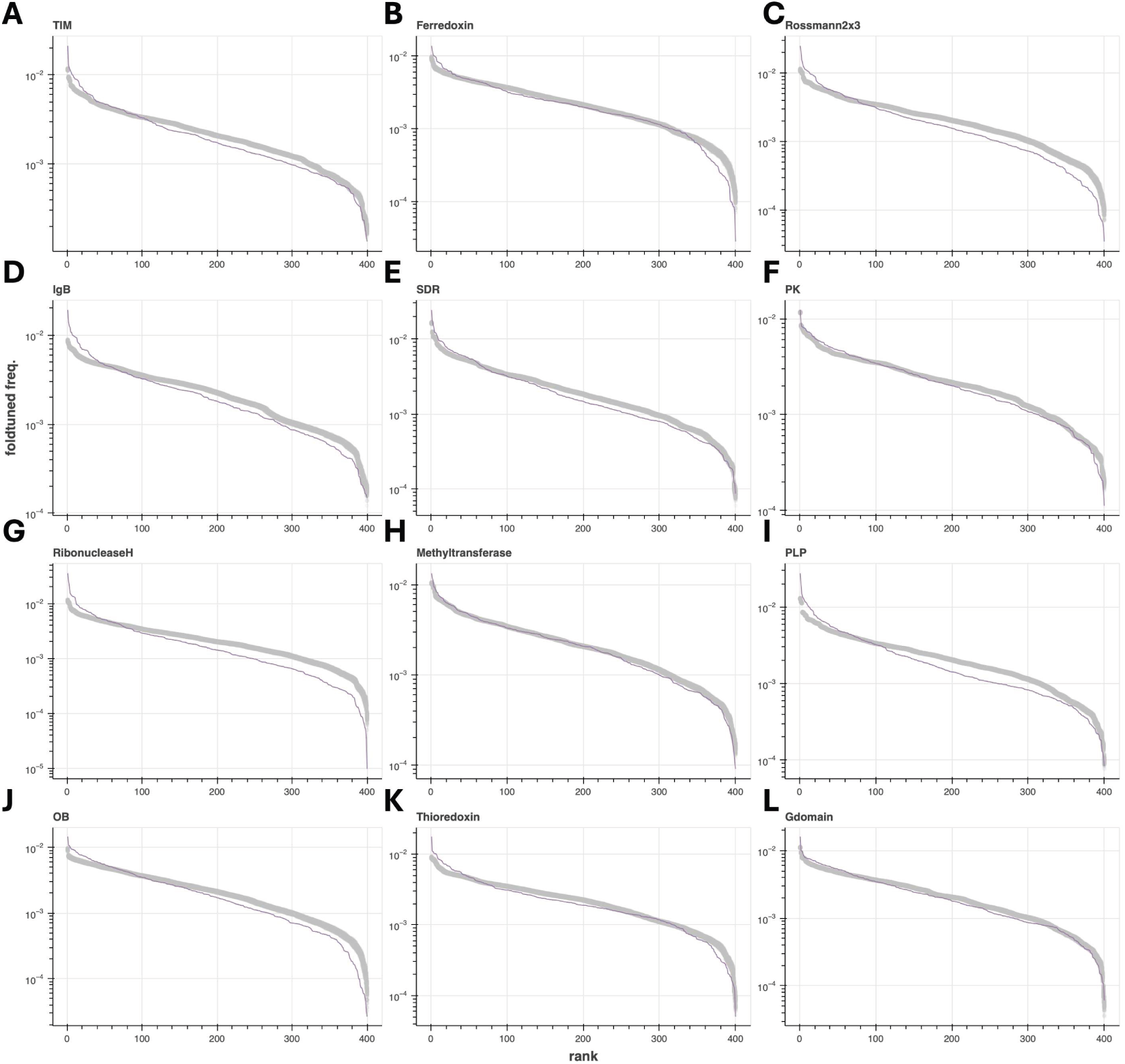
Rank-ordered usage of subsequences of length 2 in foldtuned sequences relative to natural background for most-abundant folds. Bigram frequencies for foldtuned variants (purple, labeled) and natural bootstrap samples (gray) are computed as in Figure S6. Selected folds (top 12 most-abundant in SCOP-UniRef50 database): (A) TIM *β*/*α* barrel (TIM). (B) Ferredoxin (C) Rossmann2x3oid (Rossman2x3). (D) Ig*β*-like (Ig*β*). (E) Short-chain dehydrogenase (SDR). (F) Protein kinase (PK). (G) Ribonuclease H. (H) Methyltransferase. (I) PLP-dependent transferase (PLP). (J) OB fold (OB). (K) Thioredoxin. (L) small GTPase (Gdomain).

**Figure S11.**
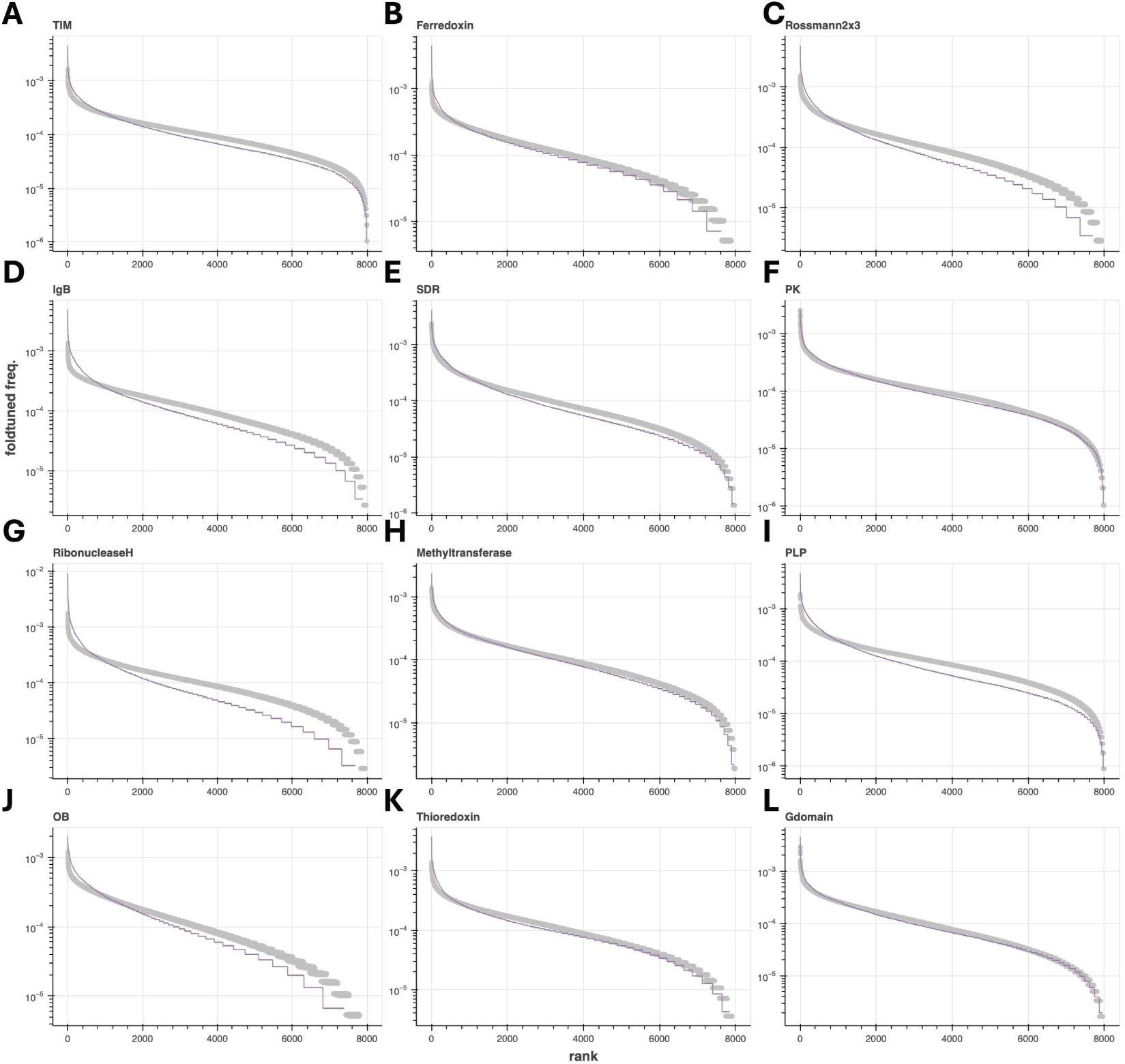
Rank-ordered usage of subsequences of length 3 in foldtuned sequences relative to natural background for most-abundant folds. Trigram frequencies for foldtuned variants (purple, labeled) and natural bootstrap samples (gray) are computed as in Figure S7. Selected folds (top 12 most-abundant in SCOP-UniRef50 database): (A) TIM *β*/*α* barrel (TIM). (B) Ferredoxin (C) Rossmann2x3oid (Rossman2x3). (D) Ig*β*-like (Ig*β*). (E) Short-chain dehydrogenase (SDR). (F) Protein kinase (PK). (G) Ribonuclease H. (H) Methyltransferase. (I) PLP-dependent transferase (PLP). (J) OB fold (OB). (K) Thioredoxin. (L) small GTPase (Gdomain).

**Figure S12.**
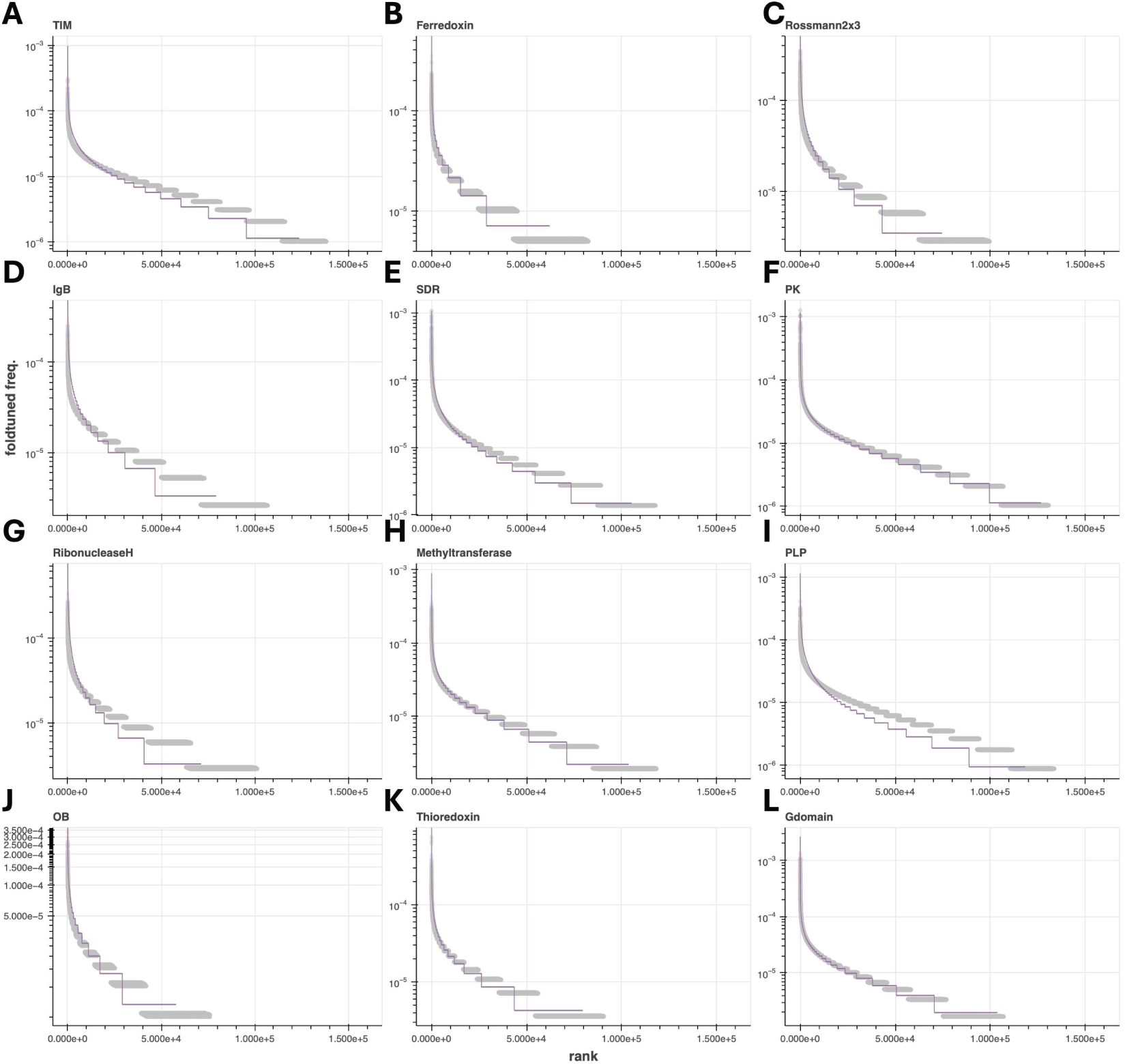
Rank-ordered usage of subsequences of length 4 in foldtuned sequences relative to natural background for most-abundant folds. Quadgram frequencies for foldtuned variants (purple, labeled) and natural bootstrap samples (gray) are computed as in Figure S8. Selected folds (top 12 most-abundant in SCOP-UniRef50 database): (A) TIM *β*/*α* barrel (TIM). (B) Ferredoxin (C) Rossmann2x3oid (Rossman2x3). (D) Ig*β*-like (Ig*β*). (E) Short-chain dehydrogenase (SDR). (F) Protein kinase (PK). (G) Ribonuclease H. (H) Methyltransferase. (I) PLP-dependent transferase (PLP). (J) OB fold (OB). (K) Thioredoxin. (L) small GTPase (Gdomain).

**Figure S13.**
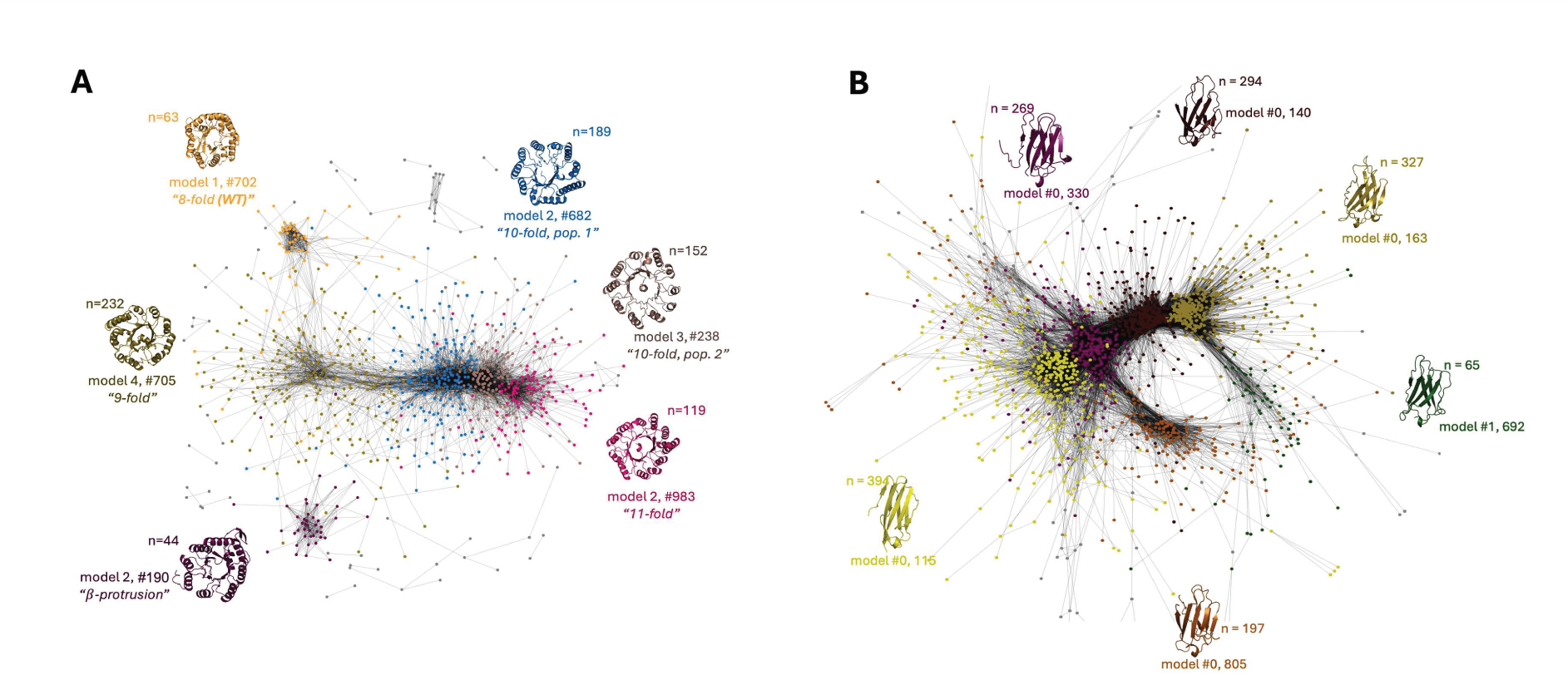
Foldtuned models access new structural innovations without explicit prompting. Network representation of structural similarity among foldtuned variants (linkage threshold: reciprocal TMscore > 0.7; isolated/orphan nodes removed). Nodes are colored by Louvain clustering assignments, for: (A) TIM *α*/*β* barrels (SCOP: 2000031, n = 799). (B) Immunoglobulin*β*-sandwich domains (SCOP 2000051, n = 1546).

**Figure S14.**
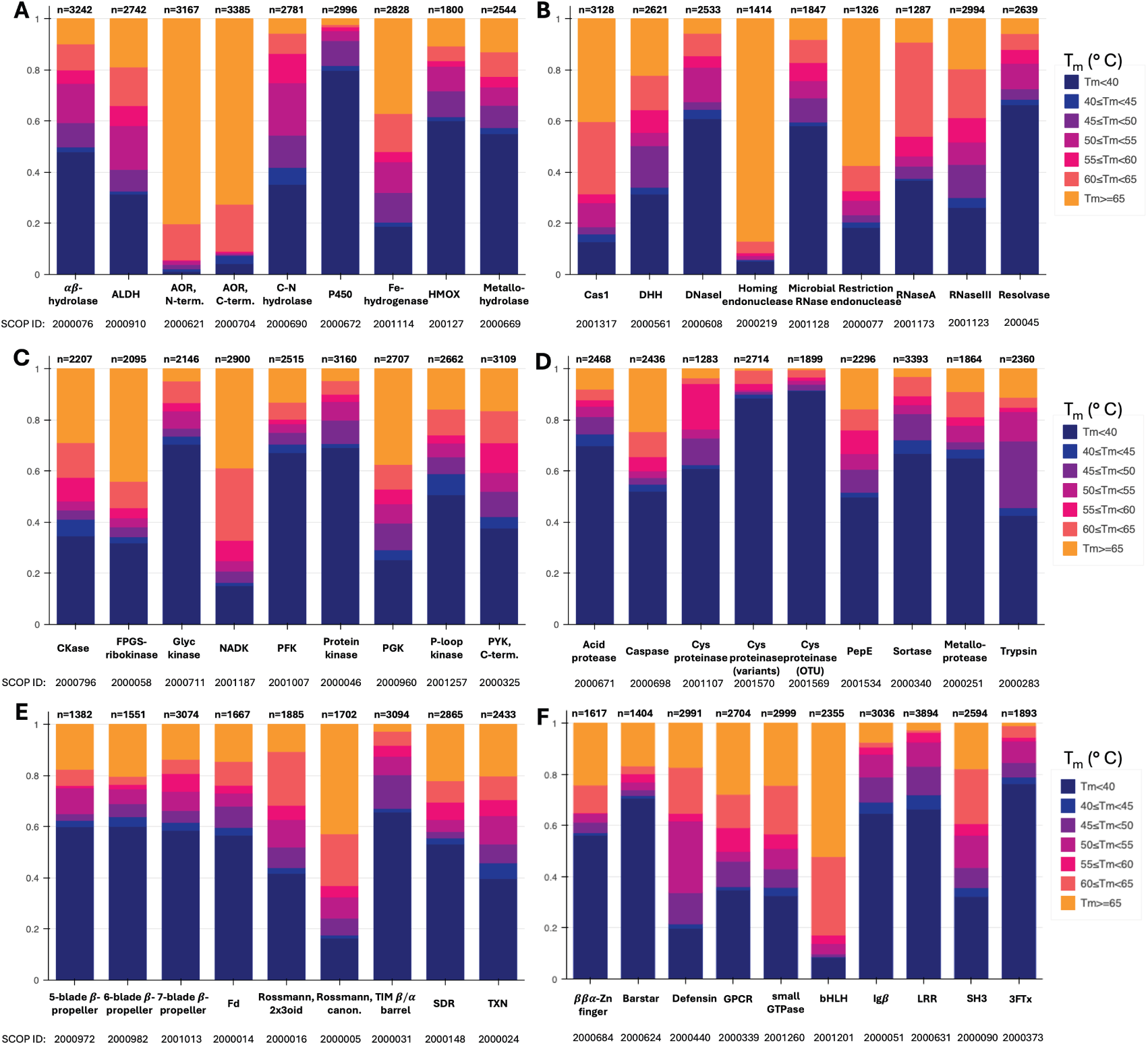
Foldtuned proteins are predicted to exhibit varying degrees of thermostability. Filtered, validated sequences generated from 55 foldtuned models of relevance to protein engineering and synthetic biology are expected to exhibit melting temperatures (T_m_) ranging from < 40^◦^C to > 65^◦^C, as predicted by TemStaPro.^56^ Selected models are grouped into: (A) Hydrolase and oxidoreductase enzymes. (B) Nucleases and other gene-editing-related proteins. (C) Kinases (D) Proteases and peptidases (E) Multi-family enzyme topologies/scaffolds. (F) Common synthetic biology “toolkit” parts for cellular engineering applications.

**Figure S15.**
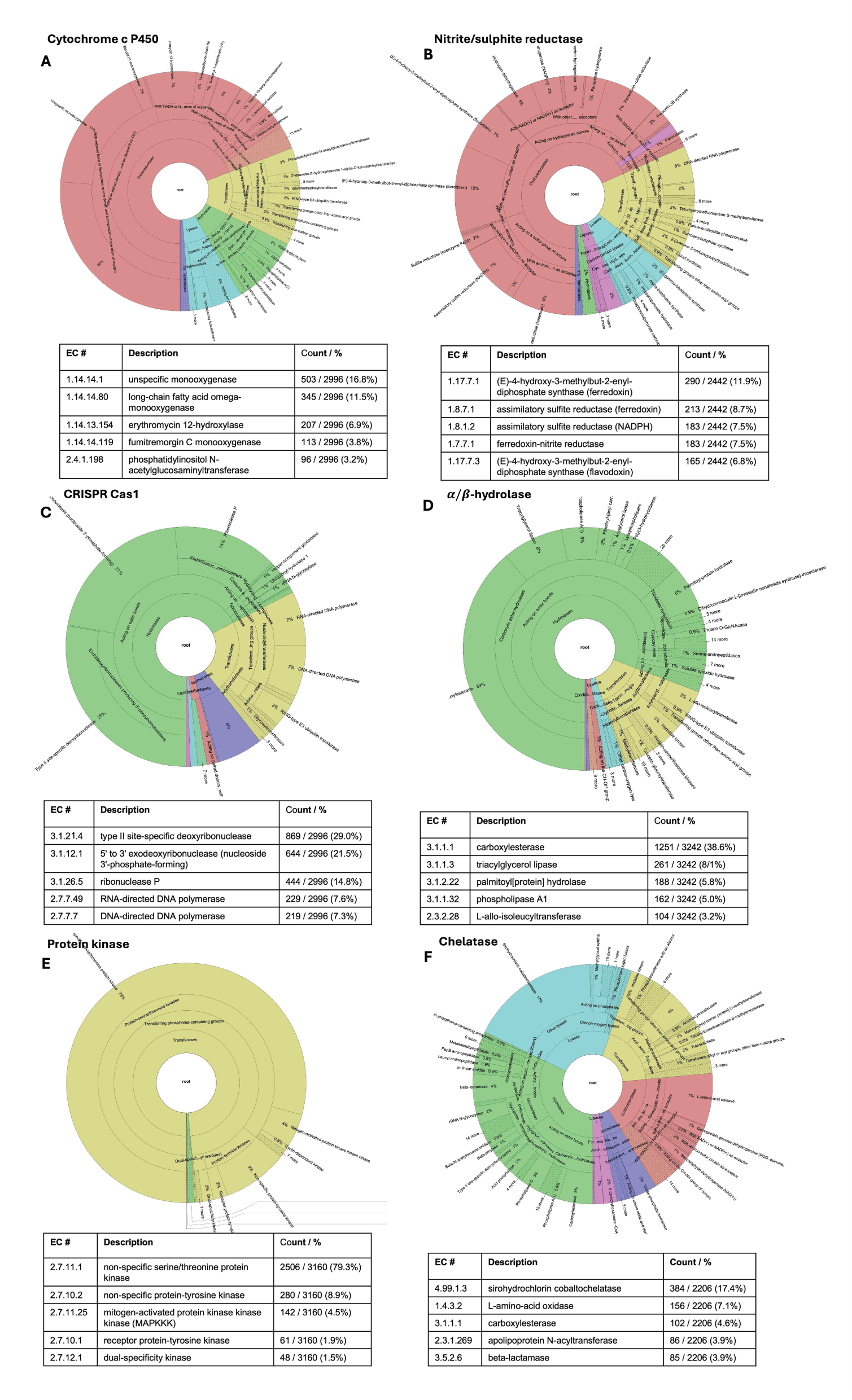
Foldtuned proteins are predicted to mimic or expand parent enzymatic functions. *Top*: Wheel plots of predicted Enzyme Commission (EC) numbers for foldtuned variants of select catalytic folds, as annotated by CLEAN.^57^ Sectors are colored by top-level EC #s – oxidoreductases (EC 1; red), transferases (EC 2; yellow), hydrolases (EC 3; green), lyases (EC 4; blue), isomerases (EC 5; purple), ligases (EC 6; pink). *Bottom*: Descriptions and frequencies of top 5 EC#s annotated per fold. Selected folds: (A) Cytochrome P450s. (B) Nitrite/sulfite reductases. (C) CRISPR Cas1 endonuclease. (D) *α*/*β*-hydrolases. (E) Protein kinases (F) Chelatases

**Figure S16.**
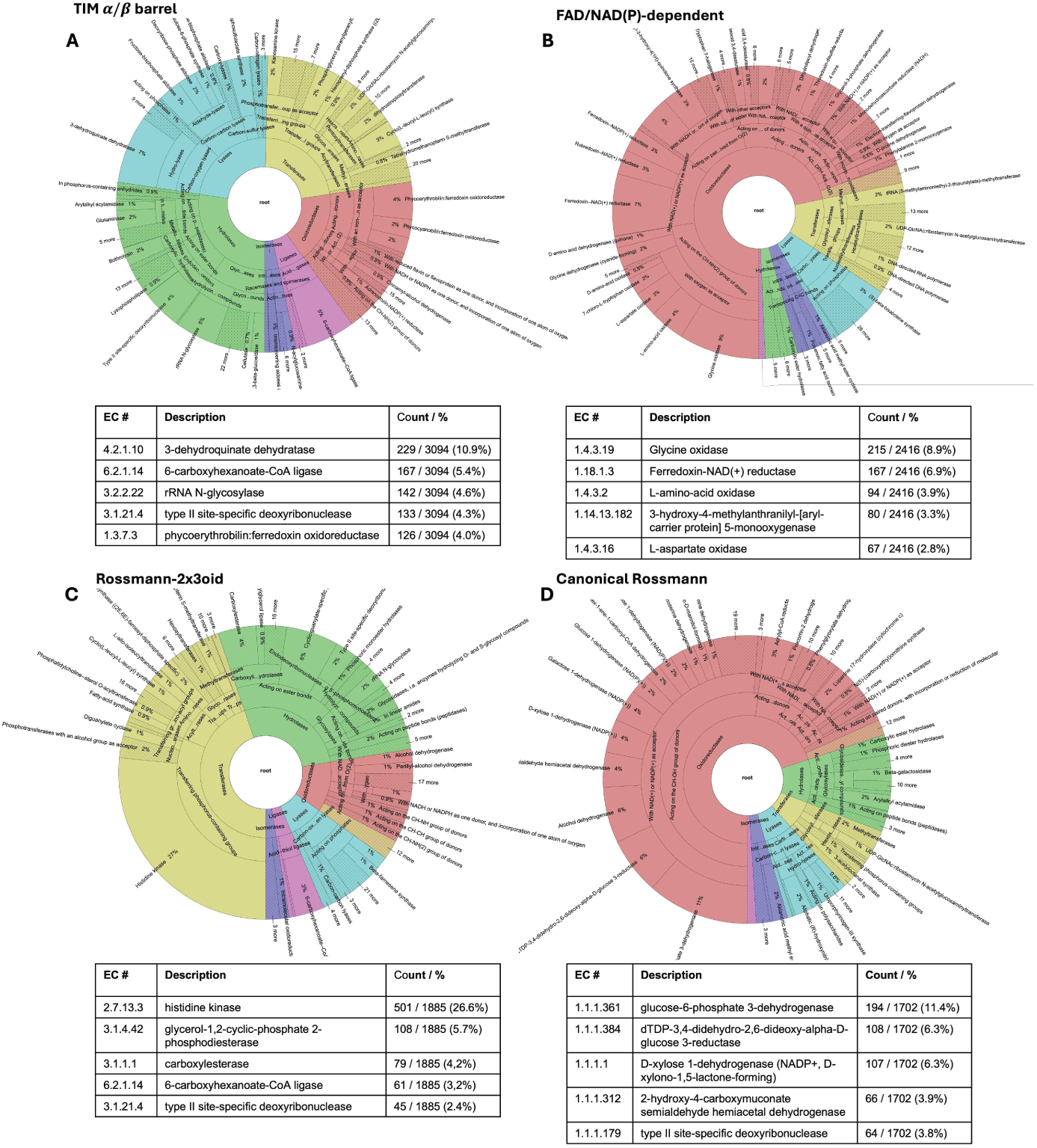
Foldtuned proteins for common enzyme scaffolds are predicted to span wide functional classes. *Top*: Wheel plots of predicted Enzyme Commission (EC) numbers for foldtuned variants of select commonly-utilized enzyme scaffold motifs, as annotated by CLEAN.^57^ Sector coloring follows Figure S15. *Bottom*: Descriptions and frequencies of top 5 EC#s annotated per fold. Selected folds: (A) TIM *β*/*α* barrels. (B) FAD/NAD(P)-dependent enzymes. (C) Rossmann 2x3oid proteins. (D) Canonical Rossmann proteins.

**Figure S17.**
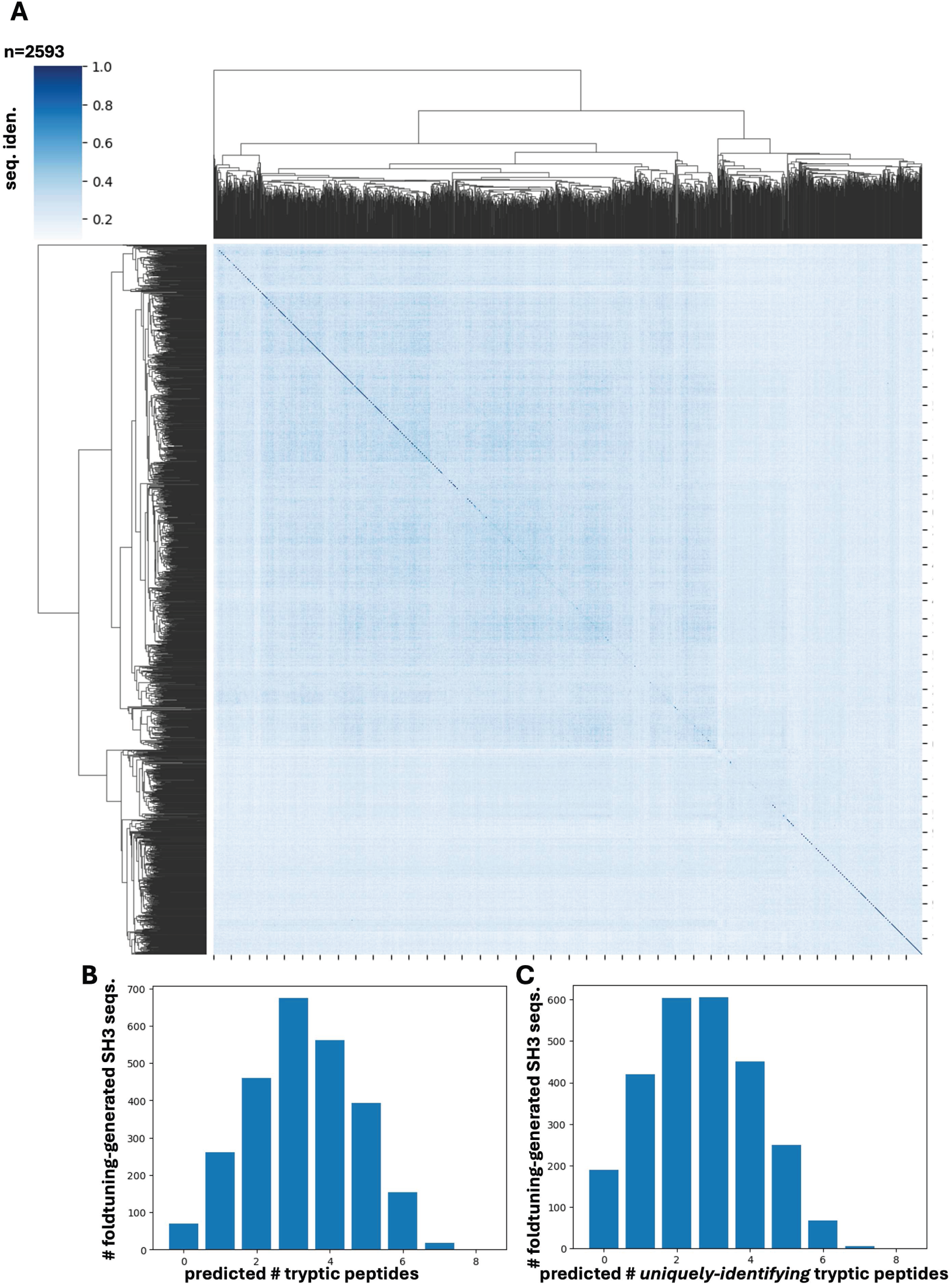
Foldtuned SH3 sequence diversity engenders unique tryptic peptide signatures for mass-spectrometric detection. (A) Hierarchically clustered heatmap of pairwise sequence identity between n = 2593 SH3 domain candidate sequences generated via foldtuning. (B) Expected detectable peptide counts predicted by *in silico* tryptic digestion. (C) Counts of predicted tryptic peptides that map uniquely to single foldtuned SH3 variants.

**Figure S18.**
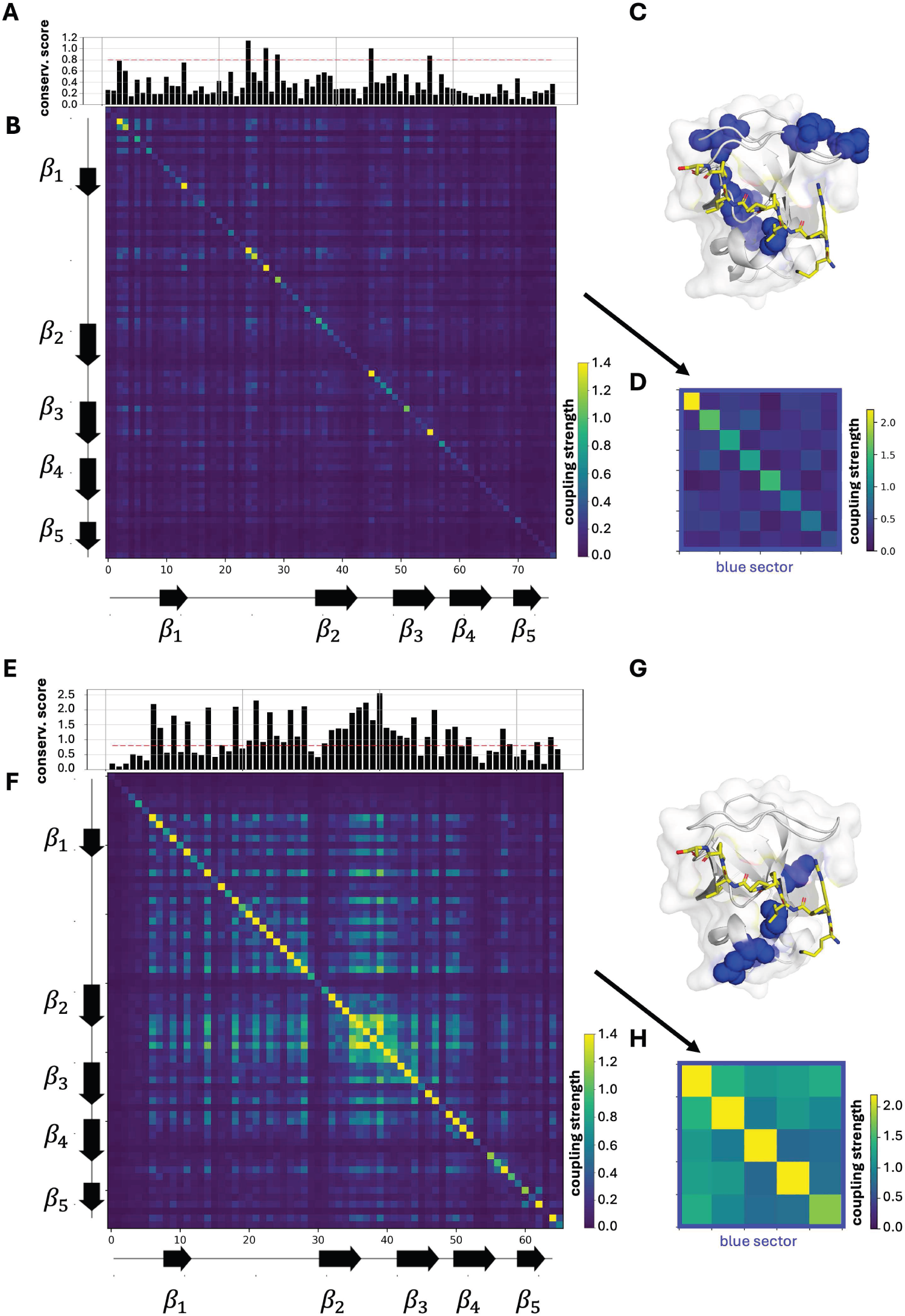
Natural and foldtuned SH3s preserve disjoint structural-functional sectors according to statistical coupling analysis. Results of statistical coupling analysis (SCA) on n ≈ 2500 natural (A)-(D) and n = 2593 foldtuned (E)-(H) SH3 sequences. (A) First-order positional conservation scores (divergence) for natural SH3s. (B) Second-order positional correlation matrix for natural SH3s. (C) Visualization of SCA-identified sectors (blue) mapped onto a representative structure of a natural SH3 domain (from PI3K) bound to a proline-rich peptide ligand (pdb: 3I5R).^97^ (D) Compressed coupling matrix, blocked into a single statistically interacting sector identified by SCA for natural SH3s. (E) First-order positional conservation scores (divergence) for foldtuned SH3s. (F) Second-order positional correlation matrix for foldtuned SH3s. (G) Visualization of SCA-identified sectors (blue) mappe1d8onto a representative structure of a natural SH3 domain (from PI3K) bound to a proline-rich peptide ligand (pdb: 3I5R).^97^ (H) Compressed coupling matrix, blocked into a single statistically interacting sector identified by SCA for foldtuned SH3s.

**Figure S19.**
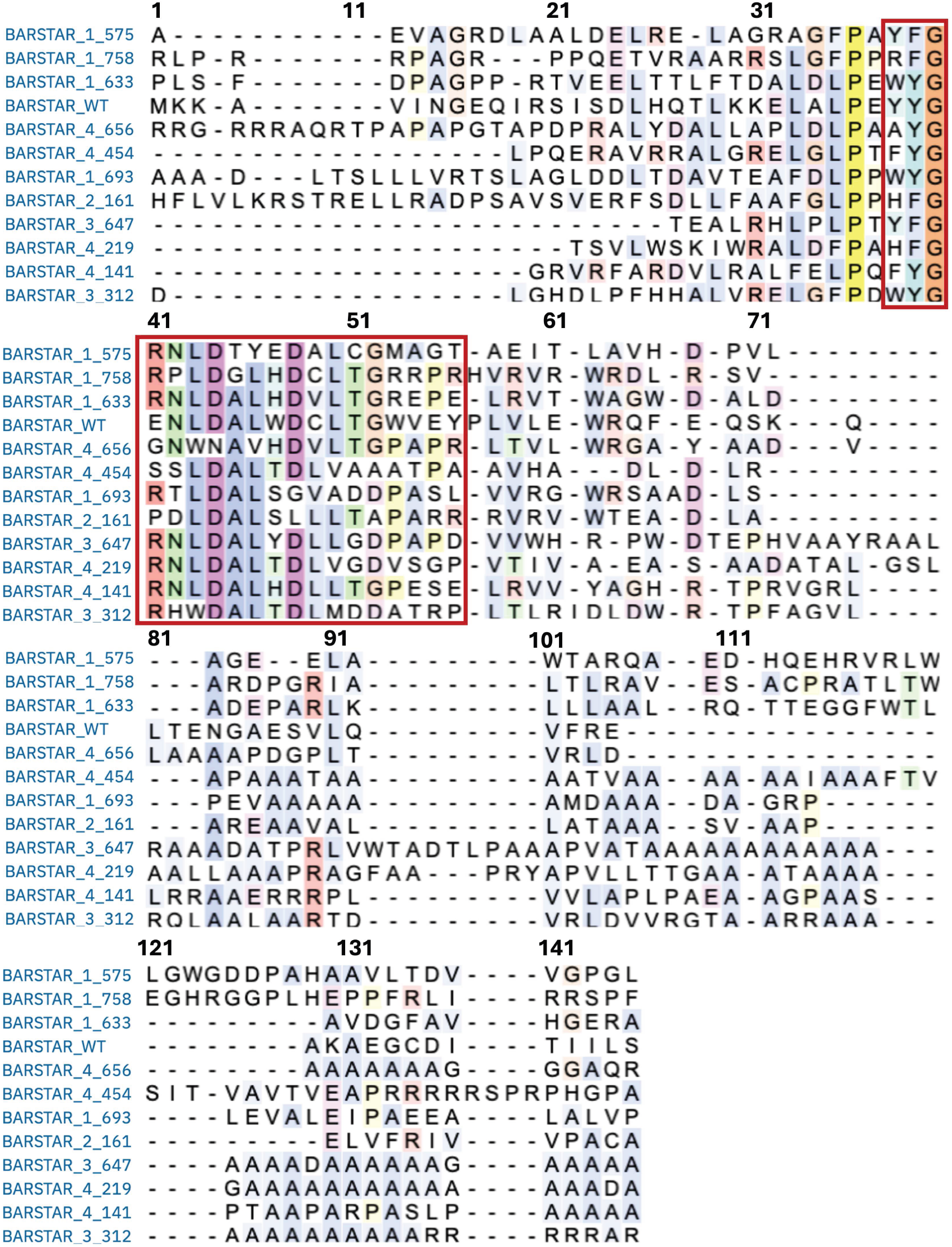
Toxicity-rescuing barstar variants do not uniformly preserve the canonical barnase-binding interface. Muliple sequence alignment (MSA) of the eleven toxicity-rescuing foldtuned barstar variants and wild-type barstar from *B. aquaforiensis*. Columns corresponding to residue positions making physical contacts (positions 38-56; distance threshold < 4.0 Å) with barnase in a reference crystal structure (pdb: 1BRS) are boxed.^72^

## Supplemental Tables

**Table S1.** Foldtuning selectively preserves meaningful sequence context. Comparison of structural hit rates for n = 23 common/characterized SCOP folds for foldtuning (aggregated over four rounds) and controls derived from representative multiple sequence alignments (MSAs). For each fold, control sequences are generated by perturbing natural training examples to retain a local alignment with target sequence identity and fractional length equivalent to that of the median among the subset of foldtuned variants exhibiting detectable homology to a natural counterpart (”non-escaping”).

| SCOP |  |  | Foldtuned |  |  | Rep. Alignment Control |  |  |
| --- | --- | --- | --- | --- | --- | --- | --- | --- |
| ID | Fold | Length (aa) | Struct. Hit Rate (agg.) | Seq. Esc. Rate (agg.) |  | Iden. (median) | Frac. Align. (median) | Struct. Hit Rate |
| 2000631 | Leucine-rich repeat (LRR) | 275 | 0.827 | 0.013 |  | 0.390 | 0.665 | 0.054 |
| 2000076 | $\alpha/\beta$-hydrolase | 277 | 0.726 | 0.226 | | 0.356 | 0.447 | 0.008 |
| 2001260 | G domain-like | 181 | 0.714 | 0.173 |  | 0.356 | 0.845 | 0.029 |
| 2000046 | Protein kinase-like | 293 | 0.707 | 0.004 |  | 0.382 | 0.679 | 0.148 |
| 2000031 | TIM $\beta/\alpha$ barrel | 300 | 0.699 | 0.958 | | 0.338 | 0.563 | 0.009 |
| 2000607 | PLP-dependent transferase-like | 386 | 0.699 | 0.021 |  | 0.383 | 0.720 | 0.025 |
| 2000672 | Cytochrome P450 | 440 | 0.687 | 0.0004 |  | 0.367 | 0.520 | 0.052 |
| 2000051 | Igβ-like | 113 | 0.658 | 0.358 | | 0.391 | 0.743 | 0.012 |
| 2000144 | Winged helix domain | 68 | 0.631 | 0.844 |  | 0.467 | 0.838 | 0.022 |
| 2000148 | SDR-type Rossmann | 252 | 0.626 | 0.087 |  | 0.383 | 0.877 | 0.120 |
| 2000090 | SH3-like barrel | 77 | 0.611 | 0.290 |  | 0.475 | 0.857 | 0.037 |
| 2000440 | Defensin | 40 | 0.605 | 0.984 |  | 0.515 | 0.825 | 0.016 |
| 2001201 | HLH-like | 67 | 0.568 | 0.804 |  | 0.540 | 0.746 | 0.068 |
| 2000339 | GPCR | 295 | 0.550 | 0.718 |  | 0.344 | 0.519 | 0.004 |
| 2000024 | Thioredoxin | 112 | 0.529 | 0.755 |  | 0.388 | 0.688 | 0.014 |
| 2000088 | Methyltransferase-like | 220 | 0.512 | 0.116 |  | 0.371 | 0.545 | 0.008 |
| 2000016 | Rossmann2x3(oid) | 163 | 0.438 | 0.395 |  | 0.410 | 0.810 | 0.019 |
| 2000067 | Ribonuclease H-like | 174 | 0.433 | 0.616 |  | 0.397 | 0.862 | 0.032 |
| 2000373 | Snake toxin-like | 84 | 0.418 | 0.957 |  | 0.379 | 0.845 | 0.001 |
| 2000014 | Ferredoxin-like | 102 | 0.394 | 0.682 |  | 0.425 | 0.706 | 0.008 |
| 2000100 | OB-fold | 104 | 0.388 | 0.820 |  | 0.420 | 0.817 | 0.017 |
| 2000684 | $\beta\beta\alpha$-zinc finger | 29 | 0.376 | 0.752 | | 0.720 | 0.897 | 0.270 |
| 2000624 | Barstar-like | 107 | 0.312 | 0.602 |  | 0.440 | 0.617 | 0.002 |

**Table S2.**
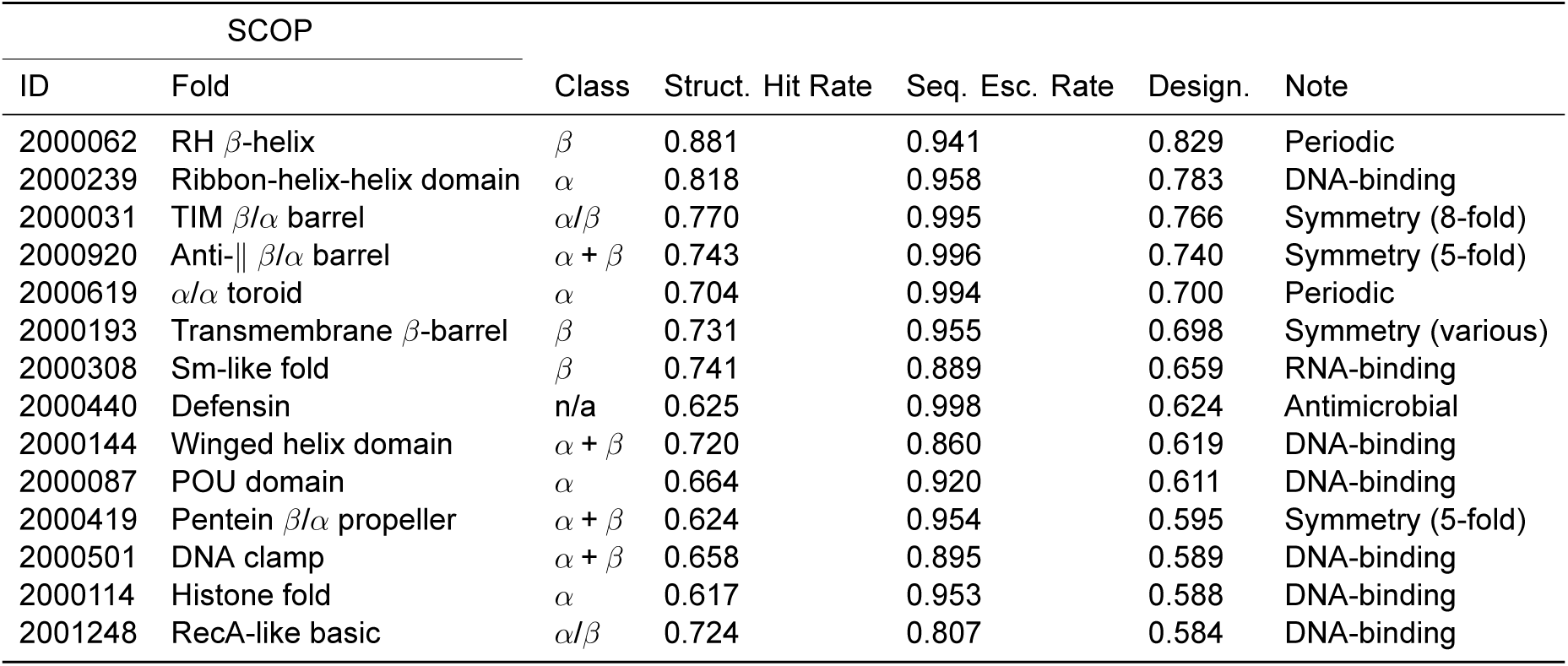
Designability of SCOP folds. Top 2% (n = 14) of succesfully foldtuned SCOP folds (N = 727), ranked by designability proxy (structural hit rate × sequence escape rate), with topology class and structural/functional annotations.

## References

1. Maynard Smith, J. (1970). Natural Selection and the Concept of a Protein Space. en. Nature 225. Publisher: Nature Publishing Group, 563–564. 10.1038/225563a0.

2. Mirdita, M., Schütze, K., Moriwaki, Y., Heo, L., Ovchinnikov, S., and Steinegger, M. (2022). ColabFold: making protein folding accessible to all. en. Nat Methods 19. Publisher: Nature Publishing Group, 679–682. 10.1038/s41592-022-01488-1.

3. Jumper, J., Evans, R., Pritzel, A., Green, T., Figurnov, M., Ronneberger, O., Tunyasuvunakool, K., Bates, R., Žídek, A., Potapenko, A., et al. (2021). Highly accurate protein structure prediction with AlphaFold. en. Nature 596. Number: 7873 Publisher: Nature Publishing Group, 583–589. 10.1038/s41586-021-03819-2.

4. Baker, D. (2000). A surprising simplicity to protein folding. en. Nature 405. Number: 6782 Publisher: Nature Publishing Group, 39–42. 10.1038/35011000.

5. Watters, A. L., Deka, P., Corrent, C., Callender, D., Varani, G., Sosnick, T., and Baker, D. (2007). The Highly Cooperative Folding of Small Naturally Occurring Proteins Is Likely the Result of Natural Selection. English. Cell 128. Publisher: Elsevier, 613–624. 10.1016/j.cell.2006.12.042.

6. Dupont, C. L., Butcher, A., Valas, R. E., Bourne, P. E., and Caetano-Anollés, G. (2010). History of biological metal utilization inferred through phylogenomic analysis of protein structures. Proceedings of the National Academy of Sciences 107. Publisher: Proceedings of the National Academy of Sciences, 10567–10572. 10.1073/pnas.0912491107.

7. Alva, V., Söding, J., and Lupas, A. N. (2015). A vocabulary of ancient peptides at the origin of folded proteins. eLife 4. Ed. by J. Kuriyan. Publisher: eLife Sciences Publications, Ltd, e09410. 10.7554/eLife.09410.

8. Vyas, P., Trofimyuk, O., Longo, L. M., Deshmukh, F. K., Sharon, M., and Tawfik, D. S. (2021). Helicase-like functions in phosphate loop containing beta-alpha polypeptides. Proceedings of the National Academy of Sciences 118. Publisher: Proceedings of the National Academy of Sciences, e2016131118. 10.1073/pnas.2016131118.

9. Yue, K. and Dill, K. A. (1995). Forces of tertiary structural organization in globular proteins. Proceedings of the National Academy of Sciences 92. Publisher: Proceedings of the National Academy of Sciences, 146–150. 10.1073/pnas.92.1.146.

10. Bornberg-Bauer, E. (1997). How are model protein structures distributed in sequence space? en. Biophysical Journal 73, 2393–2403. 10.1016/S0006-3495(97)78268-7.

11. Li, H., Helling, R., Tang, C., and Wingreen, N. (1996). Emergence of Preferred Structures in a Simple Model of Protein Folding. Science 273. Publisher: American Association for the Advancement of Science, 666–669. 10.1126/science.273.5275.666.

12. Helling, R., Li, H., Mélin, R., Miller, J., Wingreen, N., Zeng, C., and Tang, C. (2001). The designability of protein structures. Journal of Molecular Graphics and Modelling 19, 157–167. 10.1016/S1093-3263(00)00137-6.

13. Ho, S. P. and DeGrado, W. F. (1987). Design of a 4-helix bundle protein: synthesis of peptides which self-associate into a helical protein. en. J. Am. Chem. Soc. 109, 6751–6758. 10.1021/ja00256a032.

14. Regan, L. and DeGrado, W. F. (1988). Characterization of a Helical Protein Designed from First Principles. Science 241. Publisher: American Association for the Advancement of Science, 976–978. 10.1126/science.3043666.

15. Kamtekar, S., Schiffer, J. M., Xiong, H., Babik, J. M., and Hecht, M. H. (1993). Protein Design by Binary Patterning of Polar and Nonpolar Amino Acids. Science 262. Publisher: American Association for the Advancement of Science, 1680–1685. 10.1126/science.8259512.

16. Hecht, M. H., Vogel, K. M., Spiro, T. G., Rojas, N. R. L., Kamtekar, S., Simons, C. T., Mclean, J. E., and Farid, R. S. (1997). De novo heme proteins from designed combinatorial libraries. en. Protein Science 6. _eprint: https://onlinelibrary.wiley.com/doi/pdf/10.1002/pro.5560061204, 2524. 10.1002/pro.5560061204.

17. Socolich, M., Lockless, S. W., Russ, W. P., Lee, H., Gardner, K. H., and Ranganathan, R. (2005). Evolutionary information for specifying a protein fold. en. Nature 437. Publisher: Nature Publishing Group, 512–518. 10.1038/nature03991.

18. Lockless, S. W. and Ranganathan, R. (1999). Evolutionarily Conserved Pathways of Energetic Connectivity in Protein Families. Science 286. Publisher: American Association for the Advancement of Science, 295–299. 10.1126/science.286.5438.295.

19. Keefe, A. D. and Szostak, J. W. (2001). Functional proteins from a random-sequence library. en. Nature 410. Number: 6829 Publisher: Nature Publishing Group, 715–718. 10.1038/35070613.

20. Tong, C. L., Lee, K.-H., and Seelig, B. (2021). *De novo* proteins from random sequences through *in vitro* evolution. Current Opinion in Structural Biology. Protein-Carbohydrate Complexes and Glycosylation ● Sequences and Topology 68, 129–134. 10.1016/j.sbi.2020.12.014.

21. Romero, P. A. and Arnold, F. H. (2009). Exploring protein fitness landscapes by directed evolution. en. Nat Rev Mol Cell Biol 10. Publisher: Nature Publishing Group, 866–876. 10.1038/nrm2805.

22. Wilson, A. E., Kosater, W. M., and Liberles, D. A. (2020). Evolutionary Processes and Biophysical Mechanisms: Revisiting Why Evolved Proteins Are Marginally Stable. en. J Mol Evol 88, 415–417. 10.1007/s00239-020-09948-y.

23. Fahlberg, S. A., Freschlin, C. R., Heinzelman, P., and Romero, P. A. (2023). *Neural network extrapolation to distant regions of the protein fitness landscape*. en. Pages: 2023.11.08.566287 Section: New Results. 10.1101/2023.11.08.566287.

24. Hsu, C., Verkuil, R., Liu, J., Lin, Z., Hie, B., Sercu, T., Lerer, A., and Rives, A. (2022). “Learning inverse folding from millions of predicted structures”. en. Proceedings of the 39th International Conference on Machine Learning. ISSN: 2640-3498. PMLR, 8946–8970.

25. Dauparas, J., Anishchenko, I., Bennett, N., Bai, H., Ragotte, R. J., Milles, L. F., Wicky, B. I. M., Courbet, A., Haas, R. J. de, Bethel, N., et al. (2022). Robust deep learning–based protein sequence design using ProteinMPNN. Science 378. Publisher: American Association for the Advancement of Science, 49–56. 10.1126/science.add2187.

26. Tóth-Petróczy, Á. and Tawfik, D. S. (2014). The robustness and innovability of protein folds. Current Opinion in Structural Biology. New constructs and expression of proteins/Sequences and topology 26, 131–138. 10.1016/j.sbi.2014.06.007.

27. Pan, X., Thompson, M. C., Zhang, Y., Liu, L., Fraser, J. S., Kelly, M. J. S., and Kortemme, T. (2020). Expanding the space of protein geometries by computational design of de novo fold families. Science 369. Publisher: American Association for the Advancement of Science, 1132–1136. 10.1126/science.abc0881.

28. Lin, Z., Akin, H., Rao, R., Hie, B., Zhu, Z., Lu, W., Smetanin, N., Verkuil, R., Kabeli, O., Shmueli, Y., et al. (2023). Evolutionary-scale prediction of atomic-level protein structure with a language model. Science 379. Publisher: American Association for the Advancement of Science, 1123–1130. 10.1126/science.ade2574.

29. Verkuil, R., Kabeli, O., Du, Y., Wicky, B. I. M., Milles, L. F., Dauparas, J., Baker, D., Ovchin-nikov, S., Sercu, T., and Rives, A. (2022). *Language models generalize beyond natural proteins*. en. Tech. rep. Section: New Results Type: article. bioRxiv, 2022.12.21.521521. 10.1101/2022.12.21.521521.

30. Zhang, Z., Wayment-Steele, H. K., Brixi, G., Wang, H., Kern, D., and Ovchinnikov, S. (2024). Protein language models learn evolutionary statistics of interacting sequence motifs. Proceedings of the National Academy of Sciences 121. Publisher: Proceedings of the National Academy of Sciences, e2406285121. 10.1073/pnas.2406285121.

31. Ferruz, N., Schmidt, S., and Höcker, B. (2022). ProtGPT2 is a deep unsupervised language model for protein design. en. Nat Commun 13. Number: 1 Publisher: Nature Publishing Group, 4348. 10.1038/s41467-022-32007-7.

32. Madani, A., Krause, B., Greene, E. R., Subramanian, S., Mohr, B. P., Holton, J. M., Olmos, J. L., Xiong, C., Sun, Z. Z., Socher, R., et al. (2023). Large language models generate functional protein sequences across diverse families. en. Nat Biotechnol. Publisher: Nature Publishing Group, 1–8. 10.1038/s41587-022-01618-2.

33. Nijkamp, E., Ruffolo, J. A., Weinstein, E. N., Naik, N., and Madani, A. (2023). ProGen2: Exploring the boundaries of protein language models. English. cels 14. Publisher: Elsevier, 968–978.e3. 10.1016/j.cels.2023.10.002.

34. Nguyen, E., Poli, M., Durrant, M. G., Kang, B., Katrekar, D., Li, D. B., Bartie, L. J., Thomas, A. W., King, S. H., Brixi, G., et al. (2024). Sequence modeling and design from molecular to genome scale with Evo. Science 386. Publisher: American Association for the Advancement of Science, eado9336. 10.1126/science.ado9336.

35. Goodfellow, I. J., Pouget-Abadie, J., Mirza, M., Xu, B., Warde-Farley, D., Ozair, S., Courville, A., and Bengio, Y. (2014). “Generative Adversarial Nets”. Advances in Neural Information Processing Systems. Vol. 27. Curran Associates, Inc.

36. Chomsky, N. (1959). On certain formal properties of grammars. Information and Control 2, 137–167. 10.1016/S0019-9958(59)90362-6.

37. Hie, B., Zhong, E. D., Berger, B., and Bryson, B. (2021). Learning the language of viral evolution and escape. Science 371. Publisher: American Association for the Advancement of Science, 284–288. 10.1126/science.abd7331.

38. Kempen, M. van, Kim, S. S., Tumescheit, C., Mirdita, M., Lee, J., Gilchrist, C. L. M., Söding, J., and Steinegger, M. (2023). Fast and accurate protein structure search with Fold-seek. en. Nat Biotechnol. Publisher: Nature Publishing Group, 1–4. 10.1038/s41587-023-01773-0.

39. Barrio-Hernandez, I., Yeo, J., Jänes, J., Mirdita, M., Gilchrist, C. L. M., Wein, T., Varadi, M., Velankar, S., Beltrao, P., and Steinegger, M. (2023). Clustering predicted structures at the scale of the known protein universe. en. Nature 622. Number: 7983 Publisher: Nature Publishing Group, 637–645. 10.1038/s41586-023-06510-w.

40. Pavlopoulos, G. A., Baltoumas, F. A., Liu, S., Selvitopi, O., Camargo, A. P., Nayfach, S., Azad, A., Roux, S., Call, L., Ivanova, N. N., et al. (2023). Unraveling the functional dark matter through global metagenomics. en. Nature 622. Number: 7983 Publisher: Nature Publishing Group, 594–602. 10.1038/s41586-023-06583-7.

41. (N.d.). Materials and methods are available as supplementary material.

42. Munsamy, G., Illanes-Vicioso, R., Funcillo, S., Nakou, I. T., Lindner, S., Ayres, G., Sheehan, L. S., Moss, S., Eckhard, U., Lorenz, P., et al. (2024). Conditional language models enable the efficient design of proficient enzymes. en. Pages: 2024.05.03.592223 Section: New Results. 10.1101/2024.05.03.592223.

43. Andreeva, A., Kulesha, E., Gough, J., and Murzin, A. G. (2020). The SCOP database in 2020: expanded classification of representative family and superfamily domains of known protein structures. Nucleic Acids Research 48, D376–D382. 10.1093/nar/gkz1064.

44. Blum, M., Andreeva, A., Florentino, L. C., Chuguransky, S. R., Grego, T., Hobbs, E., Pinto, B. L., Orr, A., Paysan-Lafosse, T., Ponamareva, I., et al. (2025). InterPro: the protein sequence classification resource in 2025. Nucleic Acids Res 53, D444–D456. 10.1093/nar/gkae1082.

45. Subramanian, A. M., Martinez, Z. A., and Thomson, M. (2025). *Pretrained protein language models choose between sequence novelty and structural completeness*. en. ISSN: 2692–8205 Pages: 2025.10.01.679905 Section: New Results. 10.1101/2025.10.01.679905.

46. Haffke, M., Fehlmann, D., Rummel, G., Boivineau, J., Duckely, M., Gommermann, N., Cotesta, S., Sirockin, F., Freuler, F., Littlewood-Evans, A., et al. (2019). Structural basis of species-selective antagonist binding to the succinate receptor. en. Nature 574. Publisher: Nature Publishing Group, 581–585. 10.1038/s41586-019-1663-8.

47. Wester, M. R., Yano, J. K., Schoch, G. A., Yang, C., Griffin, K. J., Stout, C. D., and Johnson, E. F. (2004). The Structure of Human Cytochrome P450 2C9 Complexed with Flurbiprofen at 2.0-Å Resolution*. Journal of Biological Chemistry 279, 35630–35637. 10.1074/jbc.M405427200.

48. Nair, S. K. and Burley, S. K. (2003). X-Ray Structures of Myc-Max and Mad-Max Recognizing DNA: Molecular Bases of Regulation by Proto-Oncogenic Transcription Factors. Cell 112, 193–205. 10.1016/S0092-8674(02)01284-9.

49. Sigrist, C. J. A., Cuche, B. A., Castro, E. de, Coudert, E., Redaschi, N., and Bridge, A. (2025). The PROSITE database for protein families, domains, and sites. Nucleic Acids Res, gkaf1188. 10.1093/nar/gkaf1188.

50. McLean, K. J., Sabri, M., Marshall, K. R., Lawson, R. J., Lewis, D. G., Clift, D., Balding, P. R., Dunford, A. J., Warman, A. J., McVey, J. P., et al. (2005). Biodiversity of cytochrome P450 redox systems. eng. Biochem Soc Trans 33, 796–801. 10.1042/BST0330796.

51. O’Neil, K. T., Hoess, R. H., and DeGrado, W. F. (1990). Design of DNA-Binding Peptides Based on the Leucine Zipper Motif. Science 249. Publisher: American Association for the Advancement of Science, 774–778. 10.1126/science.2389143.

52. Islam, S. M. A., Heil, B. J., Kearney, C. M., and Baker, E. J. (2018). Protein classification using modified n-grams and skip-grams. Bioinformatics 34, 1481–1487. 10.1093/bioinformatics/btx823.

53. Huang, P.-S., Feldmeier, K., Parmeggiani, F., Fernandez Velasco, D. A., Höcker, B., and Baker, D. (2016). De novo design of a four-fold symmetric TIM-barrel protein with atomic-level accuracy. en. Nat Chem Biol 12. Number: 1 Publisher: Nature Publishing Group, 29–34. 10.1038/nchembio.1966.

54. Romero-Romero, S., Costas, M., Silva Manzano, D.-A., Kordes, S., Rojas-Ortega, E., Tapia, C., Guerra, Y., Shanmugaratnam, S., Rodríguez-Romero, A., Baker, D., et al. (2021). The Stability Landscape of de novo TIM Barrels Explored by a Modular Design Approach. en. Journal of Molecular Biology 433, 167153. 10.1016/j.jmb.2021.167153.

55. Alford, R. F., Leaver-Fay, A., Jeliazkov, J. R., O’Meara, M. J., DiMaio, F. P., Park, H., Shapovalov, M. V., Renfrew, P. D., Mulligan, V. K., Kappel, K., et al. (2017). The Rosetta All-Atom Energy Function for Macromolecular Modeling and Design. J. Chem. Theory Comput. 13. Publisher: American Chemical Society, 3031–3048. 10.1021/acs.jctc.7b00125.

56. Pudžiuvelytė, I., Olechnovič, K., Godliauskaite, E., Sermokas, K., Urbaitis, T., Gasiunas, G., and Kazlauskas, D. (2024). TemStaPro: protein thermostability prediction using sequence representations from protein language models. Bioinformatics 40, btae157. 10.1093/bioinformatics/btae157.

57. Yu, T., Cui, H., Li, J. C., Luo, Y., Jiang, G., and Zhao, H. (2023). Enzyme function prediction using contrastive learning. Science 379. Publisher: American Association for the Advancement of Science, 1358–1363. 10.1126/science.adf2465.

58. Kim, D. E., Jensen, D. R., Feldman, D., Tischer, D., Saleem, A., Chow, C. M., Li, X., Carter, L., Milles, L., Nguyen, H., et al. (2023). De novo design of small beta barrel proteins. Proceedings of the National Academy of Sciences 120. Publisher: Proceedings of the National Academy of Sciences, e2207974120. 10.1073/pnas.2207974120.

59. Rocklin, G. J., Chidyausiku, T. M., Goreshnik, I., Ford, A., Houliston, S., Lemak, A., Carter, L., Ravichandran, R., Mulligan, V. K., Chevalier, A., et al. (2017). Global analysis of protein folding using massively parallel design, synthesis, and testing. Science 357. Publisher: American Association for the Advancement of Science, 168–175. 10.1126/science.aan0693.

60. Tsuboyama, K., Dauparas, J., Chen, J., Laine, E., Mohseni Behbahani, Y., Weinstein, J. J., Mangan, N. M., Ovchinnikov, S., and Rocklin, G. J. (2023). Mega-scale experimental analysis of protein folding stability in biology and design. en. Nature. Publisher: Nature Publishing Group, 1–11. 10.1038/s41586-023-06328-6.

61. Mayer, B. J. (2001). SH3 domains: complexity in moderation. J Cell Sci 114, 1253–1263. 10.1242/jcs.114.7.1253.

62. Feng, S., Chen, J. K., Yu, H., Simon, J. A., and Schreiber, S. L. (1994). Two Binding Orientations for Peptides to the Src SH3 Domain: Development of a General Model for SH3-Ligand Interactions. Science 266. Publisher: American Association for the Advancement of Science, 1241–1247. 10.1126/science.7526465.

63. Yu, H., Chen, J. K., Feng, S., Dalgarno, D. C., Brauer, A. W., and Schrelber, S. L. (1994). Structural basis for the binding of proline-rich peptides to SH3 domains. Cell 76, 933–945. 10.1016/0092-8674(94)90367-0.

64. Musacchio, A., Noble, M., Pauptit, R., Wierenga, R., and Saraste, M. (1992). Crystal structure of a Src-homology 3 (SH3) domain. en. Nature 359. Publisher: Nature Publishing Group, 851–855. 10.1038/359851a0.

65. Abramson, J., Adler, J., Dunger, J., Evans, R., Green, T., Pritzel, A., Ronneberger, O., Willmore, L., Ballard, A. J., Bambrick, J., et al. (2024). Accurate structure prediction of biomolecular interactions with AlphaFold 3. en. Nature 630. Publisher: Nature Publishing Group, 493–500. 10.1038/s41586-024-07487-w.

66. Süel, G. M., Lockless, S. W., Wall, M. A., and Ranganathan, R. (2003). Evolutionarily conserved networks of residues mediate allosteric communication in proteins. en. Nat Struct Mol Biol 10. Number: 1 Publisher: Nature Publishing Group, 59–69. 10.1038/nsb881.

67. Halabi, N., Rivoire, O., Leibler, S., and Ranganathan, R. (2009). Protein Sectors: Evolutionary Units of Three-Dimensional Structure. English. Cell 138. Publisher: Elsevier, 774–786. 10.1016/j.cell.2009.07.038.

68. Saksela, K. and Permi, P. (2012). SH3 domain ligand binding: What’s the consensus and where’s the specificity? FEBS Letters. Modular Protein Domains 586, 2609–2614. 10.1016/j.febslet.2012.04.042.

69. Schreiber, G., Buckle, A. M., and Fersht, A. R. (1994). Stability and function: two constraints in the evolution of barstar and other proteins. English. Structure 2. Publisher: Elsevier, 945–951. 10.1016/S0969-2126(94)00096-4.

70. Schreiber, G. and Fersht, A. R. (1995). Energetics of protein-protein interactions: Analysis ofthe Barnase-Barstar interface by single mutations and double mutant cycles. Journal of Molecular Biology 248, 478–486. 10.1016/S0022-2836(95)80064-6.

71. Hartley, R. W. (2001). “[38] - Barnase–Barstar Interaction”. Methods in Enzymology. Ed. by A. W. Nicholson. Vol. 341. Ribonucleases - Part A. Academic Press, 599–611. 10.1016/S0076-6879(01)41179-7.

72. Buckle, A. M., Schreiber, G., and Fersht, A. R. (1994). Protein-protein recognition: Crystal structural analysis of a barnase-barstar complex at 2.0-.ANG. resolution. Biochemistry 33. Publisher: American Chemical Society, 8878–8889. 10.1021/bi00196a004.

73. Urakubo, Y., Ikura, T., and Ito, N. (2008). Crystal structural analysis of protein–protein interactions drastically destabilized by a single mutation. en. Protein Science 17. _eprint: https://onlinelibrary.wiley.com/doi/pdf/10.1110/ps.073322508, 1055–1065. 10.1110/ps.073322508.

74. Cao, L., Coventry, B., Goreshnik, I., Huang, B., Sheffler, W., Park, J. S., Jude, K. M., Marković, I., Kadam, R. U., Verschueren, K. H. G., et al. (2022). Design of protein-binding proteins from the target structure alone. en. Nature 605. Publisher: Nature Publishing Group, 551–560. 10.1038/s41586-022-04654-9.

75. Wang, X., Cardoso, S., Cai, K., Venkatesh, P., Hung, A., Ng, M., Hall, C., Coventry, B., Lee, D. S., Chowhan, R., et al. (2025). Tuning insulin receptor signaling using de novo-designed agonists. English. Molecular Cell 85. Publisher: Elsevier, 4064–4081.e9. 10.1016/j.molcel.2025.09.020.

76. Claeys, I., Simonet, G., Poels, J., Van Loy, T., Vercammen, L., De Loof, A., and Vanden Broeck, J. (2002). Insulin-related peptides and their conserved signal transduction pathway. Peptides. Invertebrate Neuropeptides 23, 807–816. 10.1016/S0196-9781(01)00666-0.

77. Lourenco, A. L., Subramanian, A. M., Spencer, R. K., Miao, J., Anaya, M., Fu, W., Chow, E. D., and Thomson, M. (2025). Protein CREATE enables closed-loop design of de novo synthetic protein binders. en. Pages: 2024.12.20.629847 Section: New Results. 10.1101/2024.12.20.629847.

78. Bornberg-Bauer, E. and Chan, H. S. (1999). Modeling evolutionary landscapes: Mutational stability, topology, and superfunnels in sequence space. Proceedings of the National Academy of Sciences 96. Publisher: Proceedings of the National Academy of Sciences, 10689–10694. 10.1073/pnas.96.19.10689.

79. Watson, J. L., Juergens, D., Bennett, N. R., Trippe, B. L., Yim, J., Eisenach, H. E., Ahern, W., Borst, A. J., Ragotte, R. J., Milles, L. F., et al. (2023). De novo design of protein structure and function with RFdiffusion. en. Nature. Publisher: Nature Publishing Group, 1–3. 10.1038/s41586-023-06415-8.

80. Skinnider, M. A. (2024). Invalid SMILES are beneficial rather than detrimental to chemical language models. en. Nat Mach Intell 6. Publisher: Nature Publishing Group, 437–448. 10.1038/s42256-024-00821-x.

81. Kleinberg, J. and Mullainathan, S. (2024). Language Generation in the Limit. en. Advances in Neural Information Processing Systems 37, 66058–66079.

82. Yang, K. K., Alamdari, S., Lee, A. J., Kaymak-Loveless, K., Char, S., Brixi, G., Domingo-Enrich, C., Wang, C., Lyu, S., Fusi, N., et al. (2025). The Dayhoff Atlas: scaling sequence diversity for improved protein generation. en. ISSN: 2692-8205 Pages: 2025.07.21.665991 Section: New Results. 10.1101/2025.07.21.665991.

83. Shumailov, I., Shumaylov, Z., Zhao, Y., Papernot, N., Anderson, R., and Gal, Y. (2024). AI models collapse when trained on recursively generated data. en. Nature 631. Publisher: Nature Publishing Group, 755–759. 10.1038/s41586-024-07566-y.

84. Wulsin, D., Gruver, N., Ji, C. X., Pruegsanusak, K., Scarpellini, G., Sharma, A., Swiderski, W., Bootsma, A. N., Bowen, R. S., Chen, C., et al. (n.d.). Pearl: A Foundation Model for Placing Every Atom in the Right Location. en ().

85. Varadi, M., Anyango, S., Deshpande, M., Nair, S., Natassia, C., Yordanova, G., Yuan, D., Stroe, O., Wood, G., Laydon, A., et al. (2022). AlphaFold Protein Structure Database: massively expanding the structural coverage of protein-sequence space with high-accuracy models. Nucleic Acids Research 50, D439–D444. 10.1093/nar/gkab1061.

86. Steinegger, M. and Söding, J. (2017). MMseqs2 enables sensitive protein sequence searching for the analysis of massive data sets. en. Nat Biotechnol 35. Publisher: Nature Publishing Group, 1026–1028. 10.1038/nbt.3988.

87. Suzek, B. E., Huang, H., McGarvey, P., Mazumder, R., and Wu, C. H. (2007). UniRef: comprehensive and non-redundant UniProt reference clusters. eng. Bioinformatics 23, 1282–1288. 10.1093/bioinformatics/btm098.

88. Fruchterman, T. M. J. and Reingold, E. M. (1991). Graph drawing by force-directed placement. en. Software: Practice and Experience 21. _eprint: https://onlinelibrary.wiley.com/doi/pdf/10.1002/s1164. 10.1002/spe.4380211102.

89. Blondel, V. D., Guillaume, J.-L., Lambiotte, R., and Lefebvre, E. (2008). Fast unfolding of communities in large networks. en. J. Stat. Mech. 2008, P10008. 10.1088/1742-5468/2008/10/P10008.

90. Katoh, K., Misawa, K., Kuma, K.-i., and Miyata, T. (2002). MAFFT: a novel method for rapid multiple sequence alignment based on fast Fourier transform. Nucleic Acids Res 30, 3059–3066. 10.1093/nar/gkf436.

91. Benjamini, Y. and Yekutieli, D. (2001). The control of the false discovery rate in multiple testing under dependency. The Annals of Statistics 29. Publisher: Institute of Mathematical Statistics, 1165–1188. 10.1214/aos/1013699998.

92. Rivoire, O., Reynolds, K. A., and Ranganathan, R. (2016). Evolution-Based Functional Decomposition of Proteins. en. PLOS Computational Biology 12. Publisher: Public Library of Science, e1004817. 10.1371/journal.pcbi.1004817.

93. Deorowicz, S., Debudaj-Grabysz, A., and Gudyś, A. (2016). FAMSA: Fast and accurate multiple sequence alignment of huge protein families. en. Sci Rep 6. Publisher: Nature Publishing Group, 33964. 10.1038/srep33964.

94. Edgar, R. C. (2022). Muscle5: High-accuracy alignment ensembles enable unbiased assessments of sequence homology and phylogeny. en. Nat Commun 13. Publisher: Nature Publishing Group, 6968. 10.1038/s41467-022-34630-w.

95. Martinez, Z. A., Murray, R. M., and Thomson, M. W. (2023). TRILL: Orchestrating Modular Deep-Learning Workflows for Democratized, Scalable Protein Analysis and Engineering. en. Pages: 2023.10.24.563881 Section: New Results. 10.1101/2023.10.24.563881.

96. Perez-Riverol, Y., Bandla, C., Kundu, D. J., Kamatchinathan, S., Bai, J., Hewapathirana, S., John, N. S., Prakash, A., Walzer, M., Wang, S., et al. (2025). The PRIDE database at 20 years: 2025 update. Nucleic Acids Res 53, D543–D553. 10.1093/nar/gkae1011.

97. Batra-Safferling, R., Granzin, J., Mödder, S., Hoffmann, S., and Willbold, D. (2010). Structural studies of the phosphatidylinositol 3-kinase (PI3K) SH3 domain in complex with a peptide ligand: role of the anchor residue in ligand binding. en. Biological Chemistry 391. 10.1515/bc.2010.003.

